# Single-nucleus transcriptomics and phosphoproteomics identify misregulated sperm components in mice carrying a selfish supergene

**DOI:** 10.64898/2026.10.21.760645

**Authors:** Reka K. Kelemen, Anna K. Lindholm, Beatriz Vicoso

**Affiliations:** Institute of Science and Technology Austria, Am Campus 1, 3400 Klosterneuburg, Austria; Department of Evolutionary Biology and Environmental Studies, University of Zürich, Winterthurerstrasse 190, 8057 Zürich, Switzerland

**Keywords:** meiotic drive, single cell sequencing, sperm motility

## Abstract

The *t*-haplotype is a selfish variant of the mouse chromosome 17. It reaches 90-100% transmission rates in heterozygous (+/*t*) males through a poison-antidote system that disables sperm carrying the homologous (+) haplotype. While several poisons and one antidote have been identified, how these mediate drive, and what additional molecular players are involved, is not fully understood. Here, we characterized patterns of gene expression throughout spermatogenesis in +/*t* versus +/+ males using single nucleus RNA sequencing. Genes differentially expressed in early spermatogenesis, when poisons are assumed to be expressed, were enriched for key regulators of sperm motility. These included dyneins, which provide the motor function of sperm flagella, and protein phosphatase 1 (PP1) complex genes, which regulate protein activity through dephosphorylation. Phosphoproteomics of +/*t* versus +/+ testes revealed changes in phosphorylation levels of several spermatogenesis proteins, supporting a contribution of dysregulated phosphorylation to the biology of the *t*-haplotype. In particular, the PP1 component PPP1CC2 itself showed significantly decreased phosphorylation in +/*t* testes. Further, we found that dyneins located on both the *t*-haplotype and the corresponding standard version of chromosome 17 have undergone large increases in spermatocyte expression in the house mouse lineage. We argue that this may result from an ongoing arms race between the homologs for transmission rates. Together, these results extend our understanding of the molecular changes associated with the *t*-haplotype, and highlight the value of integrating various OMICS approaches to investigate the mechanisms and evolutionary consequences of transmission distortion.

## Introduction

Meiosis and sexual reproduction are ancient processes that have highly conserved features across eukaryotes [Smith and Maynard-Smith, 1978], and whose purpose is the faithful transmission of genetic material to the next generation. However, transmission can be distorted by selfish variants (known as meiotic drivers) that are transmitted to more than 50% of the progeny [Sandler and Novitski, 1957, Henikoff et al., 2001, Presgraves et al., 2026]. Numerous meiotic drivers have been discovered in fungi, plants and animals [Lindholm et al., 2016]. In females, meiosis is asymmetric, as only one of the four products of meiosis becomes the oocyte (while the others yield polar bodies that do not get transmitted to the next generation). Female meiotic drivers usually alter the process of spindle attachment to increase their chances of segregating to the future egg [Clark and Akera, 2021, Malik and Bayes, 2006]. In contrast, meiotic drivers in males and fungi typically interfere with the development or function of gametes that do not carry them using two components: a distorter locus encoding a “poison”, or “killer”, that is cytoplasmically shared among all gametes, and a responder locus encoding an “antidote” or “insensitive target” that rescues only the gametes which carry the meiotic driver [Kruger and Mueller, 2021]. The promotion and evasion of cytoplasmic sharing of these two components are typically facilitated by the timing of expression to the early (diploid or early haploid) and late stages (haploid or late haploid) of gametogenesis, respectively [Dalstra et al., 2005, Nuckolls et al., 2022, Mange, 1968, Helleu et al., 2016, Charron et al., 2019].

These two-component systems need to be transmitted as a single genetic unit, something which is often achieved by the presence of inversions that suppress recombination with the standard non-driving chromosome [Dyer et al., 2007]. The transmission advantage of meiotic drivers creates a selective pressure for suppressors of drive to evolve on non-driving homologous chromosomes (and in the rest of the genome), and suppressors have indeed been found for many known meiotic drivers [Kruger et al., 2019, Clark, 1987, Nam et al., 2015, Helleu et al., 2016, Dawe et al., 2018]. Cycles of suppressor evolution followed by the evolution of additional enhancers of drive on the selfish chromosome can lead to long lived arms races between the chromosomes [Hall, 2004, Courret et al., 2019b, Courret et al., 2023]. Such arms races are thought to cause the high copy number of drive-related genes found on some driving chromosomes and their homologous regions [Dawe, 2022, Muirhead and Presgraves, 2021, Aravin et al., 2004]. Despite the understanding of these general principles of drive, the specific molecular mechanisms underlying transmission bias are in most cases not known (or only partly understood) [Courret et al., 2019a].

The *t*-haplotype of house mice is a selfish variant of the proximal end of chromosome 17, a region called the *t* complex. The *t*-haplotype is transmitted to more than 90% of the offspring of heterozygous males, and it was the first mammalian meiotic driver to be discovered [Dobrovolskaia-Zavadskaia and Kobozieff, 1927, Lyon, 2003]. It comprises one small and four large non-overlapping inversions that span approximately 40 megabases, each contributing to drive [Lyon, 1984, Swanepoel et al., 2025]. These inversions are estimated to have arisen approximately three million years ago [Morita et al., 1992, Hammer et al., 1989]. *t*-haplotypes cause either embryonic lethality or male sterility in a homozygous state, which prevents their fixation, but the underlying causes of these phenotypes are not fully understood [Klein et al., 1984, Lyon, 1991]. Although currently only found in the *Mus musculus* lineage, similar patterns of sequence orientation and gene amplification in *t*-haplotype inversions and homologous chromosome 17 regions in other *Mus* species support the hypothesis that *t*-haplotypes first arose in other *Mus* lineages or their common ancestors [Swanepoel et al., 2025]. Despite this long co-evolution of the *t*-haplotype with the mouse genome, no suppressors have been identified, although the fact that mice with different genetic backgrounds vary in their susceptibility to drive suggests that they may exist [Gummere et al., 1986].

Males that carry the *t*-haplotype produce + and *t*-sperm in equal amounts, but over 90% of their offspring inherit the *t*-haplotype [Silver and Olds-Clarke, 1984]. While the exact cause is unclear, sperm from +/*t* males show less progressive motility, premature hyperactivation, abnormal morphology and impaired penetration of the egg [Johnson et al., 1995, Olds-Clarke, 1989, Winkler and Lindholm, 2022, Amaral and Herrmann, 2021], some of which has been shown to affect + sperm to a larger extent [Amaral and Herrmann, 2021]. Understanding the molecular mechanisms underlying these phenotypes has been a long-standing goal [Amaral and Herrmann, 2021]. The antidote, SMOK^TCR^, which encodes a *t*-haplotype-specific version of the sperm tail kinase SMOK, was the first gene to be identified [Herrmann et al., 1999, Véron et al., 2009]. SMOK^TCR^ is hypothesized to act as a dominant-negative version of SMOK, thereby rescuing sperm from the hyperactivation of this pathway that occurs in *t*-carrying mice [Charron et al., 2019, Herrmann et al., 1999]. *Smok*^*Tcr*^ is exclusively expressed in haploid sperm cells and is not shared through the cytoplasmic bridges that connect developing sperm cells, such that its protein localizes to the tails of only half of the sperm in +/*t* males [Véron et al., 2009]. Four driver genes, *Tagap, Fgd2, Nme3* and *Tiam2* were later identified, which all showed expression starting in the early, diploid stages of spermatogenesis, and differential expression in +/*t* versus +/+ testes [Bauer et al., 2005, Bauer et al., 2007, Bauer et al., 2012, Charron et al., 2019]. All four of them are *t*-haplotype variants of members/regulators of the Rho signaling cascade, a pathway that modulates the progressive motility of sperm through the regulation of actin polymerization [Hanna and El-Sibai, 2013, Salgado-Lucio et al., 2020]. Interestingly, *Nme3* and *Tiam2* were not differentially expressed in carriers of a *t*-haplotype from Switzerland [Lindholm et al., 2019], raising the possibility that different *t*-haplotypes express and/or use different poisons to various extents. While these poisons are thought to act upstream of SMOK/SMOK^*TCR*^, the exact nature of their interaction, whether other players on the *t*-haplotype are involved, and how these pathways connect to sperm motility modulation, remain to be understood. Sperm motility is regulated by the reversible phosphorylation of dyneins, the motor proteins of the flagellum, by kinases and phosphatases [Freitas et al., 2017, King, 2016, Ishikawa, 2017]. The involvement of an antidote with a kinase function suggests that *t*-haplotype-carrying sperm may be rescued through the modulation of phosphorylation of flagellar proteins.

Other lines of evidence suggest a role for the modulation of protein phosphorylation by the *t*-haplotype. In particular, we previously showed that *t*-carrying mice overexpress several *t*-linked genes that encode components of the protein phosphatase 1 (PP1) complex in the testis [Kelemen et al., 2022]. The PP1 complex is a key regulator of sperm motility that dephosphorylates target proteins in the sperm flagellum [Chakrabarti et al., 2007, Smith et al., 1996, Freitas et al., 2017, Goswami et al., 2019]. Two PP1 regulatory subunits are encoded in the *t* complex: PPP1R11 and multiple copies of the chromosome 16 gene PPP1R2 (*Ppp1r2-ps1, Ppp1r2-ps5* and *Ppp1r2-ps6*). While these *Ppp1r2* copies are annotated as “pseudogenes”, several have a conserved protein coding sequence and high expression, suggesting they are functional (similar to what has been described in primates [Korrodi-Gregório et al., 2013, Gill et al., 2023]). The *t*-haplotype accumulated adaptive substitutions and additional gene copies of *Ppp1r11* [Kelemen et al., 2022, Swanepoel et al., 2025], *and has strongly increased testis expression of Ppp1r2-ps6* [Kelemen et al., 2022]. The *t*-haplotype further gained a gene copy encoding the catalytic PP1 subunit PPP1CB from chromosome 5, and this new copy, *Ppp1cb*^*t*^, evolved high and specific testis expression [Kelemen et al., 2022]. When in spermatogenesis these genes are overexpressed, and whether their differential expression is associated with changes in sperm protein phosphorylation, remain open questions.

Here, we investigated differential expression along spermatogenesis in mice that carry the *t*-haplotype versus mice that do not, using single nucleus RNA sequencing (snRNA-seq). snRNA-seq further allowed us to compare haploid sperm cells carrying the *t*-haplotype to those carrying the standard chromosome 17 within *t*-carrying individuals, allowing us to investigate the “battle of the cells” at unprecedented resolution. Several key spermatogenesis components, including dyneins and PP1 complex genes, were enriched among the set of genes that were differentially expressed in early spermatogenesis. To test if protein phosphorylation is misregulated in *t*-carrying mice, we compared the phosphoproteomes of +/*t* vs +/+ testes, and identified several potential targets. Dyneins further showed evolutionary patterns consistent with an arms race between the *t*-haplotype and the standard chromosome, highlighting the potential contribution of drivers to genome evolution.

## Results

### Single nucleus RNA-seq reveals differentially expressed genes at the resolution of cell types and single cell genotypes

To investigate changes in gene expression during spermatogenesis associated with the presence of a *t*-haplotype, we quantified gene expression at the single nucleus resolution in testes of two mice heterozygous for a Swiss *t*-haplotype (+/*t*) and two control (+/+) mice (see Methods and Supplementary table 1). 7493 high-quality nuclei were selected (see Methods and Supplementary figures 1 and 2), and were clustered based on their gene expression patterns after dimensionality reduction (Figure 1b). We recovered four somatic and seven germ cell clusters (Figure 1ab), using the expression of previously established markers for mouse testis cell types [Lukassen et al., 2018, Green et al., 2018] (Supplementary figure 3). While somatic and pre-meiotic germ cells are diploid, post-meiotic developing germ cells, the haploid spermatids, inherit either the *t*-haplotype or the standard chromosome 17. We previously inferred a set of *t*-haplotype-specific genetic variants, which we introduced into the reference genome to mimic a *t*-haplotype sequence [Kelemen and Vicoso, 2018]. We mapped our current data to both the reference and this pseudo-*t*-haplotype, and estimated how much mapping there was to the *t*-haplotype sequence relative to the homologous region of chromosome 17. Somatic and pre-meiotic germ cells do not show a bimodal distribution of this measure, in agreement with these diploid cells expressing both haplotypes (Figure 1c). While most diploid cells show equal average expression of genes on the *t*-haplotype and the homologous region of chromosome 17, the meiotic spermatocytes overexpress standard chromosome 17 genes compared to *t*-haplotype genes, causing a drop in the so-called *t*-score measure (Figure 1c). This trend is driven by many genes (Supplementary figure 4), and its underlying cause is yet to be explained. Haploid spermatid nuclei, on the other hand, could be classified as carrying either the *t*-haplotype or the standard chromosome 17 based on the set of alleles they primarily express (see Methods and Supplementary figures 5 and 6). In line with an equal ratio of + and *t* sperm produced by +/*t* males [Silver and Olds-Clarke, 1984], we found no significant deviation from an equal representation of + and *t* spermatids in any of the haploid clusters (p>0.2, Binomial test).

**Figure 1.**
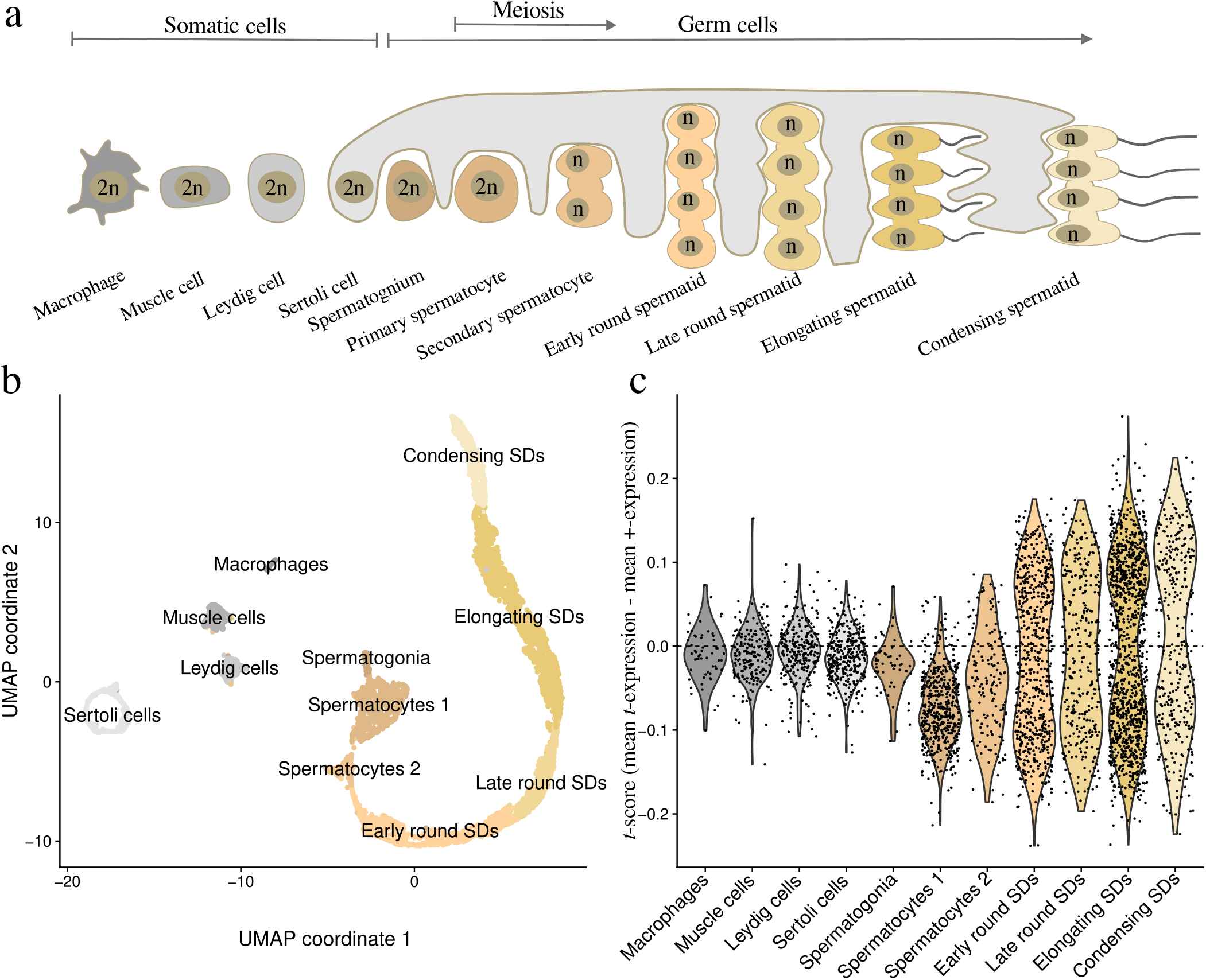
Nuclei cluster according to cell type and show *t*-haplotype- or +-haplotype-specific expression bias in haploid cell types. (a) Cartoon of the recovered somatic (shades of grey) and germ cell types (shades of beige), with Sertoli cells encompassing developing germ cells. Cell ploidy is indicated in the nuclei (2n for diploid and n for haploid nuclei). (b) The gene expression counts of the high quality nuclei were subject to dimensionality reduction and were plotted according to their UMAP coordinates (see Methods). Clusters were annotated using established *Mus musculus* testis celltype markers from [Lukassen et al., 2018, Green et al., 2018] (Supplementary figure 3). (c) Violin plots of the mean +-allele expression subtracted from the mean pseudo-*t*-allele expression for each nucleus. To increase the signal to noise ratio only those *t* complex genes were included, whose alleles were differentially expressed between the putative *t* and + spermatids (see Methods).

In order to fully represent the gene content of the *t*-haplotype and of the genetic background of the strain of the mice used in this study, we extracted reads from the haploid nuclei that carry the *t*-haplotype to assemble transcripts, which we then combined with transcripts assembled from a +/+ testis. We re-quantified gene expression using these combined transcripts in the high quality nuclei. We measured differential expression within each of the eleven cell type clusters that we identified, considering only genes that were expressed by at least 25% of the focal cells of either genotype. In case of diploid cells we compared +/*t* vs. +/+ nuclei, while in the case of haploid spermatid clusters we compared *t* spermatids and + spermatids of the +/*t* samples to each other, and to spermatids in the +/+ samples. We found the largest number of differentially expressed genes in meiotic spermatocytes and in haploid spermatids (Supplementary table 2). 20 of the 67 differentially expressed genes were encoded by the *t* complex, including ten overexpressed and ten underexpressed genes (Supplementary table 2). This is in line with previous observations that a substantial fraction of differentially expressed genes in *t*-carriers are encoded outside of the *t* complex and may represent downstream effects of factors encoded by the *t*-haplotype [Lindholm et al., 2019].

### Genes differentially expressed during early spermatogenesis are enriched for regulators of sperm motility

Previous work showed that *Smok*^*Tcr*^, the antidote of the *t*-haplotype, is expressed exclusively at the haploid stages of spermatogenesis, and remains close to the nucleus, which likely allows it to remain exclusively in *t*-haplotype carrying sperm [Véron et al., 2009]. Our data further refines the timing of *Smok*^*Tcr*^ expression to the latest haploid stages (elongating and condensing spermatids, Supplementary figure 7), in agreement with the idea that late expression may limit diffusion across spermatids. In contrast, poisons need to be shared across genotypes. Such sharing can be achieved either by expression in pre-meiotic or early haploid cells, where the cytoplasm of *t* and + haplotypes is directly shared or by exchange through the intercellular bridges that connect developing haploid spermatids. Therefore, we made the assumption that genes that showed their most significant germline differential expression (i.e. had the lowest adjusted p-value, see Supplementary table 2) in pre-meiotic spermatogonia, meiotic spermatocytes and in round spermatids were more likely to include poisons (“candidate poisons” in orange in Figure 2bc). On the other hand, genes that were most differentially expressed in elongating and condensing spermatids, similar to *Smok*^*Tcr*^, were under this assumption more likely to include additional antidotes (purple in Figure 2bc). It should be noted that, while consistent with the expression patterns of many known drivers, such expression-based classification may misclassify poisons that are expressed throughout spermatogenesis. Using a published measure of spermatid allelic bias [Bhutani et al., 2021], we find that genes classified as candidate poisons share a significantly higher fraction of their transcripts among haploid spermatids than genes primarily differentially expressed in later stages of spermatogenesis (p=0.022, Wilcoxon rank sum test, Figure 2c). Only two (*Nme3* and *Fgd2*) of the four previously identified poisons were detected in our dataset, and similar to the results of [Lindholm et al., 2019], they were not differentially expressed in any of the genotype and cell type comparisons (Supplementary Figure 7).

**Figure 2.**
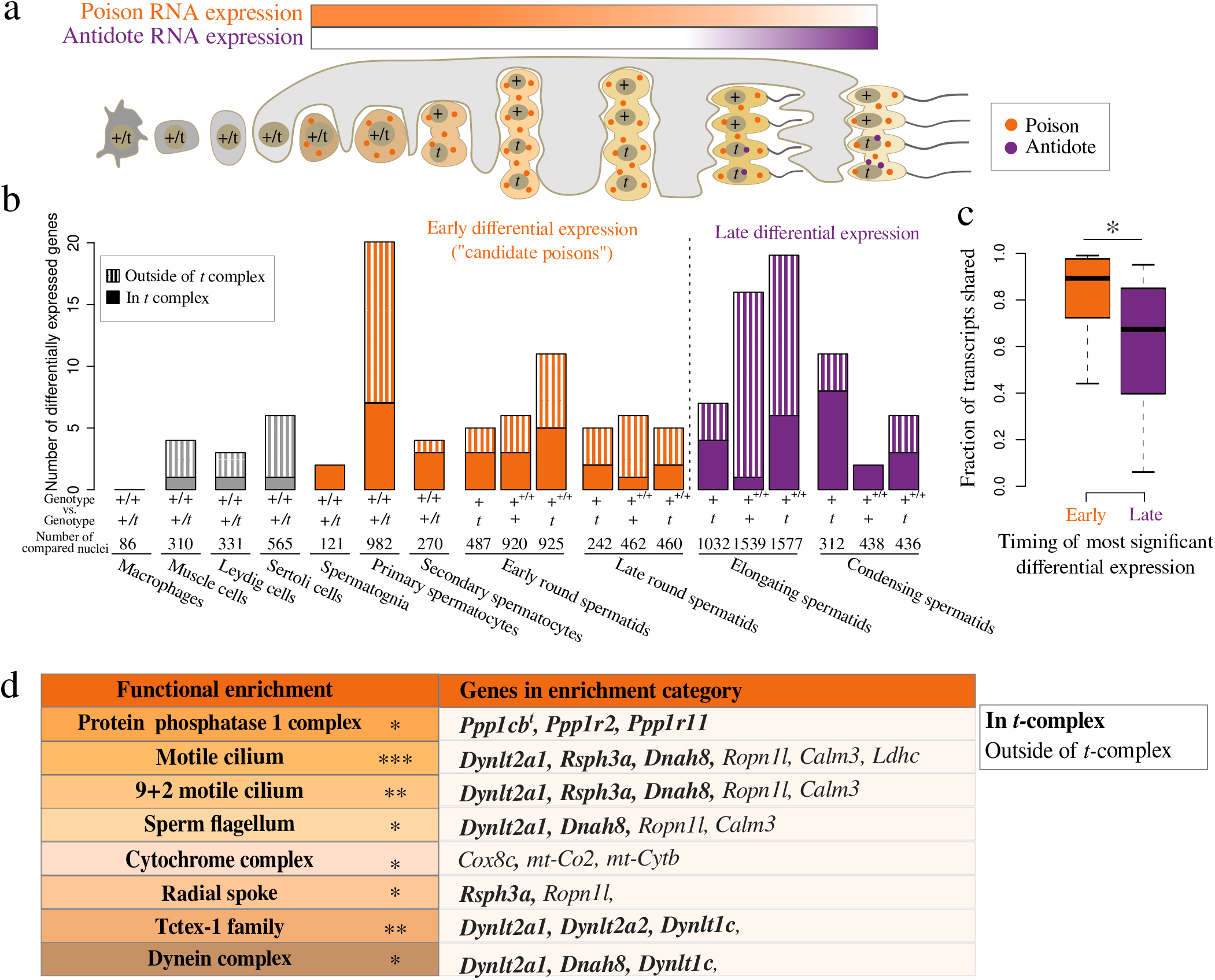
Differentially expressed genes throughout spermatogenesis. (a) Cartoon of testis cell types and the expected timing of poison and antidote expression during spermatogenesis, under the assumption that early expression facilitates the distribution of poisons to all gametes. (b) Numbers of differentially expressed genes in each cell type and genotype comparison. Expression in haploid spermatids was compared between + and t-genotypes within +/*t* testes and between both + and t-genotypes in +/*t* testes and + spermatids in +/+ testes. Solid bars show numbers of differentially expressed genes encoded in the *t* complex (5-40 Mb on chromosome 17), and striped bars show those encoded outside of the *t* complex. Numbers of total nuclei in each comparison are shown below the bars. (c) Boxplot of the estimated fraction of transcripts shared between haploid spermatids (extracted from [Bhutani et al., 2021]) for early- and late-differentially-expressed genes. (d) Functional categories detected with the tool StringDB, which show significant enrichment among candidate poisons (* FDR<0.05, ** FDR<0.01, *** FDR < 0.001), with bold (*t*-complex-encoded) and normal (non-*t*-complex-encoded) font gene names that fall into those categories.

To identify pathways that were modulated by the presence of a *t*-haplotype, we searched for gene ontology and gene pathway enrichments among our differentially expressed genes using StringDB. We found significantly more protein-protein interactions among the gene products that were differentially expressed in the germline than expected by chance (FDR=1.37e-07), and several functional categories were significantly enriched among candidate poisons (Figure 2d). First, we recovered a significant enrichment of protein phosphatase 1 (PP1) complex genes (3 genes, FDR=0.0412). These genes are comprised of *Ppp1cb*^*t*^, which encodes the *t*-haplotype-specific catalytic subunit of PP1, and two genes encoding PP1 inhibitors, *Ppp1r11* and *Ppp1r2*, the former of which is encoded in the *t* complex. As a copy of *Ppp1r2* was previously found to be encoded and overexpressed by the *t*-haplotype [Kelemen et al., 2022], we investigated if the *Ppp1r2* transcript that we found to be overexpressed (Supplementary figure 8a) is encoded on the *t*-haplotype. Indeed, we found that one of the nucleotide differences between this *Ppp1r2* transcript and the reference *Ppp1r2* transcript shows perfect and exclusive association with *t*-haplotype-carrier mice sampled across Eurasia (Supplementary figure 8b) [Harr et al., 2016]. These three PP1 genes were most differentially expressed in spermatocytes and round spermatids (adjusted pvalues < 0.05, Supplementary table 2). Dyneins, which, among other functions, act as the force-generating motor proteins of the sperm flagellum, were also overrepresented among candidate poisons (FDR=0.0186 and 0.0010 for “dynein complex” and “Tctex-1 family”, respectively). Specifically, four *t*-complex genes, *Dynlt1c, Dynlt2a1, Dynlt2a2* and *Dnah8*, which show adaptive coding sequence substitutions and copy amplification on the *t*-haplotype [Lyon, 2003, Pilder, 2012, Kelemen and Vicoso, 2018, Swanepoel et al., 2025] (Figure 4d-f), were differentially expressed in spermatogonia, spermatocytes and round spermatids (adjusted p-values < 0.0001, Supplementary table 2). Proteins that form the radial spokes, essential flagellar structures that coordinate sperm motility [Meng et al., 2024], were also enriched among candidate poisons (FDR=0.0472). These included the *t*-complex-encoded *Rsph3a*, which showed differential expression in spermatocytes and early round spermatids (adjusted p-values 1.35e-22 and 0.027, Supplementary table 2), and the non-*t*-complex-encoded gene *Ropn1l*, which was differentially expressed in spermatocytes (adjusted p-value 0.0497, Supplementary table 2). Other enriched functional categories were motile cilium (FDR=0.00097), 9+2 motile cilium (FDR=0.0056) and sperm flagellum (FDR=0.0371), which included some of the genes in the categories described above (see Figure 2d), and two additional non-*t*-complex-encoded genes, *Calm3* and *Ldhc. Ldhc* codes for the sperm-specific lactate dehydrogenase, which is an essential energy provider for sperm motility [Hereng et al., 2011, Odet et al., 2013]. *Calm3* codes for a calmodulin, and is involved in regulating sperm motility in response to calcium [Schlingmann et al., 2007, Freitas et al., 2017]. No copy of these genes or of *Ropn1l* have been found on the *t*-haplotype, such that they likely represent downstream effects of the *t*-haplotype.

Finally, the mitochondrial cytochrome complex was enriched among the candidate poisons (FDR=0.0227), potentially suggesting misregulation of mitochondrial number or activity. Specifically *mt-Co2, mt-Cytb* and *Cox8c* were differentially expressed in spermatogonia, spermatocytes and round spermatids (adjusted p-values < 0.05, Supplementary table 2). However, we found no increase in mitochondrial genomic copy number or expression level in bulk testis or epididymal sperm samples of +/*t* vs. +/+ males using quantitative PCR (Supplementary figures 9-11). The abundance of mitochondrial expression is known to be associated with cell quality in single cell RNA-seq experiments [Ilicic et al., 2016], such that this may represent an artefact rather than a true change in transcriptional rates.

The previously identified antidote, *Smok*^*Tcr*^, was significantly differentially expressed at the latest stage of spermatogenesis in our dataset (adjusted p-value = 9.55e-03, Supplementary Table 2). The rest of the genes that showed differential expression at the last two stages of spermatogenesis had no significant functional enrichments, but 5/35 were interaction partners of the candidate poisons (Supplementary figure 12). Classical genetics support the *Smok*^*Tcr*^ locus being the only responder locus on the *t*-haplotype [Lyon, 2003], such that any additional antidotes would have to be located there. Since mapping of these late-expressed genes to a published *t*-haplotype assembly [Swanepoel et al., 2025] shows that they are not clustered in this region (Supplementary figure 13), their late differential expression is more likely to be either unrelated to drive or a downstream effect of the *t*-haplotype, rather than evidence for additional antidotes.

### Proteomics of +/*t* and +/+ testes highlight differences in PPP1CB^*t*^ abundance and PPP1CC2 phosphorylation levels

Differentially expressed genes between +/*t* and +/+ mice during early spermatogenesis are enriched for PP1 complex genes. Since both PP1 [Goswami et al., 2019, V Silva et al., 2014, Freitas et al., 2017] and SMOK^TCR^, the known antidote of the *t*-haplotype [Herrmann et al., 1999], likely regulate the phosphorylation levels of proteins in the sperm tail, we hypothesized that the presence of the *t*-haplotype should be associated with differential regulation of protein phosphorylation. To test this hypothesis, and to identify potential targets, we used mass spectrometry on five +/*t* and five +/+ samples to quantify both protein abundance and, after experimental phospho-peptide enrichment, the abundance of phosphopeptides. We quantified the abundance of 120,681 peptides belonging to 9,818 proteins and 3566 phosphopeptides belonging to 2065 proteins in five +/*t* and five +/+ testes using mass spectrometry. To be able to quantify protein expression from the *t*-haplotype we predicted protein sequences from the published *t*-haplotype assembly [Swanepoel et al., 2025] and added these to the reference mouse proteome (see Methods). We first identified differentially abundant proteins between the two genotypes. We found that 11 of the 26 proteins for which we detected a significant difference in abundance between +/*t* and +/+ samples (adjusted p-value < 0.05) were encoded by the *t* complex. Our results show that the *t*-haplotype alleles of the PP1 complex proteins PPP1CB^*t*^ and PPP1R2^*t*^, are strongly overexpressed at the protein level in +/*t* vs. +/+ testes (Figure 3a), similarly to their RNA expression patterns. However, it should be noted that the protein abundance of PPP1R2 on the standard chromosome 17 was not quantified due to its annotation as a pseudogene and consequent absence from protein databases.

**Figure 3.**
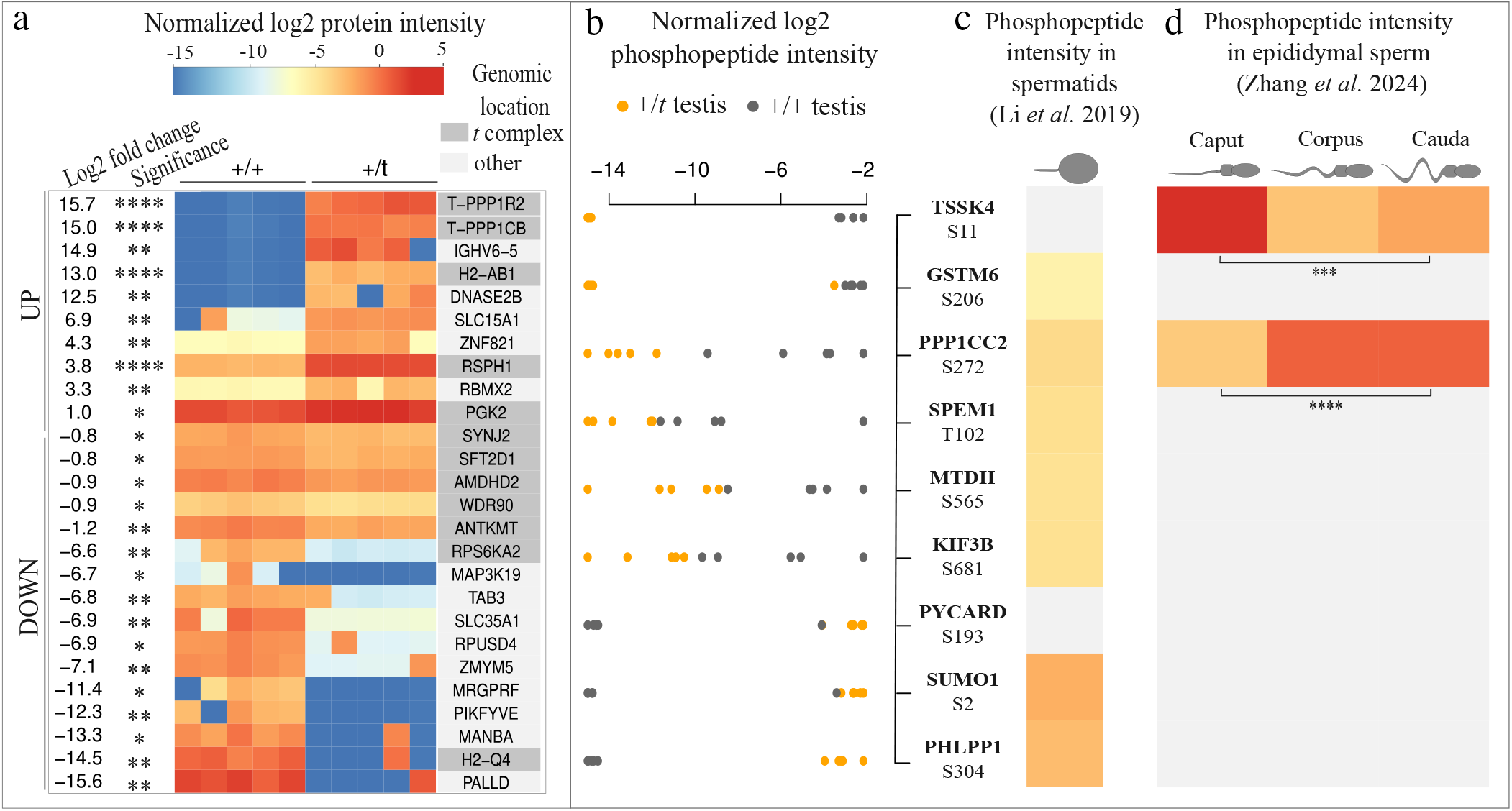
Differentially abundant and phosphorylated proteins in +/*t* vs. +/+ testes include PP1 complex proteins. (a) Heatmap showing the median-centered log2 aggregated intensities of proteins in five +/*t* and five +/+ testes where the effect of genotype is significant in a linear model fitted to the data (adjusted p-value < 0.05). The average log2 fold change between +/*t* vs. +/+ testes and the adjusted p-value levels (* p<0.05, ** p<0.01, *** p < 0.001) are indicated on the left side. Proteins encoded by the *t* complex (5-40Mb on chromosome 17) have their names on a dark grey background. UP and DOWN on the left side indicate proteins that are more or less abundant in +/*t* vs. +/+ testes, respectively. (b) The log2 intensities of phosphopeptides (normalized for varying parent protein abundances by subtracting the log2 intensities of their corresponding proteins within each replicate, see Methods), for which there was no overlap of intensity values between different genotypes and we observed a significant genotype effect after fitting a linear model to their intensities (p-value < 0.05). Orange and grey dots represent +/*t* and +/+ samples, respectively. Phosphopeptide names show the gene name in bold, and the amino acid and the position of the phosphorylated residue below. (c) The relative intensities of our differentially abundant phosphopeptides in spermatids as reported by [Li et al., 2019], color-coded by shades of orange (darker for higher values). Grey fields indicate that the phosphopeptide was not detected. (d) The relative intensities of our differentially abundant phosphopeptides in sperm from the caput, corpus and cauda epididymis as reported by [Zhang et al., 2024], representing the stages of motility acquisition, indicated by the sperm cartoons on top. Grey fields indicate that the phosphopeptide was not detected. Serine272 on PP1 and Serine11 on TSSK4 show significantly increased and decreased phosphorylation levels, respectively, in motile sperm of the cauda epididymis compared to the immotile sperm of caput epididymis [Zhang et al., 2024]

We then compared the abundances of phosphopeptides in +/*t* and +/+ testes after normalizing for their respective parent protein abundances within each replicate (see Methods). Because the sparsity of the data can yield false positives and negatives, we applied a multi-step filtering procedure to infer differential phosphorylation.

Phosphopeptides were inferred to be significantly different in their abundance between +/*t* and +/+ samples if 1. they were detected in all five samples of either genotype, 2. genotype groups did not overlap in their normalised intensity values for the phosphopeptide of interest and 3. the genotype effect was significant after fitting a linear model to the intensity values (p-value < 0.05, see Methods, Supplementary figure 14). We detected 11 phosphopeptides whose intensities were significantly different between +/*t* and +/+ testes (Figure 3b). Two of those were *t*-haplotype-specific peptides, and were excluded as they could not be compared between +/*t* and +/+ mice (Supplementary figure 15). Published phosphoproteomic data showed that 7 of the remaining 9 phosphopeptides are present in spermatids [Li et al., 2019] (Figure 3c). Several of these phosphopeptides affect proteins important for sperm development and function, such as the sperm maturation protein SPEM1, which is indispensable for male fertility [Zheng et al., 2007], SUMO1, which has an effect on sperm motility [Marchiani et al., 2014], TSSK4, a kinase influencing sperm tail morphology and motility [Wang et al., 2015] and the phosphatase PPP1CC2, which is necessary for sperm development and motility [Chakrabarti et al., 2007]. Of the differentially phosphorylated peptides only S272 of PPP1CC2 and S11 of TSSK4 are present in mature sperm [Zhang et al., 2024] (Figure 3d). PPP1CC2 is the testis-specific and sperm-enriched isoform of a non-*t*-complex-encoded PP1 protein, and this phosphorylation site is located on its unique C-terminal tail that is absent from other PP1 isoforms, including PPP1CB^*t*^ [Goswami et al., 2019]. The phosphorylation level of this PPP1CC2 phosphopeptide significantly increases, while the phosphorylation level of S11 on TSSK4 significantly decreases as sperm acquire motility in the epididymis in males (Figure 3d) [Zhang et al., 2024]. The significant underphosphorylation of both of these peptides in +/*t* testes makes them promising candidates for the sperm motility misregulation observed in +/*t* males.

### Expression evolution in *Mus musculus* spermatogenesis suggests an arms race of dyneins in the *t* complex

The *t* complex encodes three dynein gene families, which have multiple copies on both the standard chromosome 17 and on the *t*-haplotype (Figure 4d-f) [Swanepoel et al., 2025], and have elevated rates of coding sequence evolution in the *M. musculus* lineage compared to their homologs in other mouse species and rat (Figure 4d-f) [Kelemen et al., 2022]. Four of these dynein genes, which we found to be differentially expressed between the +/*t* and +/+ germlines, are among the most highly expressed *t*-complex-encoded genes in spermatocytes of both +/+ and +/*t* genotypes (Figure 4a). As gene copy amplification, fast coding sequence evolution and high expression are hallmarks of arms races between meiotic drivers and their homologs [Dawe et al., 2018, Muirhead and Presgraves, 2021, Aravin et al., 2004, Cocquet et al., 2009], we investigated the expression evolution of genes in the *Mus* genus. In particular, we tested whether the high spermatocyte expression of *t*-complex-encoded dyneins is specific to the *M. musculus* lineage, where the *t*-haplotype is found, as would be expected if they result from an arms race with the latter.

**Figure 4.**
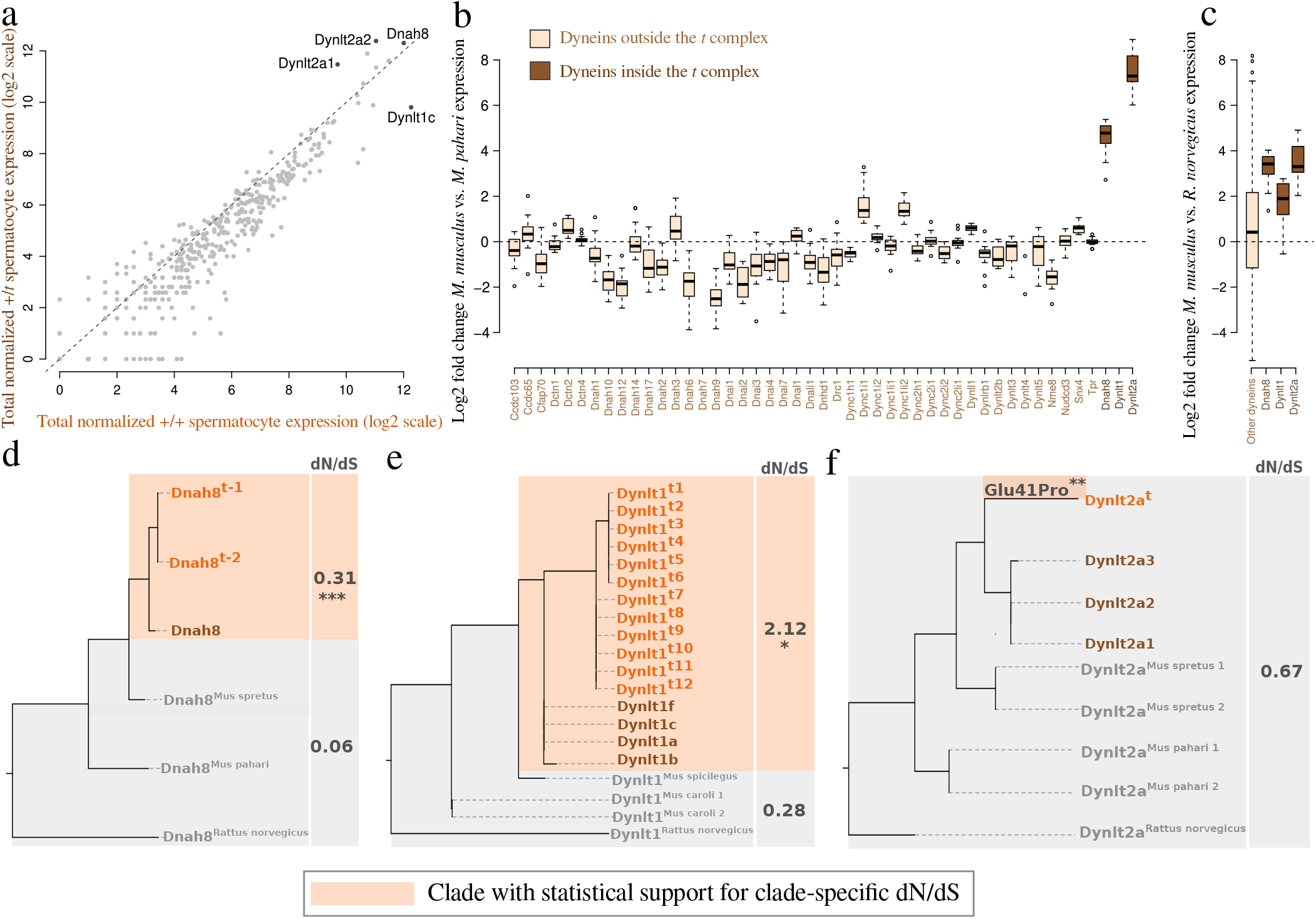
Dyneins are among the most highly expressed genes in the *t* complex, and have significantly increased in expression since the common ancestor of *Mus musculus* and *Mus pahari*. (a) Aggregated and cell-number-normalized log2 expression of *t*-complex-encoded genes in +/*t* vs. +/+ spermatocytes. Dyneins are indicated by their gene names next to their corresponding points. (b) Boxplots of log2 fold changes in spermatocyte expression levels of all *M. musculus* dynein complex and Tctex-1 family genes vs. the mean spermatocyte expression of their orthologs in the mouse *M. pahari*. Dynein complex genes and Tctex-1 family genes in the *t* complex are shown in brown, while those encoded outside the *t* complex are shown in beige. (c) Boxplots of log2 fold change in spermatocyte expression level of dynein complex and Tctex-1 family genes in *M. musculus* vs. the mean spermatocyte expression of their orthologs in the rat, *Rattus norvegicus*. The expression of dynein genes outside of the *t* complex were averaged for each sample and divided with the averaged dynein gene expression in rat samples. (d-f) Phylogenetic trees of the *M. musculus* (+ allele in brown, *t* allele in orange), *M. spretus, M. pahari* and *R. norvegicus* (all in grey) coding sequences of the three dynein gene families encoded by the *t* complex, as estimated by IQTree. The ratios of nonsynonymous to synonymous substitution rates (dN/dS), as estimated by PAML, represent the model that best fits the data. Clades or branches with a significantly increased dN/dS are shown on an orange background, and stars under the dN/dS values indicate how low the associated p-value is (* p<0.05, ** p<0.01, *** p < 0.001, Chi-square test of the difference in log likelihoods of the branch (for the *Dnah8* and *Dyntl1* genes) or branch-site model (for the *Dynlt2a* genes) to those of the null models of one shared dN/dS, or no positively selected sites, respectively). The amino acid substitution Glu41Pro on DYNLT2A^*t*^ was inferred to have been positively selected.

We used published expression data for spermatocytes of *M. musculus*, its sister species, *M. spretus*, and the more distant *M. pahari* [Kopania et al., 2022]. We also included spermatocyte expression estimates from the outgroup *Rattus norvegicus* [Chalmel et al., 2014]. It should be noted that no *t*-carriers were included in this analysis, such that the expression values of chromosome-17-encoded genes in house mice are represented only by the non-driving haplotype. We then performed comparisons of expression of orthologous genes for each pair of species; in case of gene copy number expansion within a species the expression of all ortholog copies was summed to allow for direct comparisons. We found that dyneins on the (non-driving) *t* complex have higher expression in *M. musculus* both relative to *M. pahari* and to *R. norvegicus* (Figure 4b and c). Phylogenetic modeling of gene expression evolution suggests that these increases in expression happened in the *M. musculus* lineage or the common ancestor of *M. musculus* and *M. spretus* (Supplementary figure 16). The evolutionarily recent and dramatic expression increase of *t*-complex-encoded dyneins is consistent with an arms race with their possibly poisonous homologs on the *t*-haplotype. Interestingly, the increase in dynein expression is to a lesser extent present in the close outgroup *M. spretus* (Supplementary figure 16). While the *t*-haplotype is not currently found in *M. spretus*, these high expression levels of *t*-complex-encoded dyneins in this species may provide some indirect support for the hypothesis that it was once present in this lineage [Swanepoel et al., 2025, Delarbre et al., 1988, Hammer et al., 1989].

We repeated this analysis of expression evolution for the other significantly enriched functional categories among our candidate poisons, namely for motile cilium, 9+2 motile cilium, sperm flagellum, radial spoke and PP1 phosphatase complex. While single genes, such as *Ppp1r11* (Supplementary figure 17) and *Rsph1* (Supplementary figure 18-26) stand out as overexpressed on the *M. musculus* lineage, we did not detect a systematic expression increase of *t*-complex-encoded genes.

## Discussion

Despite being the subject of research for a hundred years, the complex phenotype and molecular regulation of the *t*-haplotype have proven difficult to decipher [Lyon, 2003]. In this study we conducted the first genome-wide characterization of expression differences between +/*t* and +/+ males during spermatogenesis, providing new insights into what biological processes might be dysregulated in drive.

### Strengths and limitations of snRNA-seq and phosphoproteomics for the study of drive

Comparisons of testis transcriptomes of +/*t* vs +/+ males showed that many genes both on the *t* complex and elsewhere in the genome differ in their expression between the two, suggesting that more genes than the five known players may contribute to drive [Kelemen and Vicoso, 2018, Kelemen et al., 2022, Lindholm et al., 2019]. However, these bulk RNA-seq studies were unable to identify which changes in expression affected the germline, and could not distinguish between changes that affected all sperm versus those that differed between *t* and + sperm cells. [Swanepoel et al., 2025] compared spermatid bulk RNA-seq from *t*-carriers and controls, but did not assess expression at earlier diploid stages. By using single nucleus RNA sequencing, we obtained a cell-type- and cell-genotype-resolved picture of gene expression regulation. Similar to the discrepancy in single cell and bulk approaches that has been observed recently for a meiotic driver in stalk-eyed flies [Price et al., 2025], we identified many changes that had not been found with bulk RNA-analyses. Indeed, 46 of the 67 genes that we find differentially expressed in our celltype comparisons were not found at the testis level in [Lindholm et al., 2019], despite the fact that mice that carry the same *t*-haplotype variant and genetic background were used in both studies. However, some commonalities also emerge from the two approaches. First, our study confirms the observation of [Lindholm et al., 2019] that while *t*-complex-encoded genes are overrepresented among the differentially expressed genes, most of the differentially expressed genes are located elsewhere in the genome, suggesting downstream gene regulatory effects of the *t*-haplotype. We also recover many of the most differentially expressed genes of the bulk studies. In particular, the 12 genes that are differentially expressed in both our single nucleus study and [Lindholm et al., 2019]’s bulk testis study include the three PP1 complex genes (*Ppp1cb*^*t*^, *Ppp1r11* and *Ppp1r2*), the four dynein genes (*Dnah8, Dynlt1c, Dynlt2a1* and *Dynlt2a2*) and the radial spoke gene *Rsph3a*, highlighting a core set of genes with consistently modified expression in the presence of the *t*-haplotype. It should be noted that genes may contribute to drive through changes in their sequence without being differentially expressed, and such genes would not have been detected with our approach.

Several limitations of the data presented here should be noted. One main challenge when studying gene expression on the *t*-haplotype is the choice of reference genome or transcriptome to map reads against. Early studies used the mouse reference genome, which did not include *t*-specific sequences. [Kelemen et al., 2022] inferred *t*-specific sequences from RNA-seq data of +/*t* heterozygous mice for a few dozen genes, a small subset of the large number of genes found in the *t* complex. Recently, two genomic assemblies of the *t*-haplotype were published. However, one was based on a *t*-haplotype from the United States [Swanepoel et al., 2025]. The evolution of gene content is highly dynamic on *t*-haplotypes [Kelemen and Vicoso, 2018, Kelemen et al., 2022, Swanepoel et al., 2025], and using a diverged *t*-haplotype could limit our power to infer differential expression, or miss important variant-specific genes. The second is derived from the same Swiss *t*-haplotype used here [Runge et al., 2024]. However, its annotated sequence is incomplete, with key genes such as *Ppp1r11* not being represented despite previous work showing its presence and overexpression [Kelemen et al., 2022, Lindholm et al., 2019]. We therefore decided to create our own reference transcriptome from the single nucleus RNA-seq data. By assembling transcripts from haploid nuclei that carry the *t*-haplotype, we captured the expression signal from *t*-haplotype-specific sequences. However, due to the sparsity of single nucleus data, genes with low levels of expression may be missing from the assembly. Furthermore, while single nucleus RNA-seq has the key advantage of showing the transcriptional output of each cell, it provides little information on what transcripts are shared across cells, an important parameter to distinguish poisons and antidotes. Combining single nucleus and single-cell RNA sequencing may in the future shed light on what is shared and not shared between haploid spermatids. Finally, our bulk testis phosphoproteomics could not distinguish between changes happening in *t*- and +-carrying sperm, and may have missed functionally relevant phosphopeptides that are overrepresented in one but underrepresented in the other. Despite these limitations, our study yielded several new insights into the biology of the *t*-haplotype, highlighting the promise of single cell technologies for the study of drive [Price et al., 2025]. Since our study is correlational in nature, further work will be needed to functionally test the role of our candidate genes, but we conjecture below how some of them may influence drive.

### The sperm motility regulating PP1 complex is highly modified in *t*-carriers

PP1 genes were found to be differentially expressed in *t*-carriers compared with controls [Kelemen and Vicoso, 2018, Lindholm et al., 2019, Swanepoel et al., 2025], but whether they play a role in drive, and if so whether as poisons or antidotes, is not known. Our study strengthens the case for a role of PP1-mediated regulation of protein phosphorylation, and suggests that PP1 could work at least partly as a poison. The *t*-haplotype-specific *Ppp1cb*^*t*^ is highly expressed throughout meiosis (Supplementary table 2), which likely results in its incorporation in all sperm of +/*t* males. PPP1CB^*t*^ has a highly altered amino acid sequence compared to other PP1 isoforms [Kelemen et al., 2022], arguing against the conservation of its ancestral function. We therefore hypothesize that PPP1CB^*t*^ may instead interfere with normal PP1 function in the sperm of +/*t* males. Given the importance of regulatory subunits to PP1 biology [Bollen et al., 2010], one plausible mechanism could be for PPP1CB^*t*^ to sequester regulatory subunits required for PPP1CC2 activity, but without being able to replace the function of the latter (as it lacks the PPP1CC2-specific tail that is needed for correct sperm function [Dudiki et al., 2019]). Furthermore, the *t*-variant of PPP1R11 has lower phosphorylation levels compared to the standard PPP1R11 [Cheng, 2008], which may make it an abnormally strong or constitutive inhibitor of PPP1CC2 [Goswami et al., 2019]. Similarly, *t*-haplotype-associated sperm motility defects have been mapped in part to *Ppp1r11* [Pilder et al., 2007], supporting a potential role in drive.

Our finding that the C-terminal phosphorylation of residue S272 of PPP1CC2, the sperm-specific isoform of PP1, is significantly reduced in +/*t* testes compared to +/+ testes (Figure 3b) suggests that post-translational regulation of PP1 function may itself be altered by the *t*-haplotype. PPP1CC2 is repressed as sperm acquire motility during sperm maturation in the epididymis [Smith et al., 1996, Goswami et al., 2019], and this is accompanied by an increased phosphorylation of S272 on PPP1CC2 [Zhang et al., 2024]. The decreased PPP1CC2 phosphorylation that we observed in *t*-carriers may therefore imply increased PPP1CC2 activity and repressed sperm motility in these mice. Whether such a decrease in PPP1CC2 phosphorylation reduces the swimming potential of + sperm, or instead counteracts the *t*-caused prematurely hyperactive motility in order to rescue *t* sperm, still needs to be established. While single sperm proteomics will be needed to decipher if the reduced phosphorylation of PPP1CC2 is associated with + or *t* sperm in +/*t* males, the fact that the sperm abnormalities of *t*-carriers include premature hyperactivation [Olds-Clarke and Johnson, 1993, Amaral and Herrmann, 2021, Winkler and Lindholm, 2022] appears at least more consistent with the latter.

### Dyneins have highly dynamic evolution but are not differentially phosphorylated

Flagellar dyneins are known to be regulated through protein phosphorylation, and given the predicted role of both dyneins and and disrupted phosphorylation in drive, we might have expected these flagellar proteins to be differentially phosphorylated in *t*-carriers. Our phosphoproteomic study shows that both the + and *t*-alleles of DYNLT2A1 and RSPH3 (the only detected *t*-complex-encoded phosphoproteins) are targets of phosphorylation. However, they do not differ in their extent of phosphorylation at the testis level (Supplementary figure 19). It is possible that our bulk approach could not detect a sperm-specific differential phosphorylation of the t-alleles of these proteins, or that the changes in + and *t* sperm cancel each other out. Phosphoproteomic comparisons of sperm would be necessary to test if there is differential phosphorylation of *t*-haplotype-encoded dynein and radial spoke components. However, it is also possible that these proteins affect sperm function through their modified protein sequence and changes in dosage rather than by the rate at which they are phosphorylated. For example, the Glu41Pro amino acid substitution in DYNLT2A^t^ (Figure 4f) is a high impact amino acid substitution that affects a highly conserved region of the protein upstream of the dynein light chain domain (Supplementary Figure 27).

Alongside their putative role in drive, dyneins and radial spokes have unusual evolutionary patterns, and we interpret the latter as possible evidence of an arms race between the + and *t* alleles of these proteins. Evidence of positive selection was detected on the *t*-haplotype-allele of DYNLT2A and the radial spoke protein RSPH3 [Kelemen et al., 2022], and DYNLT1 and DNAH8 underwent gene copy amplification on the *t*-haplotype [Swanepoel et al., 2025]. This fast evolution of *t*-haplotype flagellar proteins was accompanied by equally dynamic evolutionary patterns of their homologues on the standard *t* complex (compared to genes with similar functions located elsewhere in the genome). In particular, there has been an increase in both copy number evolution of dyneins and radial spokes on the mouse chromosome 17, and in expression during spermatogenesis, since the split of the mouse and rat lineages (Figure 4, Supplementary figure 20). Interestingly, dyneins on the non-driving chromosome 17 show high copy number variation in house mouse populations, something that is not observed for other dynein complex genes (Supplementary figure 28). It is possible that the relative abundances of the + and *t* alleles of these dyneins influence the transmission ratios of the chromosome 17 homologs, similarly to what has been observed in selfish arms races in maize, fruit flies and the mouse sex chromosomes [Dawe, 2022, Muirhead and Presgraves, 2021, Kruger et al., 2019]. If this is the case, it could provide a potential explanation for the variable drive suppression observed in mice with different genetic backgrounds [Gummere et al., 1986]. Such long term arms races between the *t*-haplotype and the standard *t* complex may also provide an explanation for why homozygote *t*-carriers are sterile, despite expressing the antidote. If the standard *t*-complex has co-evolved to mitigate the effects of the *t*-haplotype, its absence in *t*-homozygotes may contribute to defective spermatogenesis, or, if it affects highly pleiotropic genes, to lethality. Similar single-nucleus studies on viable homozygote *t*-carriers may in the future help decipher which pathways are disrupted in these sterile mice, and whether candidate drivers and fast-evolving dyneins may also be involved.

### Extending the molecular model of *t*-haplotype drive

The current molecular model of *t*-haplotype drive is that four *t*-haplotype-encoded poisons (TAGAP, FGD2, NME3 and TIAM2) hyperactivate wild type SMOK proteins to abnormal levels in the sperm of +/*t* males, leading to the abnormally hyperactivated sperm motility observed in the case of +/*t* males. Under this model, the *t*-haplotype-encoded antidote, the hypomorphic SMOK^TCR^, acts as a dominant negative to counteract this hyperactivation exclusively in *t*-sperm to rescue normal sperm motility [Charron et al., 2019]. Some aspects of our data do not at first sight seem fully consistent with this model. First, despite the *t*-haplotype harboring *Smok* genes other than *Smok*^*Tcr*^ (two *Smok2*-like and four *Smok4*-like genes, see Supplementary figure 29) [Swanepoel et al., 2025], *t*-haplotype-carrying haploid spermatids show a significantly reduced *Smok4a* expression compared with non-*t*-haplotype-carrying spermatids (Supplementary figure 30a, Supplementary table 2). While *Smok2* genes were not assembled separately in our reference transcriptome, our initial expression analysis performed using the reference genome and pseudo-*t* sequence shows that they are also strongly down-regulated in *t* sperm of +/*t* mice, similarly to *Smok4a* (Supplementary figure 30b). This suggests that *t*-haplotype-carrying sperm may contain mostly SMOK^TCR^ proteins, arguing against a primary dominant negative role of SMOK^TCR^ towards other SMOK proteins. Second, while Herrmann and colleagues found differential expression of all four poison genes in their studies [Bauer et al., 2005, Bauer et al., 2007, Charron et al., 2019, Bauer et al., 2012] we find only very low expression of *Nme3* and *Fgd2*, which are also not differentially expressed between carrier and control mice, and no expression of *Tiam2* and *Tagap*. One possibility is that the Swiss *t*-haplotype studied here and the *t*-haplotype variants used by Herrmann and colleagues [Bauer et al., 2007, Bauer et al., 2012] use various poisons to different extents. In line with this, different *t*-haplotypes vary in their strength of transmission ratio distortion and sperm motility abnormalities even on the same mouse genetic background [Olds-Clarke and Johnson, 1993]. However, as noted above, our approach can miss genes with low expression, and it is possible that our study is simply uncovering a complementary aspect of *t*-haplotype biology.

Indeed, our results connect in several ways to the SMOK^TCR^ model (Figure 5c). The first, as we already noted, is the dysregulation of protein phosphorylation. Herrmann and colleagues [Charron et al., 2019] hypothesized that + sperm of *t*-carrying mice suffer from unbalanced protein phosphorylation due to the enhanced kinase function of hyperactivated SMOK proteins, contributing to their non-progressive motility. If the early-expressed and therefore likely shared PPP1CB^*t*^ acts as a dominant negative on PP1 function (see above), it may have a synergistic effect with SMOK by reducing protein dephosphorylation, further tilting the balance towards overphosphorylation of target proteins. In both cases, the rescue of *t*-haplotype-carrying sperm likely occurs by restoring the balance in phospho-rylation. The hypomorphic kinase activity of SMOK^TCR^ reduces phosphorylation rates [Herrmann et al., 1999]. An additional way of restoring balanced phosphorylation levels may be through increasing PPP1CC2 activity by reducing its phosphorylation levels.

**Figure 5.**
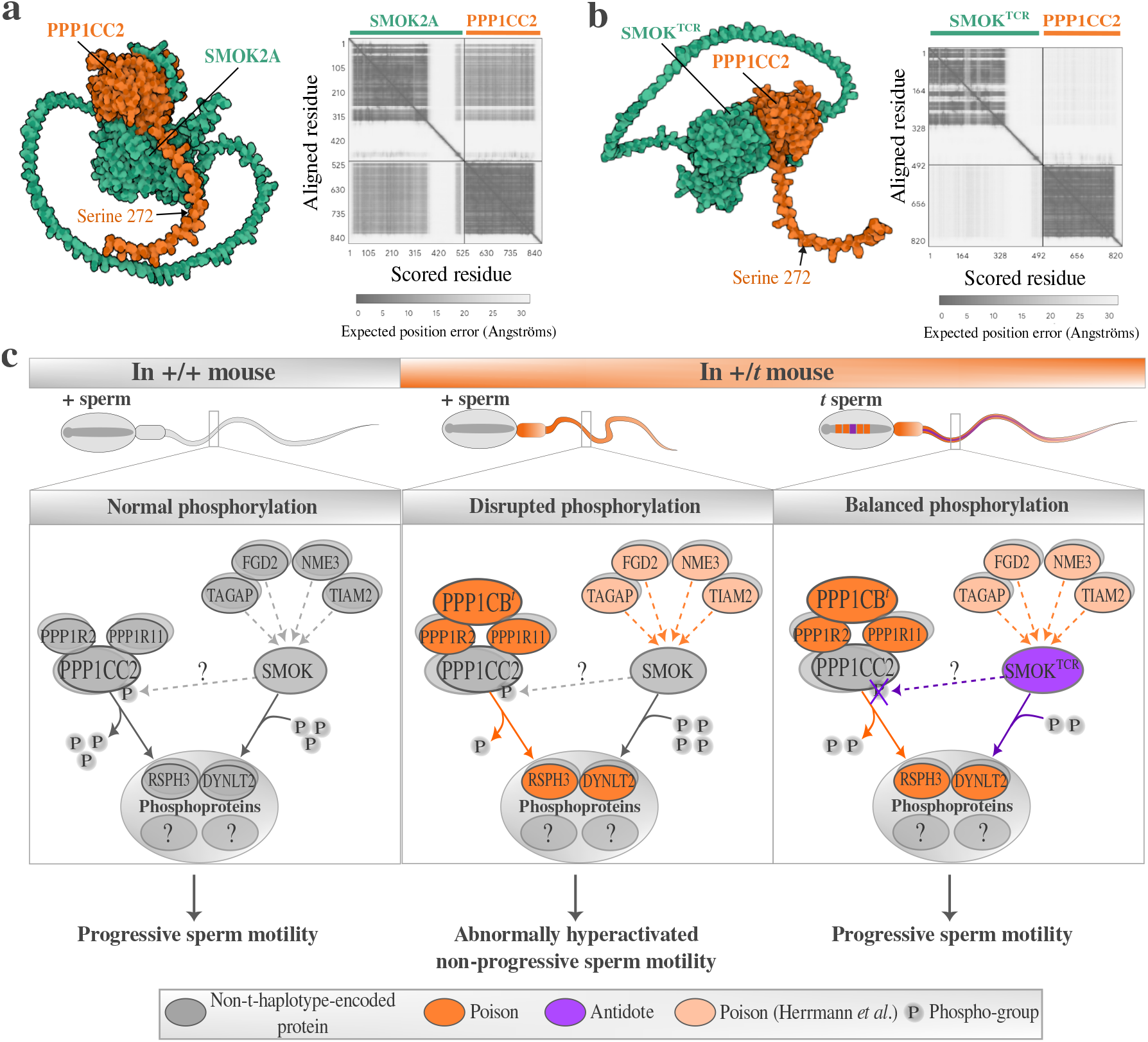
An extended hypothetical model of the molecular basis of *t*-haplotype drive. Three dimensional model of the binding of PPP1CC2 (orange) with (a) SMOK2A (green) and (b) SMOK^TCR^ (green), as predicted by Al-phaFold. The differentially phosphorylated S272 on PPP1CC2 is indicated by an arrow. The shading of the off-diagonal squares in the heatmap is inversely proportional to the expected position error of the amino acids. For the corresponding AlphaFold predictions for the interaction between PPP1CC2 and SMOK4A see Supplementary figure 32. (c) A molecular model of disrupted protein phosphorylation leading to non-progressive sperm motility and transmission ratio distortion by the *t*-haplotype. In the progressively moving sperm of +/+ mice, PP1 dephosphorylates, while SMOK phosphorylates target phosphoproteins, including *t*-complex-encoded dyneins and radial spokes (DYNLT2 and RSPH3 proteins). The wild type alleles of previously identified poisons [Charron et al., 2019] regulate SMOK function. In +/*t* mice all sperm receive the *t*-haplotype-encoded PPP1CB^*t*^, PPP1R11^*t*^ and PPP1R2^*t*^ (shown in dark orange), reduce the function of PPP1CC2. Synergistically, the poisons identified by Herrmann and colleagues (shown in light orange) hyperactivate the wild type SMOK proteins expressed by + sperm, further tilting the balance towards hyperphosphorylation. Disrupted phosphorylation leads to the abnormally hyperactivated non-progressive motility of + sperm in +/*t* males. In *t* sperm, PPP1CC2 function is increased by SMOK^TCR^, as a weaker phosphorylator of Serine 272 on PPP1CC2 than standard SMOK proteins. Further, SMOK^TCR^, a hypomorphic kinase, restores balanced phosphorylation of other target proteins, and rescues progressive motility of *t* sperm.

One further speculative but intriguing connection between the two pathways arose from an attempt to bioinformatically infer what kinase may be phosphorylating PPP1CC2 at S272, which predicted the likely role of a calcium/calmodulin-dependent protein kinase (CAMK) (prediction by tool Kinase Library, Supplementary figure 31). This prediction is interesting in light of previous findings that the sperm motility aberrations of sperm from +/*t* males become more severe with increasing calcium concentration in the medium [Olds-Clarke and Johnson, 1993, Olds-Clarke and Sego, 1992, Olds-Clarke and Wivell, 1992]. Further, while the targets of SMOK genes have not been identified, SMOKs are most similar to MARKs (MAP/microtubule affinity-regulating kinases), which belong to the CAMK kinase family, raising the possibility of a direct interaction between SMOK proteins and PPP1CC2. An AlphaFold2 analysis predicted binding between PPP1CC2 and SMOK2A (Figure 5b) (with a pTM value of 0.59, above the typical cut-off of 0.5 for similarity to a true structure) and SMOK2B (pTM = 0.5). On the other hand, the evidence is weaker for binding of PPP1CC2 with SMOK^TCR^ and SMOK4A (pTM = 0.43 in both cases, Supplementary figure 32). If SMOK2A phosphorylates PPP1CC2, thereby reducing PP1-mediated dephosphorylation, its replacement by SMOK^TCR^ in *t*-carrying sperm may contribute to the rescue of PP1 function and sperm motility. Experimentally identifying which proteins are phosphorylated by SMOK proteins will be crucial for understanding what the downstream targets of the driving mechanism are, and to uncover whether PPP1CC2 is indeed one of them.

## Materials and methods

### Single nucleus sequencing

We isolated nuclei from testis samples of four adult *M. m. domesticus* mice of the Illnau strain, which originated from Illnau, Switzerland, and were maintained in the lab for ten generations. Three of the mice were full siblings raised under standardized conditions: two were of +/*t*^*wILL*^ genotype, carrying a *t*-haplotype from Illnau, Switzerland (which harbors a recessive embryonic lethal gene [Lindholm et al., 2013]), and one of +/+ genotype. The other +/+ mouse was not a sibling of the other three mice. Mice we genotyped for being carriers of the *t*-haplotype by PCR amplification of the *t*-specific allele of *Hba-ps4*. Nuclei were isolated using the 10x Genomics nuclei isolation kit, and sequenced using Illumina Novaseq S4 PE150 XP.

### Finding high-quality nuclei

We created a pseudo-*t*-haplotype reference sequence by substituting *t*-specific SNPs into the *Mus musculus* reference genome (GRCm39). These SNPs were identified by Kelemen *et al*. [Kelemen and Vicoso, 2018] in a *t*-haplotype-carrying *M. m. domesticus* mouse from France, which was sampled by Harr *et al*. [Harr et al., 2016]. We appended this pseudo-*t*-haplotype sequence to the GRCm39 mouse reference genome. Further, we appended to these sequences the transcript sequences of two *t*-specific gained genes, *Ppp1cb*^*t*^ and *Rnpepl1*^*t*^, which were identified in [Kelemen et al., 2022], and also the sequence of the *t*-haplotype-specific antidote, *Smok*^*Tcr*^, identified by [Herrmann et al., 1999]. We annotated the pseudo-*t*-haplotype sequence by finding the coordinates of the best BLAT hits of the GRCm39 exons in the region of 5-40 Mb on chromosome 17. CellRanger (version 7.1.0) was used to map and count the UMIs (unique molecular identifiers) for each gene and barcode in a library. The fraction of intronic reads per nucleus was calculated with the nuclear_fraction_tag function of the DropletQC library in R (version 4.3.2) [Muskovic and Powell, 2021]. We used CellBender (version 0.3.0) with parameters expected-cells 7000, epochs 150, total-droplets-included 30000, fpr 0.05, learning-rate 0.00005, empty-drop-training-fraction 0.3, low-count-threshold 500, z-dim 50 and z-layers 200, to remove artefact barcodes as well as reads that likely represent ambient RNA. Using the Seurat package (version 5.0.1) [Butler et al., 2018] in R we filtered for nuclei with <5% mitochondrial UMI counts, an intronic read fraction >0.4 (Supplementary figure 1), and after [Germain et al., 2020], for nuclei that were within two median absolute deviations from the median of at least two of the following nuclei statistics: log10 of total UMI count, log10 of total gene count, ratio of log10 total gene count and log10 total UMI count, percent of counts coming from the top 20 most abundant genes (only the right tail was removed), percent mitochondrial UMIs, percent ribosomal UMIs. We used the SCTransform function to normalize each dataset by regressing on the percent mitochondrial and percent ribosomal UMIs and on the fraction of intronic reads, and identified the top 10,000 most variable genes. After executing the RunPCA and RunUMAP functions we removed barcodes that likely represent two nuclei with the DoubletFinder (version 2.0.4) package in R.

### Cell type identification

We merged all four filtered datasets and a dataset from a different mouse testis single nucleus RNA-seq study [Murat et al., 2023], which we used as an additional reference to flag artefact clusters. We normalized this merged dataset and found the 10,000 most variable genes with SCTransform, then we executed the RunPCA, FindNeighbors, FindCluster and RunUMAP functions with parameters dims=1:30 and resolution=2. We integrated these five datasets with the CCAIntegration method in the IntegrateLayers function, and executed the RunPCA, FindNeighbors, FindCluster and RunUMAP functions with parameters dims=1:20, resolution=1.2 and min.dist=0.1. For each cluster we assessed their expression of testis cell type marker genes, as identified by [Lukassen et al., 2018, Green et al., 2018], the fraction of contained cells in our vs. the Murat *et al*. dataset, the ratio of mean X chromosomal to autosomal expression and the distribution of intronic read fraction. We discarded clusters that were deviant in all of these aspects (Supplementary figure 2). We re-integrated the remaining nuclei without the Murat *et al*. dataset, ran the RunUMAP, FindNeighbors and FindClusters functions with parameters dim=1:12 and resolution=0.5, and annotated the clusters based on which cell type markers they most strongly expressed (Supplementary figure 3).

### Identification of haploid spermatids carrying the *t*- or the +-haplotypes

We computed a preliminary *t*-score for each nucleus by subtracting the mean + allele expression from that of the *t* allele expression of each expressed *t* complex gene. We fit two normal distributions to the resulting bimodal distributions of *t*-scores in the haploid cell clusters of the +/*t* samples using the *mix* function of the *mixdist* package (version 0.5-5) in R with parameters mixparam(mu=c(−0.02,0.02), sigma=c(0.01,0.01)), where *mu* parameters were estimated from the data using the *multimode* package (version 1.05) in R. Nuclei with *t*-scores above the 10th percentiles of the high-*t*-scores distributions were tested against nuclei below the 90th percentiles of the low-*t*-scores distributions for differential expression of *t* complex gene alleles using the MAST test of the FindMarkers function of Seurat. We used the 25th and 75th percentiles for the early and late round spermatid clusters, because of less bimodal *t*-scores (Supplementary figures 4 and 5). Genes with differentially expressed alleles and an FDR-adjusted p-value below 0.05 were used to compute a refined *t*-score using the same formula as above. We fit two normal distributions to the bimodal *t*-score distributions in each haploid cluster. Nuclei above the 10th percentile of the high-*t*-score distribution were classified as *t*-spermatids, and those below the 90th percentile of the low-*t*-score distribution were classified as +-spermatids. We used the 25th and the 75th percentiles, respectively, for the round spermatid clusters (Supplementary figure 5).

### Assembling transcriptomes of *t*-spermatids and +/+ testes

We extracted reads from the raw FASTQ files of the two +/*t* samples using umi_tools’ *extract* function in Python (version 3.9.7) and the list of our *t*-spermatid barcodes. Reads were trimmed with *fqtrim* (version 0.9.7) with parameters -5 AAGCAGTGGTATCAACGCAGAGTACATGGG (template switch oligo sequence) -a 10 -q 20 -l 50. Files containing read 2’s from both +/*t* samples were concatenated and assembled in a forward stranded mode (–SS_lib_type F) using Trinity and a minimum contig length of 500 (version 2.15.1). Reads from the +/+ library were trimmed with the above parameters and read 2’s were assembled using the same procedure as mentioned above. We filtered for contigs that showed a minimum TPM of 0.5, estimated using kallisto (version 0.50.1), in both samples of the focal genotype. Contigs with an overlap of at least 100 basepairs and less than 0.01 sequence differences, as inferred by BLAT [Kent, 2002], were collapsed by choosing the longest of such matching contigs using a custom-made Perl script. The *t*-spermatid assembly and the +/+ testis assembly were concatenated, and contigs with highly similar overlapping regions were collapsed, as before. We mapped each contig to the GRCm39 transcriptome and to the *t*-specific gained genes, *Ppp1cb*^*t*^ and *Smok*^*Tcr*^, and we assigned each to a gene if it had a minimum of 200 basepair alignment to one of its transcripts.

### Differential expression analysis

We quantified the expression of each contig using CellRanger. Then for each gene in our assembly and high-quality nucleus identified before we added up the UMI counts of the contigs mapping to it, and created four Seurat objects, which we merged and normalized with the SCTransform function and identified the 5,000 most variable genes. We then transferred the PCA, UMAP, cell type annotation, t-score and inferred cell genotype for each nucleus from the Seurat object that was based on mapping to the GRCm39 reference. We then used the built-in MAST [Finak et al., 2015] test of the FindMarkers function of Seurat, specifically suited for single cell datasets, to find differentially expressed genes between the two +/*t* and the two +/+ samples within each diploid cell cluster. For haploid cell clusters we compared *+*^+*/t*^ spermatids to *+*^+*/*+^ spermatids, *t*^+*/t*^ spermatids to *+*^+*/*+^ spermatids and *t*^+*/t*^ spermatids to *+*^+*/ t*^ spermatids. We considered only those genes to be differentially expressed, which were expressed in at least 25% of at least one of the compared cell populations, had a minimum log2 fold change in any direction of at least 0.1 and had an adjusted p-value below 0.05.

### Gene ontology enrichment analysis

We provided our list of differentially expressed genes to the online database and gene ontology search tool, STRING (https://string-db.org/,version12.0)[Szklarczyk et al., 2025] to identify enriched gene ontology categories. The default *Mus musculus* proteome was used as the background protein set to identify functionally enriched categories. Tissue and KEGG enrichments were not shown, as all our proteins were derived from as single tissue (testis), and KEGG enrichments were largely driven by the mitochondrial genes (which we believe are artefacts), but are provided in Supplementary table 3.

### Quantitative PCR of mitochondrial DNA and RNA in testes and epididymal sperm of +/*t* and +/+ males

We extracted DNA from testis samples of two +/*t* and two +/+ males from the Illnau strain, and DNA and RNA samples from epididymal sperm samples from two +/*t* and two +/+ males from the Illnau strain. For three technical replicates per sample we performed quantitative PCR measurements for primers designed for *Ppp1cb*^*t*^ and the mitochondrial genes *Mt-Co3, Mt-Rnr1* and *Mt-Cytb*. As a control, we designed primers for the nuclear single copy genes *Gtf2f2* and *Ppp1cb*. The log2 fold change in RNA or DNA compared to the single copy nuclear genes was calculated as the difference in the number of cycles for each primer and sample from the number of cycles of the control genes.

### Proteomics and phosphoproteomics

#### Sample preparation

Both testes of each of 5 +/*t* and 5 +/+ *M. m. domesticus* adult mice of the Illnau strain were frozen in liquid nitrogen. The testes were homogenized and cleaned up by SP3 using a commercial kit (PreOmics GmbH, 100 mg of beads per sample), then processed using the iST kit (PreOmics GmbH) using the manufacturer’s modified protocol for tissue samples. Tryptic digestion was stopped after overnight incubation and cleaned-up samples were vacuum dried. Finally, samples were re-dissolved with 10 min sonication in a 9:1 mix of the iST kit’s LC-LOAD buffer and 0.15% w/v N-Dodecyl-Beta-D-maltoside (DDM, 0.015% final).

#### Phosphoenrichment

Phosphopeptides were enriched starting from 20 µL input per pooled peptide sample using Zr-IMAC HP beads (ReSyn Biosciences) according to the manufacturer’s protocol. Phosphopeptide samples were re-dissolved with 10 min sonication in 0.15% N-Dodecyl-beta-D-maltoside (DDM) in 0.1% Formic Acid.

#### LC-MS/MS analysis

Samples were analyzed by LC-MS/MS on a nanoElute 2 nano-HPLC (Bruker Daltonics) coupled with a timsTOF HT (Bruker Daltonics), concentrated over an experimental Inertsil ODS-4 column (C18-coated particles, 3 cm), then bound to a monolithic capillary MOSAIC column (C18-coated particles, 15 cm * 75 µm ID, Bruker P/N 1908374) heated at 50°C and eluted over the following 60 min gradient: solvent A, MS-grade H O + 0.1% formic acid; solvent B, 100% acetonitrile + 0.1% formic acid; constant 0.30 nL/min flow; B percentage: 0 min, 5%; 35 min, 17%; 55 min, 35%; 60 min, 95%, followed immediately by a 5 min plateau at 95%. MS method: M/Z range = 99.993933-1700 Th, ion mobility range = 0.7-1.4 1/K0; transfer time = 60 µs, pre-pulse storage time = 12 µs, enable high sensitivity modus = off, ion polarity = Positive, scan mode = dia-PASEF; TIMS parameters: ramp time = 180 ms, accumulation time = 180 ms; PASEF parameters: ms/ms scans = 4, total cycle time = 0 s, charge range = 0-5, intensity threshold for scheduling = 1500, scheduling target intensity = 15000, exclusion release time = 0.4 min, reconsider precursor switch = on, current/previous intensity ratio = 4, exclusion window mass width = 0.015 m/z, exclusion window v·s/cm^2^ width = 0.015 V·s/cm^2^.

#### Data analysis

Raw files were searched in DiaNN version 2.2.0 [Demichev et al., 2020] against the spectral library generated from the *Mus musculus* complete proteome sourced from UniprotKB and the bioinformatically predicted proteome of the *t*-haplotype version *tw5* [Swanepoel et al., 2025]. Variable modifications were set to Acetyl (protein N-term) and Phospho (STY). The data was filtered at 1% FDR. Peptide intensity tables were analyzed with the msqrob2 package (version 1.18.0) [Vandenbulcke et al., 2025] in R (version 4.5.1). Peptides that were detected in at least 2 out of 10 samples were kept, log2-transformed and normalized by median centering to remove sample-specific variability. The *aggregateFeatures* function was used to summarize the peptide abundances to their respective proteins or protein groups, adding both *t* and + alleles for *t*-complex-encoded proteins. We estimated the genotype effect using the *msqrob* function tested for significance using the *makeContrasts* and *hypothesisTest* functions of the limma package (version 3.66.0) in R. Proteins with a Benjamini-Hochberg-adjusted p-value below 0.05 were considered to be significantly different in abundance.

For the phosphoenriched samples, we used the same DiaNN search and used phosphosites predicted with below 1% FDR. Using the PhosR package (version 1.20.0) [Kim et al., 2021] in R, we kept only those phosphopeptides that were detected in all five samples of either genotype group, log2-transformed and normalized the intensity values using the *medianScaling* function of PhosR, with scale=F. Then for each phosphopeptide and sample we subtracted the corresponding quantile-normalized log2 protein abundance value in order to account for variable parent protein abundances across samples. Samples that had neither phosphopeptide nor the corresponding protein detected were assigned the minimum value for that phosphopeptide minus three log2 units. We estimated the genotype effect for each phosphopeptide with the limma package in R, and kept those with a p-value below 0.05, and whose intensity values did not overlap between genotype groups.

### Expression evolution analysis

We used the expression estimates in spermatocytes from the study of [Kopania et al., 2022] sampled from 18 *Mus musculus* (10 *M. m. domesticues* and 8 *M. m. musculus*), 11 *M. spretus* and 3 *M. pahari* individuals. As an outgroup, we added spermatocyte expression from 2 *Rattus norvegicus* individuals from the study of [Chalmel et al., 2014]. The branch lengths of the phylogenetic tree comprising these species were based on the estimates of [Chevret et al., 2005] and [López-Antoñanzas et al., 2024]. The R package evemodel (version 0.0.0.9008)[Rohlfs and Nielsen, 2015] was used to estimate for each dynein the log likelihood of an expression shift on each of the branches of the *Mus* phylogeny, and the likelihood of no expression shift. For each of these models the AIC score was calculated using the formula 2*k − LogLikelihood*, where k is the number of parameters of the model (1 for the model with shared mean expression, and 2 for a model with lineage-specific shifted mean expression). A model was considered best fitting if its AIC score was lower than the next lowest AIC score by at least 2 AIC units. p-values were obtained by comparing the likelihood ratio test of the best-fitting model to the null model of shared expression mean using the Chi-square distribution with 2 degrees of freedom.

### Coding sequence evolution of dyneins in the *t* complex

We used the tool miniprot (version 0.15-r270) and the mouse proteome to predict dynein coding sequences from the *t*-haplotype sequence from [Swanepoel et al., 2025]. We aligned the coding sequences of dynein orthologs from the *t*-haplotype, the *M. musculus, M. pahari, M. spretus* and *R. norvegicus* reference genomes using the web tool MUSCLE [Madeira et al., 2024]. We estimated a phylogenetic tree using the software IQTree (version 2.1.3) [Minh et al., 2020]. We estimated the ratios of nonsynonymous to synonymous substitution rates using the tool PAML (version 4.10.10) [Yang, 2007], and used likelihood ratio tests to find the best fitting model for each phylogenetic tree.

## Acknowledgments

We thank Paul Thomas for comments on the manuscript, and Carrie Bernecky for guideance on PP1 biology. Further, we thank the Vicoso group for discussion of the work and the ISTA Scientific Computing and Proteomics facilities for their support.

## Data availability statement

All sequencing data generated for this study has been deposited on the NCBI short read archive under Bioproject number [will be provided before publication]. The single nucleus expression dataset in an R-compatible RDS format is provided under the link: https://seafile.ist.ac.at/d/cf2ac94a23ad462a8833/. The mass spectrometry proteomics data have been deposited to the ProteomeXchange Consortium via the PRIDE [Perez-Riverol et al., 2025] partner repository with the dataset identifier PXD084971. All code used in this study is provided on the Github repository https://github.com/reka-kelemen/t-haplotype-multiomics. Data from the following NCBI samples and Bioprojects was used: SAMN02214043 and SAMN02214046 for the *Rattus norvegicus* spermatocyte expression data and all spermatocyte RNA-sequencing samples from the bioproject PRJNA735780 for the *M. musculus, M. spretus, M. pahari* spermatocyte expression data. The list of copy number variant regions in wild-caught mice is available on the ftp site http://wwwuser.gwdg.de/evolbio/evolgen/wildmouse/.

## Notes

### Competing Interest Statement

The authors have declared no competing interest.

### Funding

- Austrian Science Fund (FWF): 10.55776/F88
- Swiss National Science Foundation: 310030_189145
- Australian Research Council: DP250101115

