## Supplementary figures and tables for "Single-nucleus transcriptomics and phosphoproteomics identify misregulated sperm components in mice carrying a selfish supergene"

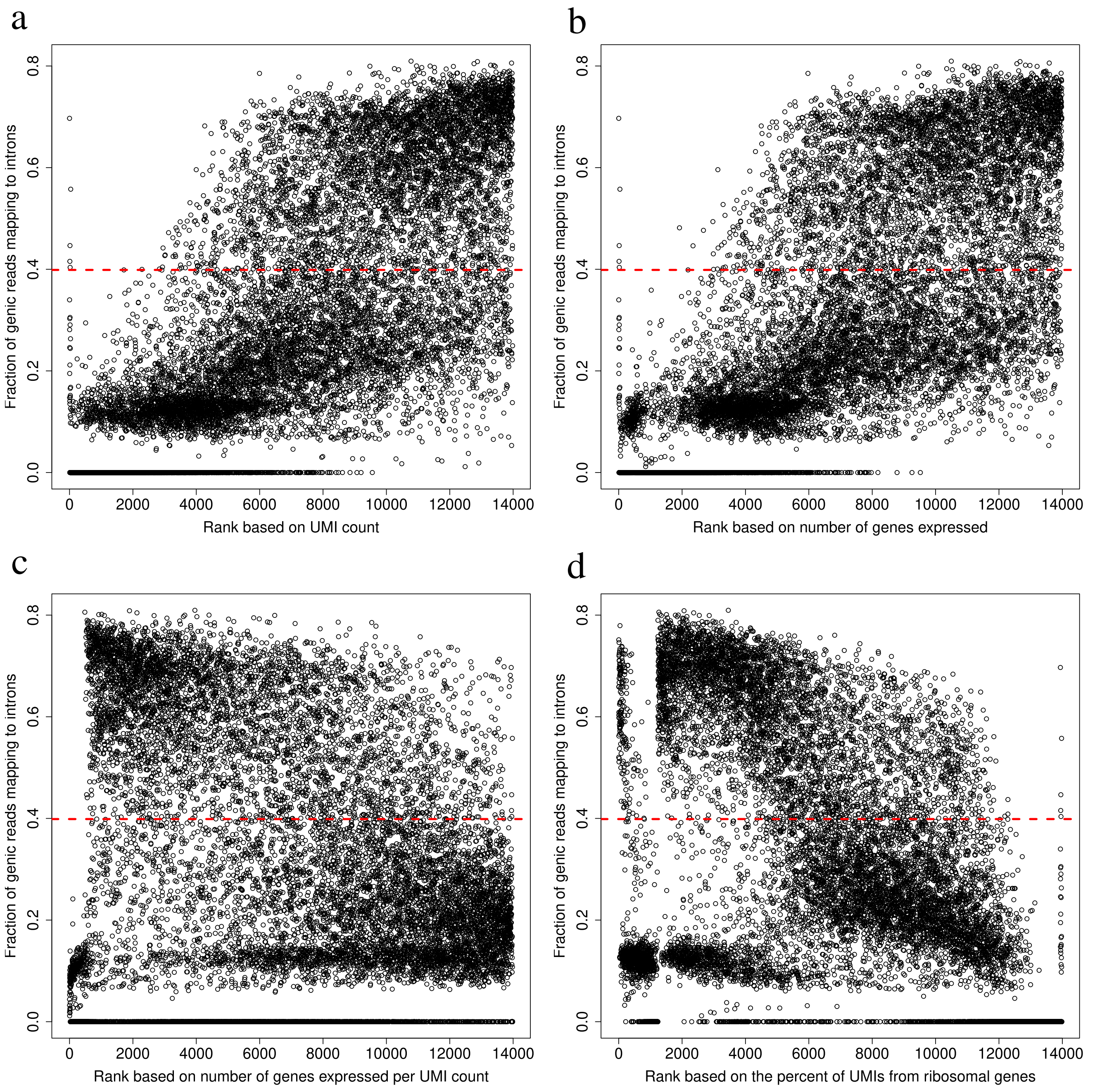


Supplementary Figure 1: The fraction of genic reads mapping to introns for each nucleus are plotted against the rank of each nucleus based on the (a) total UMI count, (b) number of genes expressed, (c) number of genes expressed per UMI count and (d) the percent of UMIs from ribosomal genes. The red dashed lines indicate the cutoff (0.4) of the fraction of genic reads mapping to introns, above which nuclei were considered for further analysis.

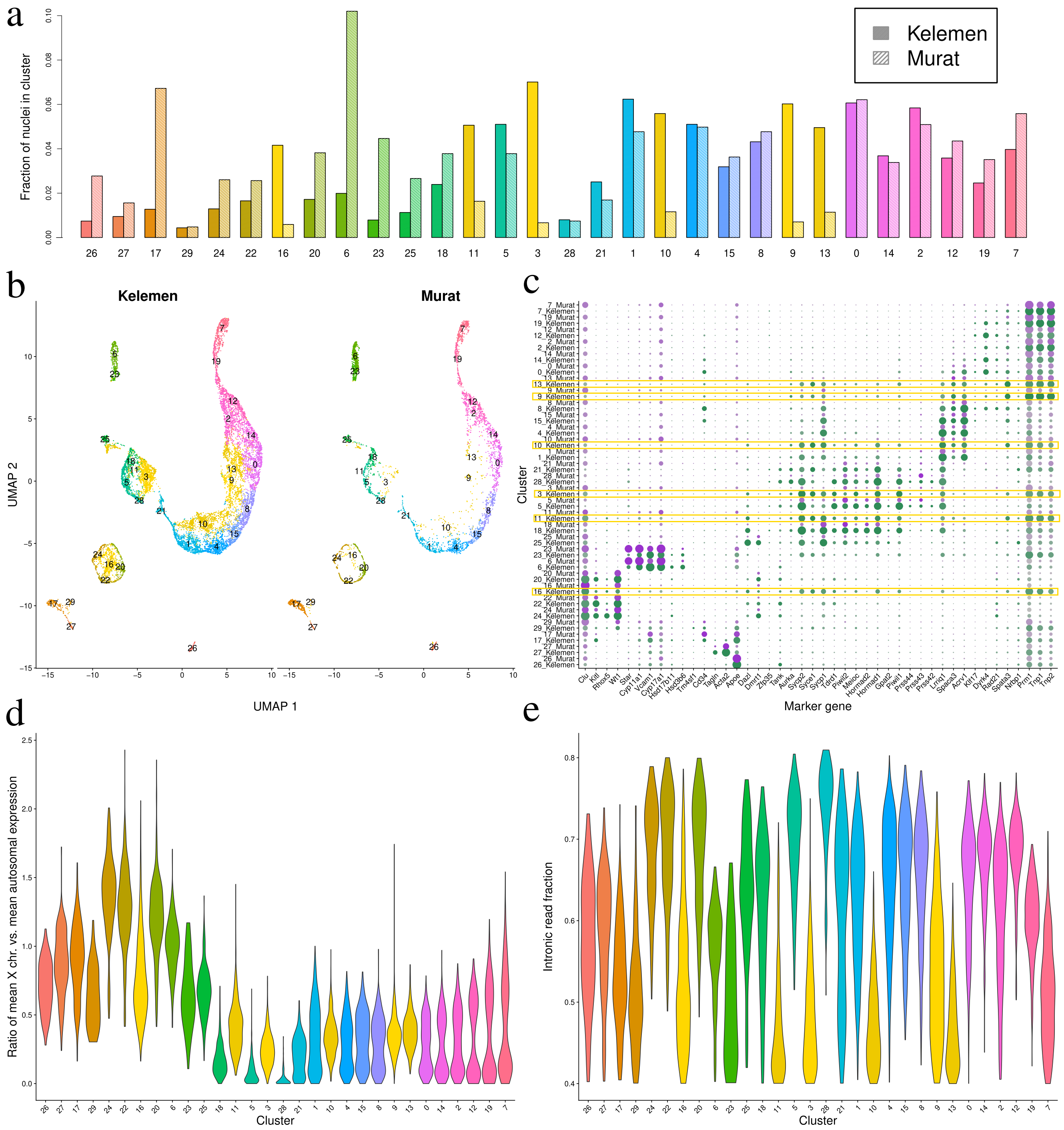

Supplementary Figure 2: Diagnostic measures for identifying artefact clusters (highlighted in yellow), which may contain whole cells or empty droplets with ambient RNA. (a) Fraction of nuclei in each dataset present in each cluster. Striped bars correspond to the dataset of [Murat et al., 2023], full bars represent the dataset of this study. (b) Nuclei from the current study Kelemen *et al.* (left panel) and the Murat *et al.* dataset (right panel) are shown after dimensionality reduction according to their first two UMAP coordinates, color coded according to their clusters. (c) The percent of nuclei expressing each of the cell type marker genes is shown by the size of the circles for each cluster in the Kelemen *et al.* (green) and the Murat *et al.* (purple) datasets. (d) Violin plots showing the distribution of the ratio of mean X chromosomal vs. mean autosomal expression for each cluster. (d) Violin plots showing the distribution of the intronic read fraction for each cluster.

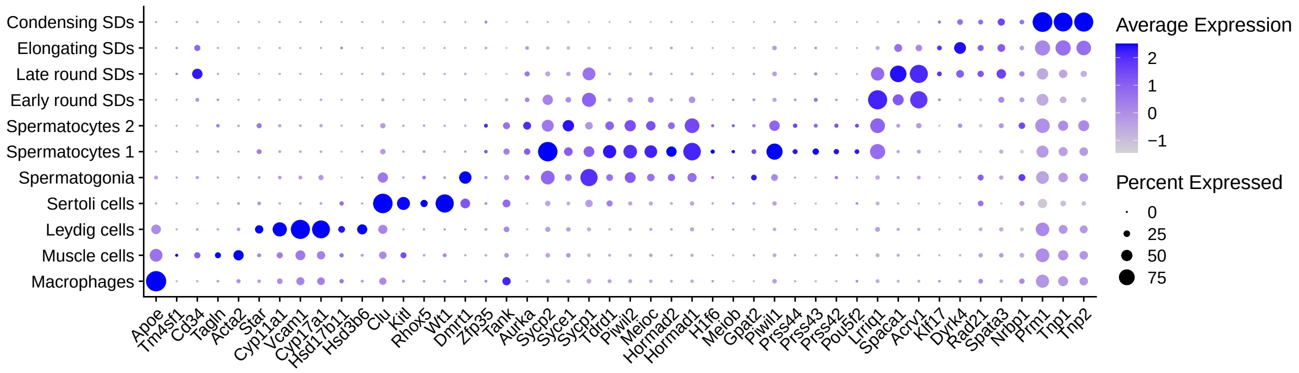


Supplementary Figure 3: Expression of known testis cell type marker genes identified by [Lukassen et al., 2018, Green et al., 2018] are shown in the different cell type clusters. SDs = spermatids

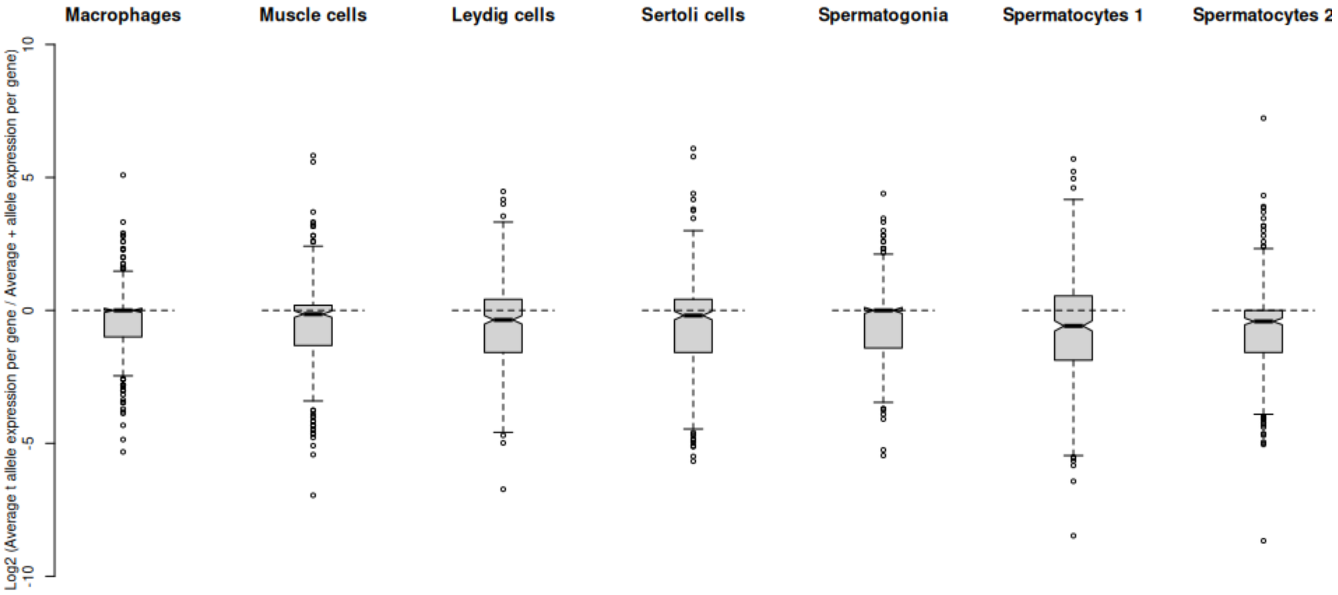


Supplementary Figure 4: Log2 ratios of average t allele versus average + allele expression for genes encoded in the t complex. Many genes show higher + allele than t allele expression in spermatocytes. SDs = spermatids

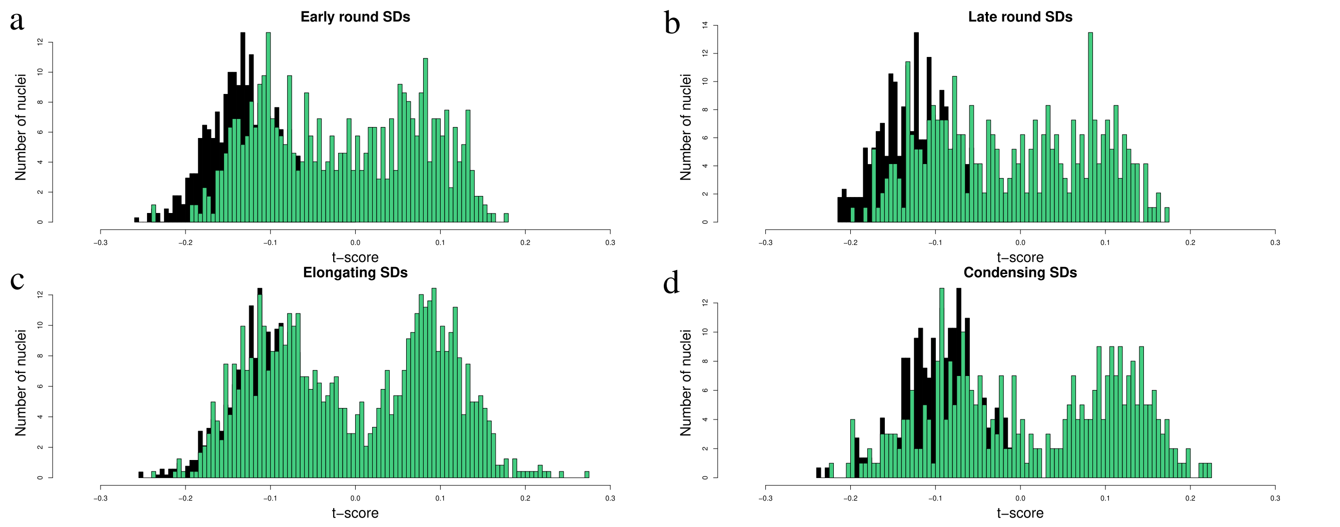


Supplementary Figure 5: Distributions of $t$-scores in haploid clusters of $+/t$ (green) samples and the $+/+$ (black) samples. SDs = spermatids

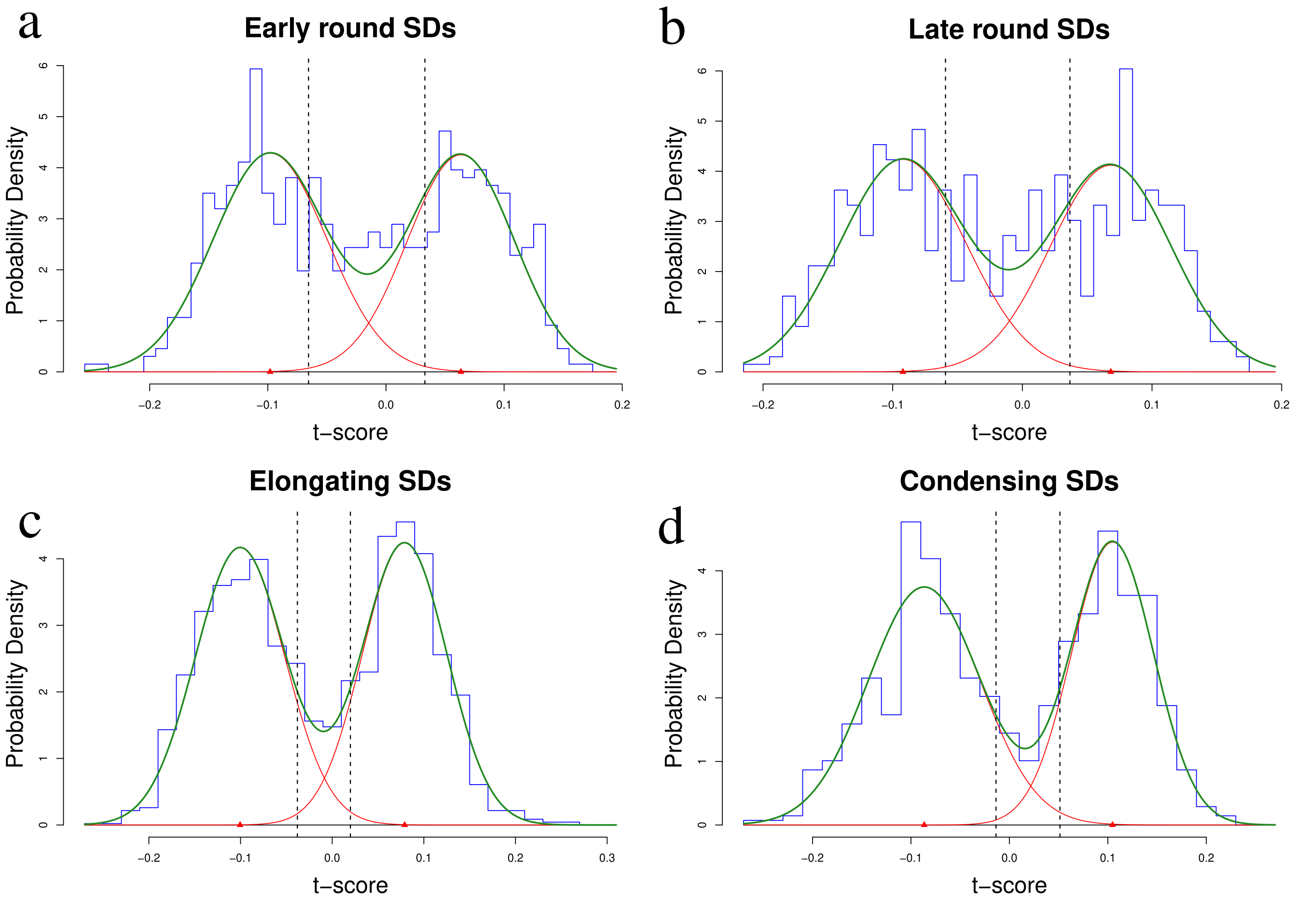


Supplementary Figure 6: Distributions of $t$-scores in haploid clusters of $+/t$ samples and normal distributions fitted to them. Blue histograms show the frequency distribution of $t$-scores. Red smooth lines represent the two normal distributions fitted to the bimodal distributions, while the green smooth lines show the bimodal distribution fitted to the data. Dotted lines mark the 90th and 10th percentiles of the fitted low-t-score and high-t-score dsitributions, respectively, which were used to call spermatid genotypes. Nuclei in the early and late round spermatid clusters showed a more unimodal distribution, therefore the 25th and 75th percentiles were used for genotype calling. SDs = spermatids

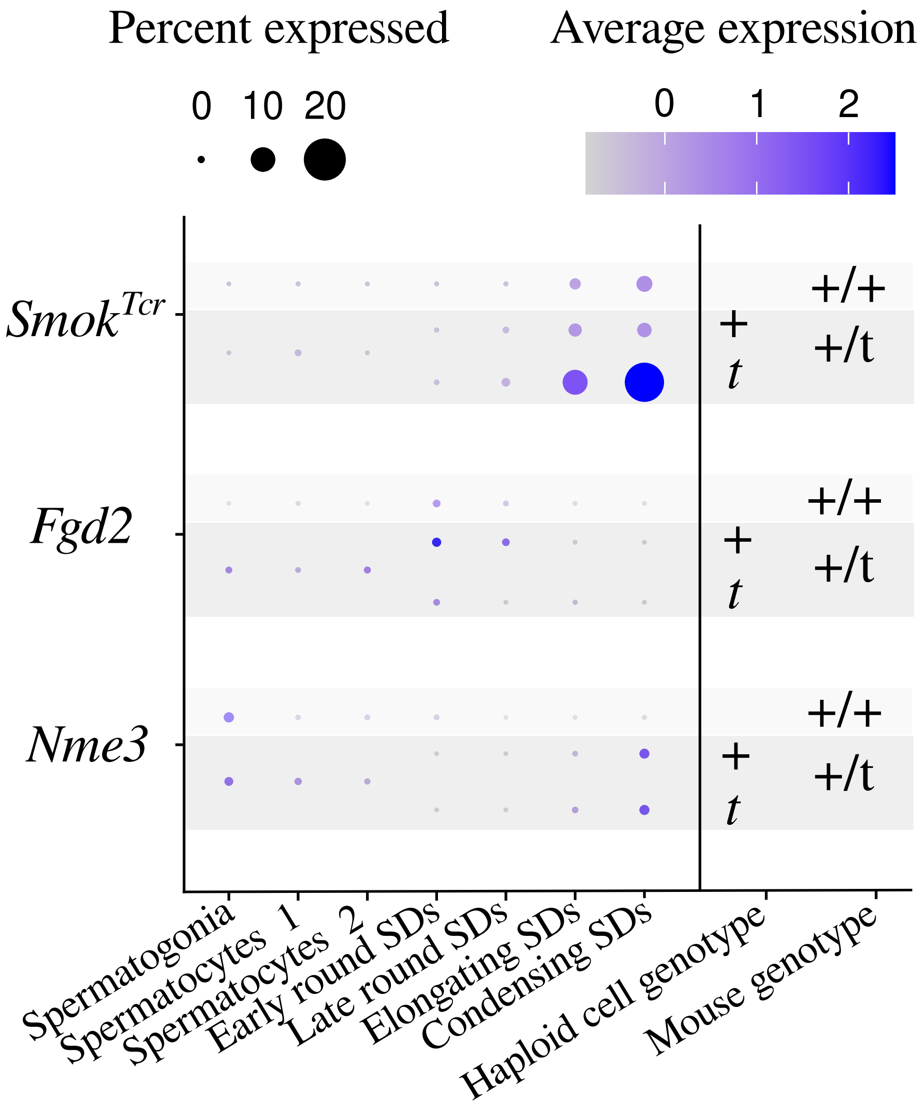


Supplementary Figure 7: Percent of nuclei per cell type expressing the previously identified (a) antidote of the *t*-haplotype, $Smok^{Tcr}$, and (b-c) poisons, (b) *Nme3* and (c) *Fgd2*, of the *t*-haplotype. SDs - spermatids

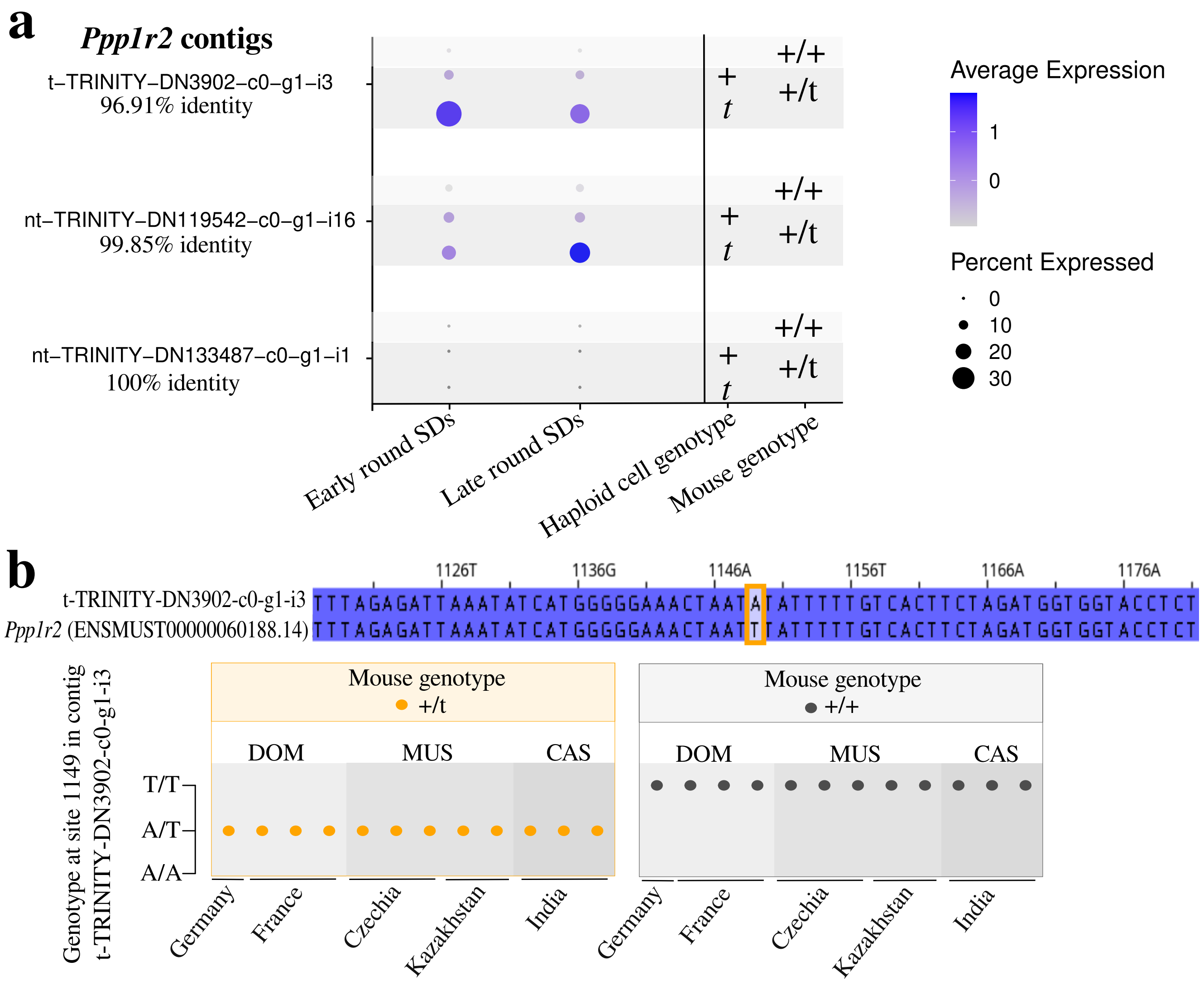


Supplementary Figure 8: Evidence for one of the assembled contigs of Ppp1r2 being specific to the *t*-haplotype. (a) Expression pattern of the three contigs representing *Ppp1r2* in our dataset, with the percent of identity of each to the longest transcript (ENSMUST0000006088.14) of the *Ppp1r2* gene in the GRCm39 reference genome. The size of each circle in the plot is proportional to the percent of nuclei expressing each contig, and the shading of purple denotes the average scaled expression level in each cell type. Only those cell types are shown where Ppp1r2 is differentially expressed between + and *t* genotypes (see Supplementary table 2). SD - spermatids. (b) The alignment of the highly diverged Ppp1r2 contig is shown to the reference transcript sequence ENSMUST0000006088.14 in the region where the T to A mutation lies at position 1149. In the lower panel the inferred genotypes of 12 +/*t* individuals and 12 +/+ individuals are shown from the dataset of [Harr et al., 2016]. Genotype was inferred by mapping genomic reads to the Ppp1r2 contig t-TRINITY-DN3902-c0-g1-i3 and calling variants using the tool *bcftools*. The source location of each mouse is indicated on the x axis, while the subspecies of each mouse is indicated above the plot with DOM - *M. m. domesticus*, MUS - *M. m. musculus* and CAS - *M. m. castaneus*.

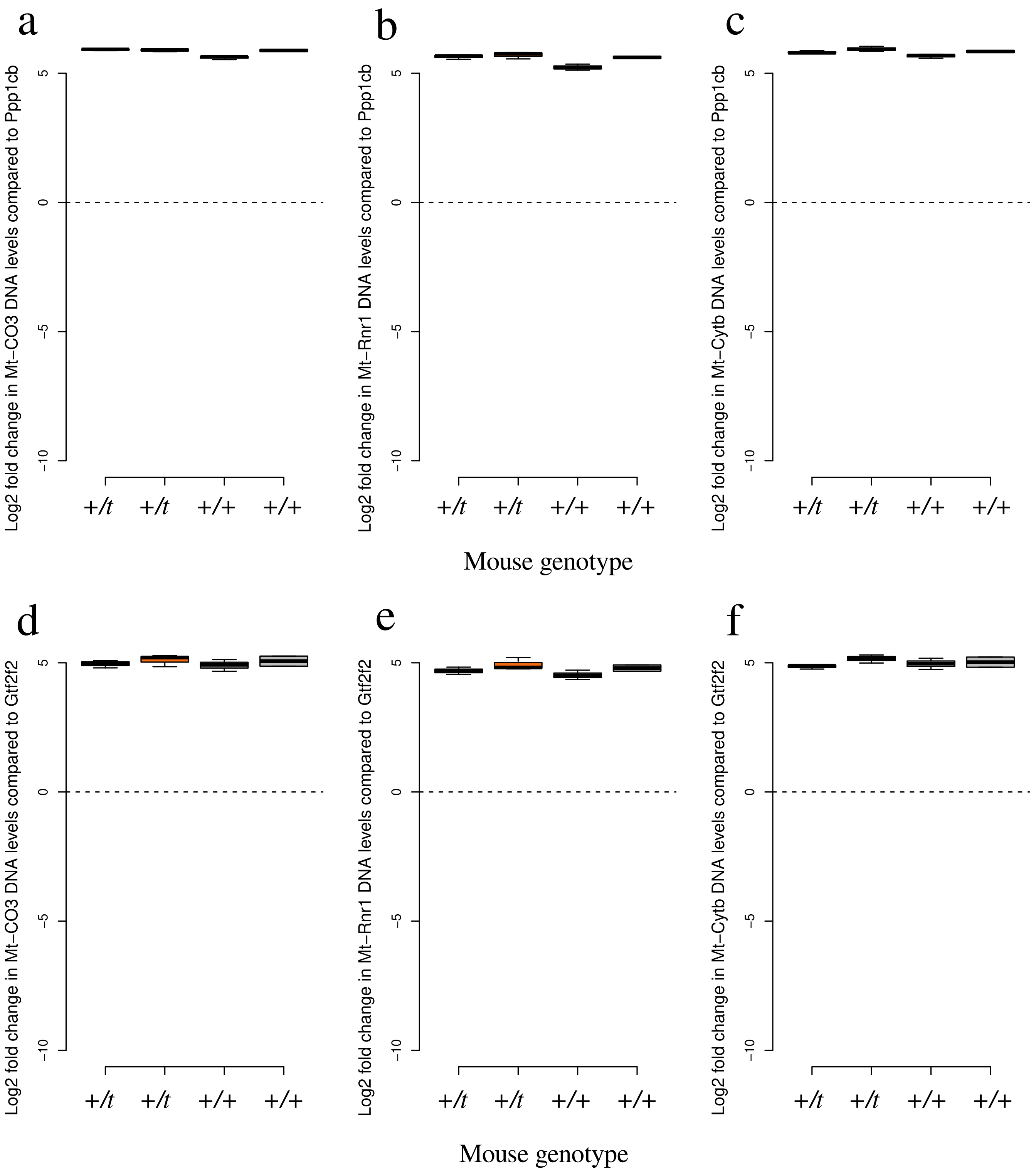


Supplementary Figure 9: Mitochondrial DNA copy number is not significantly different between +/*t* and +/+ testes. Quantitative PCR cycles were measured for primers designed for the mitochondrial genes (a,d) *Mt-Co3*, (b,e) *Mt-Rnr1* and (c,f) *Mt-Cytb* and the number of cycles of primers designed for the nuclear single copy genes (a-c) *Ppp1cb* or (d-f) *Gtf2f2* were subtracted. For each testis three technical replicates were taken and their values are shown in boxplots.

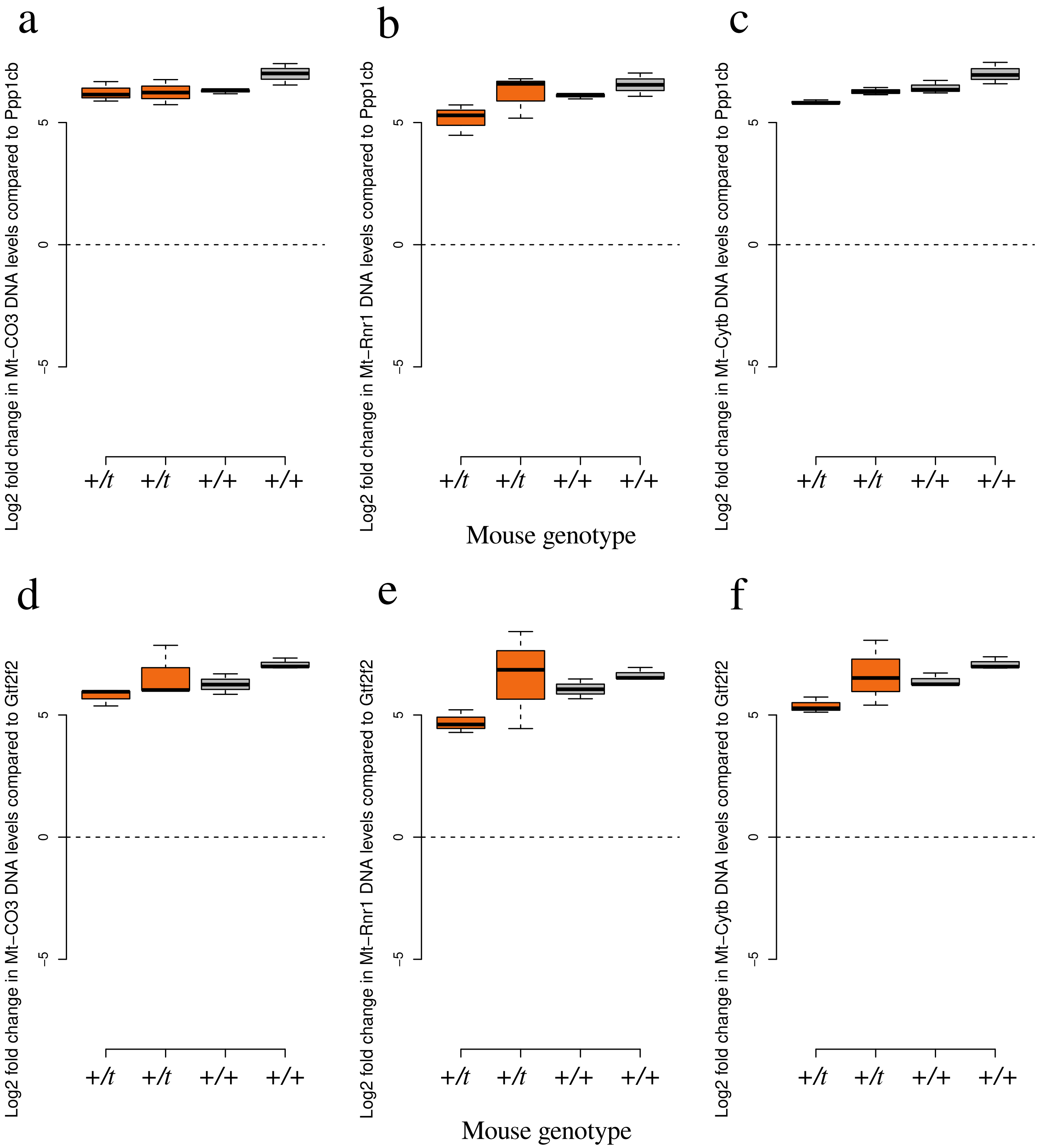


Supplementary Figure 10: Mitochondrial DNA copy number is not significantly different between +/*t* and +/+ epididymal sperm. Quantitative PCR cycles were measured for primers designed for the mitochondrial genes (a,d) *Mt-Co3*, (b,e) *Mt-Rnr1* and (c,f) *Mt-Cytb* and the number of cycles of primers designed for the nuclear single copy genes (a-c) *Ppp1cb* or (d-f) *Gtf2f2* were subtracted. For each epididymal sperm sample three technical replicates were taken and their values are shown in boxplots.

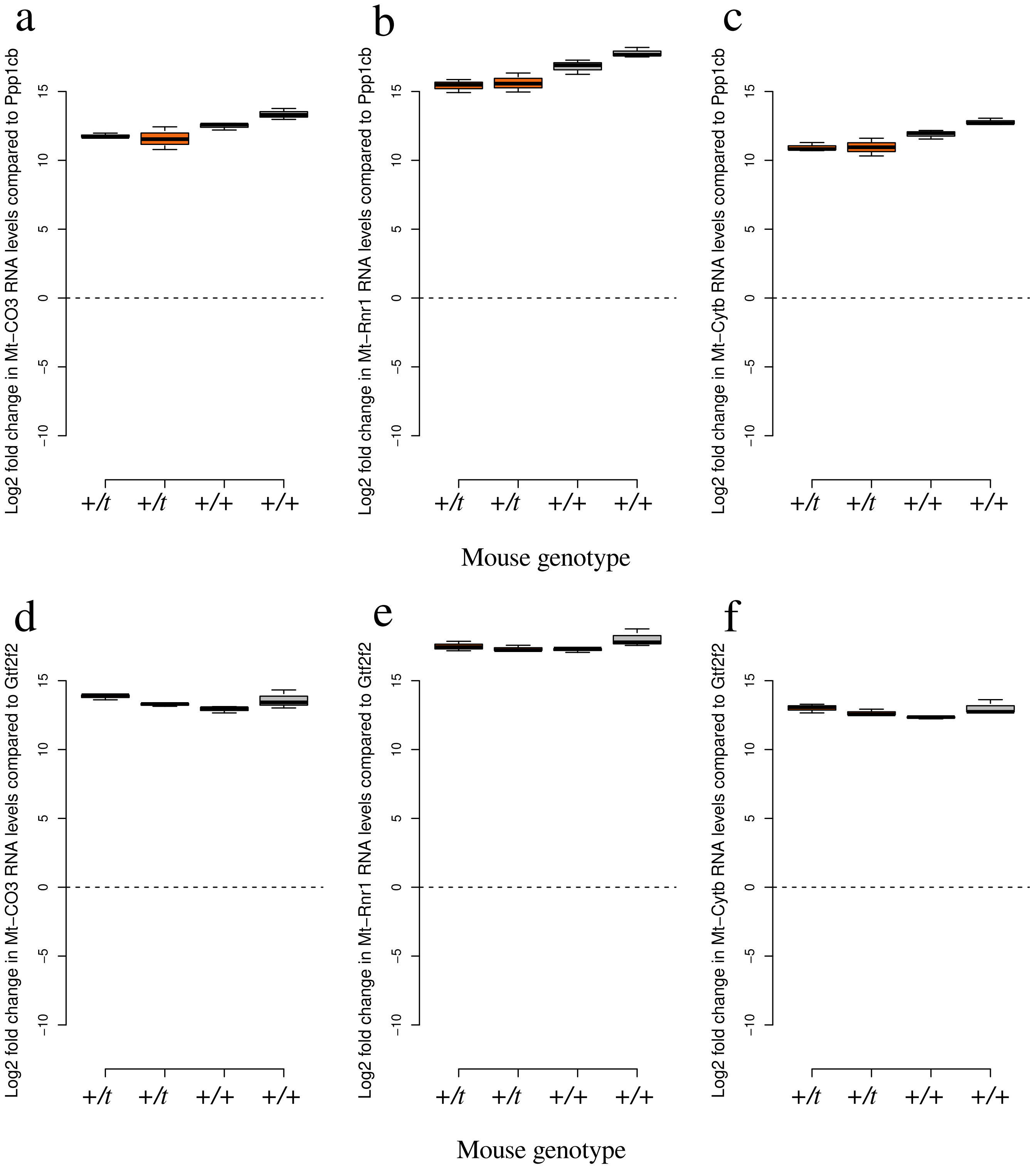


Supplementary Figure 11: Mitochondrial RNA levels are not significantly different between +/*t* and +/+ epididymal sperm. Quantitative PCR cycles were measured for primers designed for the mitochondrial genes (a,d) *Mt-Co3*, (b,e) *Mt-Rnr1* and (c,f) *Mt-Cytb* and the number of cycles of primers designed for the nuclear single copy genes (a-c) *Ppp1cb* or (d-f) *Gtf2f2* were subtracted. For each epididymal sperm sample three technical replicates were taken and their values are shown in boxplots.

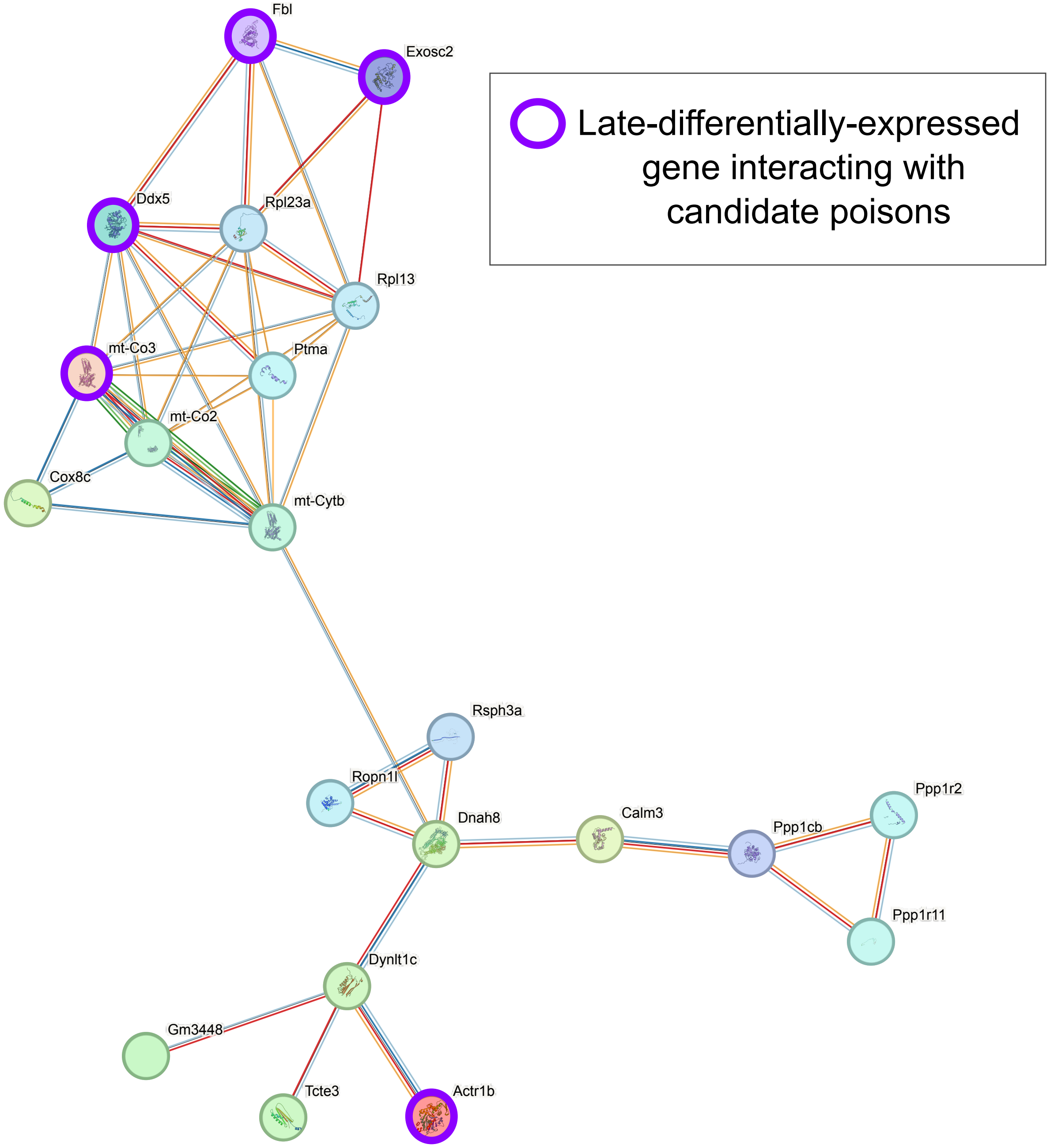


Supplementary Figure 12: Protein-protein interaction network computed by StringDB showing differentially expressed genes in germline cell types, which interact with each other. Candidate antidotes (proteins differentially expressed at the late stages of spermatogenesis), which interact with a candidate poison (proteins differentially expressed at the early stages of spermatogenesis), are shown by purple circles.

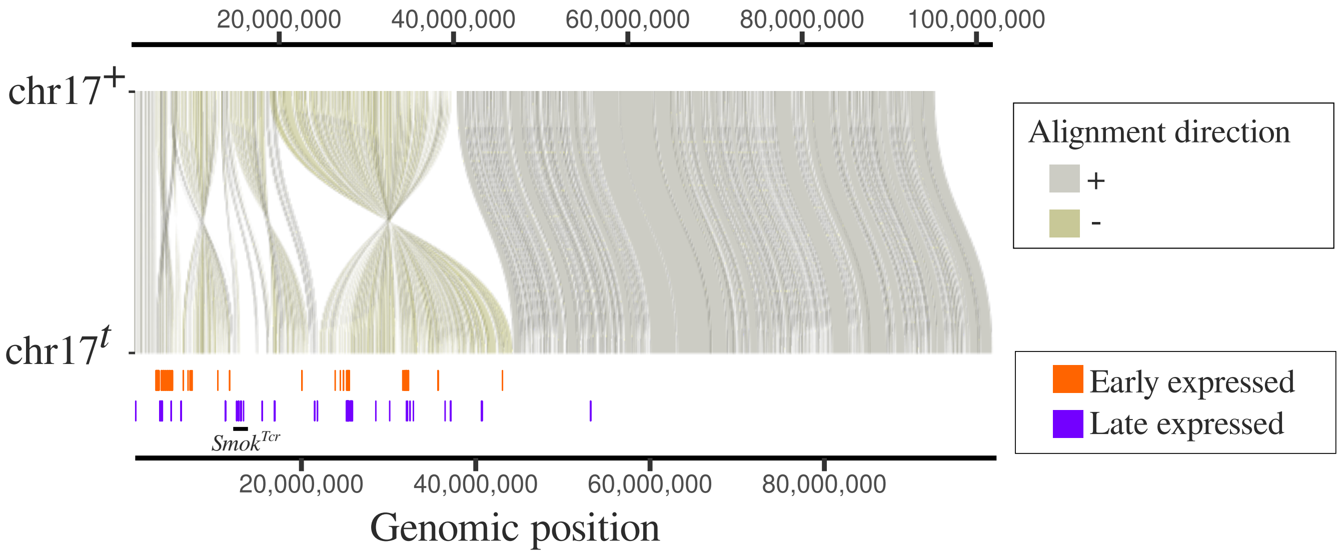


Supplementary Figure 13: Locations of candidate poisons and antidotes on the *t*-haplotype assembly of [Swanepoel et al., 2025]. Assembled contigs for each differentially expressed gene in the germline were mapped against the *t*-haplotype assembly and mappings longer than 300 basepairs and above 95% identity are shown for poisons (orange) and antidotes (purple). The region that encodes $Smok^{Tcr}$ is indicated on the bottom.

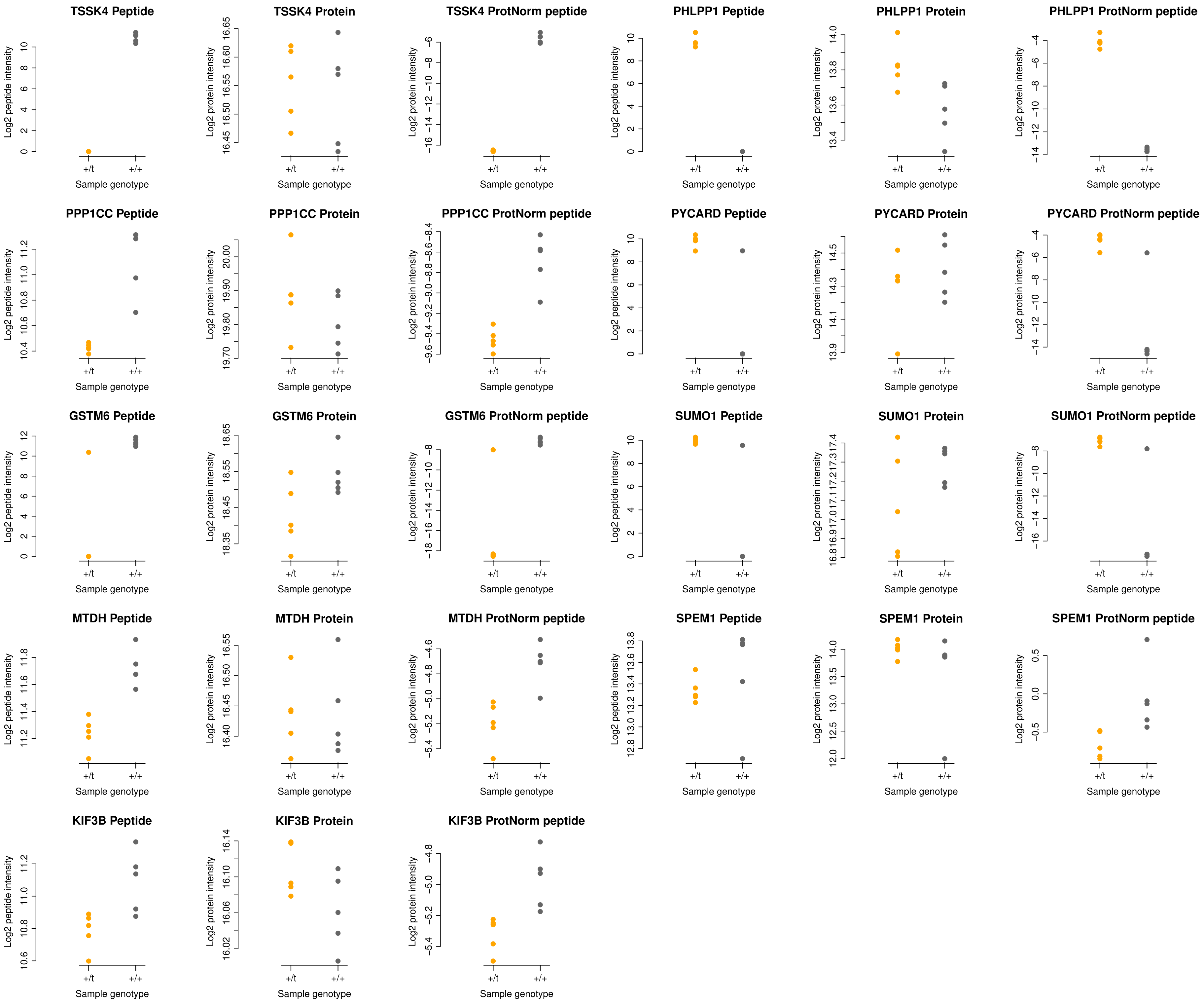


Supplementary Figure 14: Phosphopeptide, protein, and protein-abundance-normalized phosphopeptide abundance for the 11 differentially phosphorylated peptides between $+/t$ and $+/+$ testes.

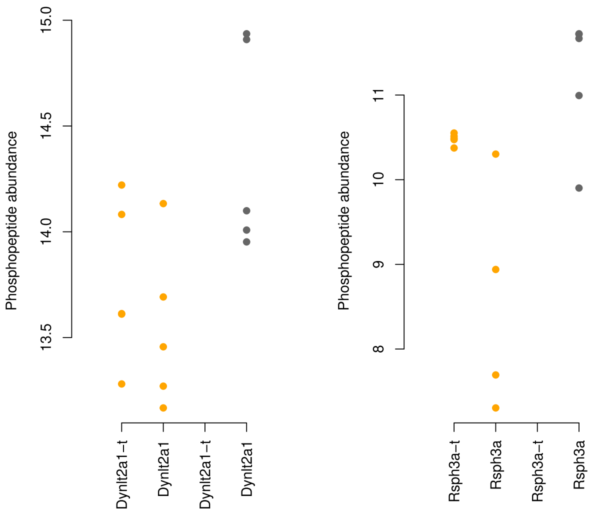


Supplementary Figure 15: Log2 intensities of *t*-specific and +-specific phosphopeptides of the *t*-complex-encoded dyneins and radial spoke proteins, DYNLT2A1 and RSPH3A, in +/*t* (orange) and +/+ (grey) testes.

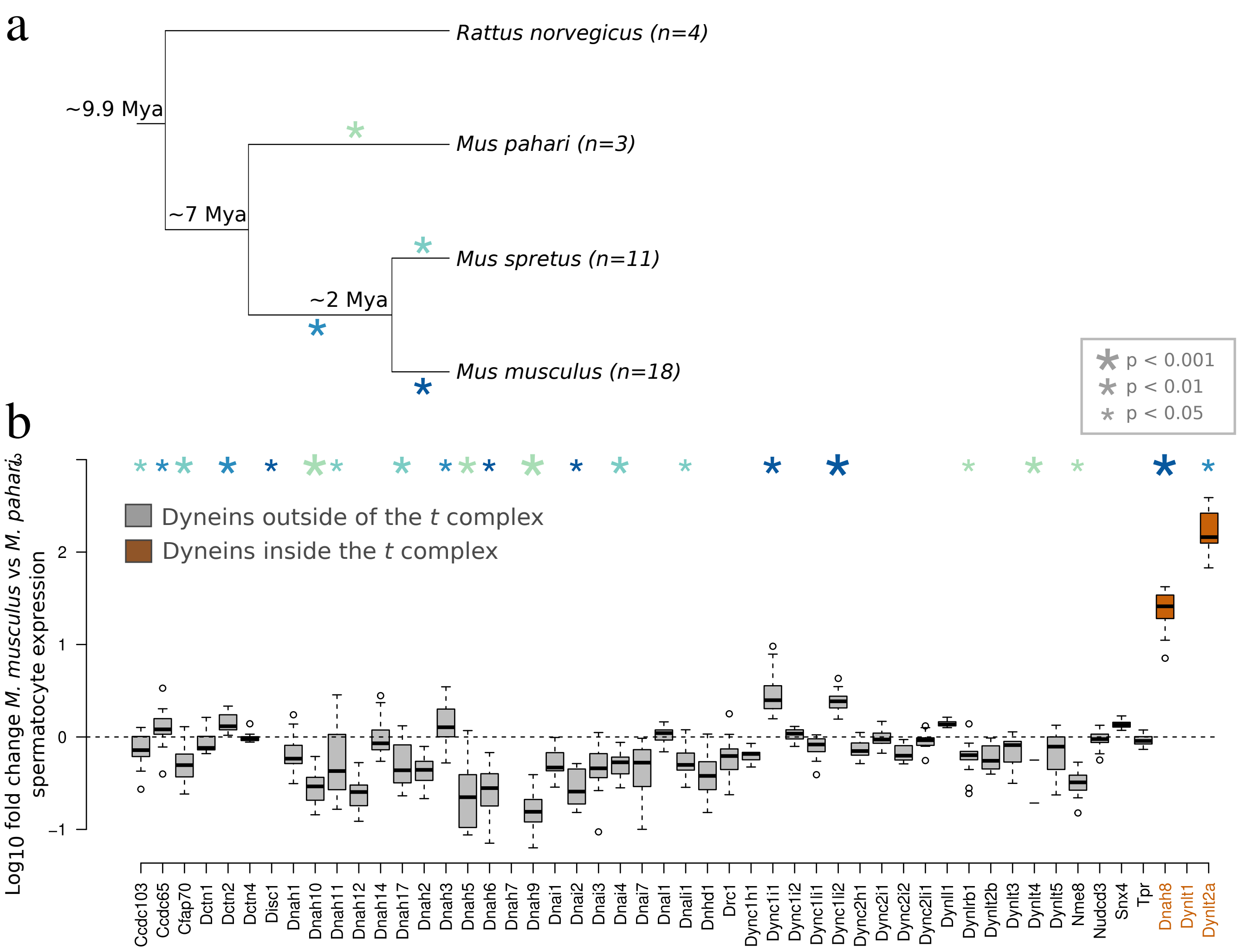


Supplementary Figure 16: The evolution of expression levels in spermatocytes of dynein complex genes in *Mus* phylogeny (a). (b) Genes with a significant shift in expression on a specific branch are marked stars, whose colors correspond to different branches of the *Mus* phylogeny shown in (a). The sizes of the stars are negatively proportional to the associated p-values, which result from likelihood ratio tests comparing a model with a shift in mean expression versus a null model of no shift. For modeling, we used the R package evemodel [Rohlfs and Nielsen, 2015].

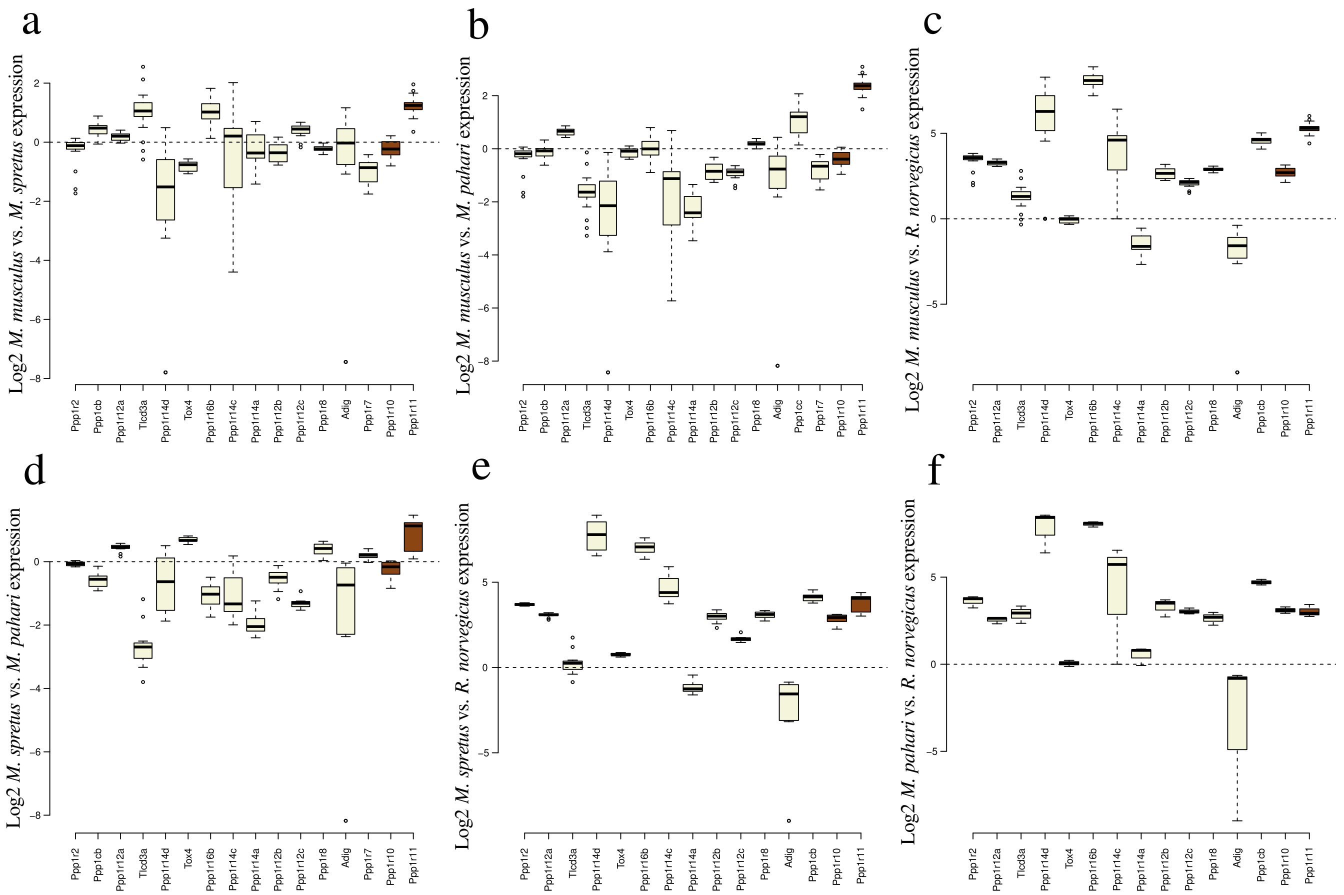


Supplementary Figure 17: Boxplots of log2 fold changes in spermatocyte expression of PP1 complex genes between pairs of species. (a) *M. musculus* vs. *M. spretus*, (b) *M. musculus* vs. *M. pahari*, (c) *M. musculus* vs. *R. norvegicus*, (d) *M. spretus* vs. *M. pahari*, (e) *M. spretus* vs. *R. norvegicus* and (f) *M. pahari* vs. *R. norvegicus*. PP1 complex genes in the *t* complex are shown in brown, while those encoded outside the *t* complex are shown in beige.

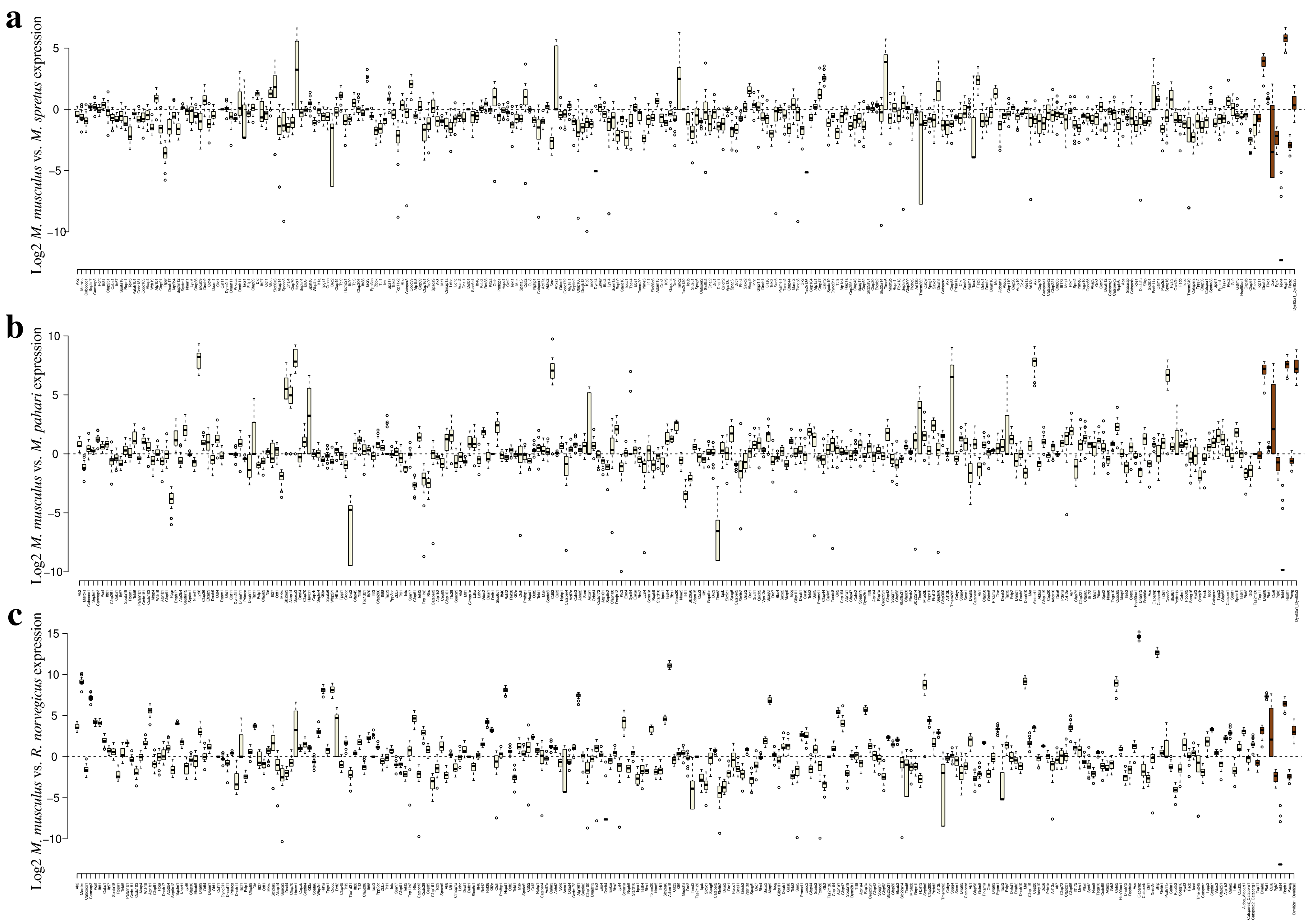


Supplementary Figure 18: Boxplots of log2 fold changes in spermatocyte expression of genes in the "motile cilium" gene ontology category between pairs of species. (a) *M. musculus* vs. *M. spretus*, (b) *M. musculus* vs. *M. pahari*, (c) *M. musculus* vs. *R. norvegicus*, (d) *M. spretus* vs. *M. pahari*, (e) *M. spretus* vs. *R. norvegicus* and (f) *M. pahari* vs. *R. norvegicus*. Motile cilium genes in the *t* complex are shown in brown, while those encoded outside the *t* complex are shown in beige. For better visual representation, the first half of the genes are shown, and the second half is shown in Supplementary figure 15.

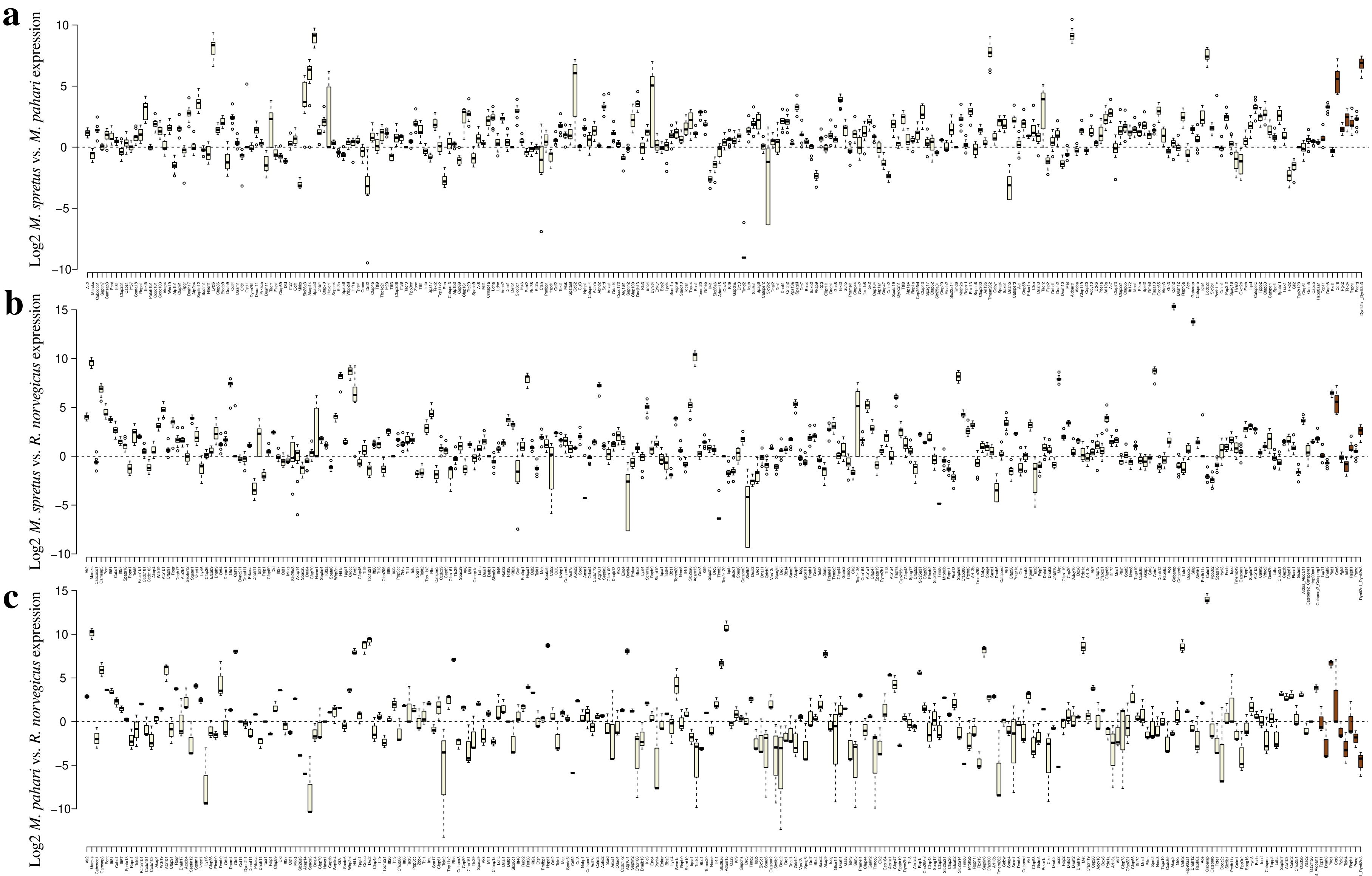


Supplementary Figure 19: Boxplots of log2 fold changes in spermatocyte expression of genes in the "motile cilium" gene ontology category between pairs of species. (a) *M. musculus* vs. *M. spretus*, (b) *M. musculus* vs. *M. pahari*, (c) *M. musculus* vs. *R. norvegicus*, (d) *M. spretus* vs. *M. pahari*, (e) *M. spretus* vs. *R. norvegicus* and (f) *M. pahari* vs. *R. norvegicus*. Motile cilium genes in the *t* complex are shown in brown, while those encoded outside the *t* complex are shown in beige. For better visual representation, the second half of the genes are shown, and the first half is shown in Supplementary figure 14.

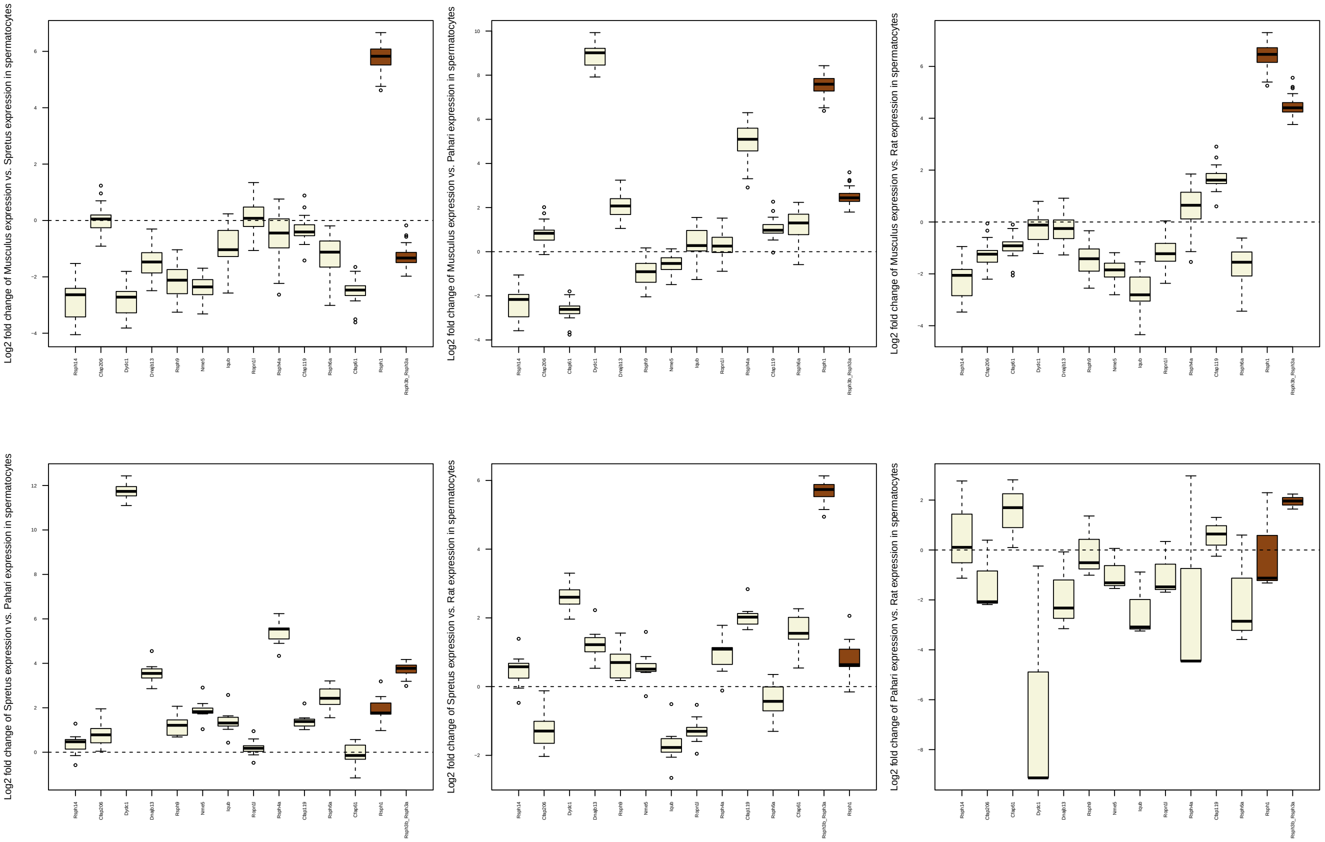


Supplementary Figure 20: Boxplots of log2 fold changes in spermatocyte expression of radial spoke genes between pairs of species. (a) *M. musculus* vs. *M. spretus*, (b) *M. musculus* vs. *M. pahari*, (c) *M. musculus* vs. *R. norvegicus*, (d) *M. spretus* vs. *M. pahari*, (e) *M. spretus* vs. *R. norvegicus* and (f) *M. pahari* vs. *R. norvegicus*. Genes in the *t* complex are shown in brown, while those encoded outside the *t* complex are shown in beige.

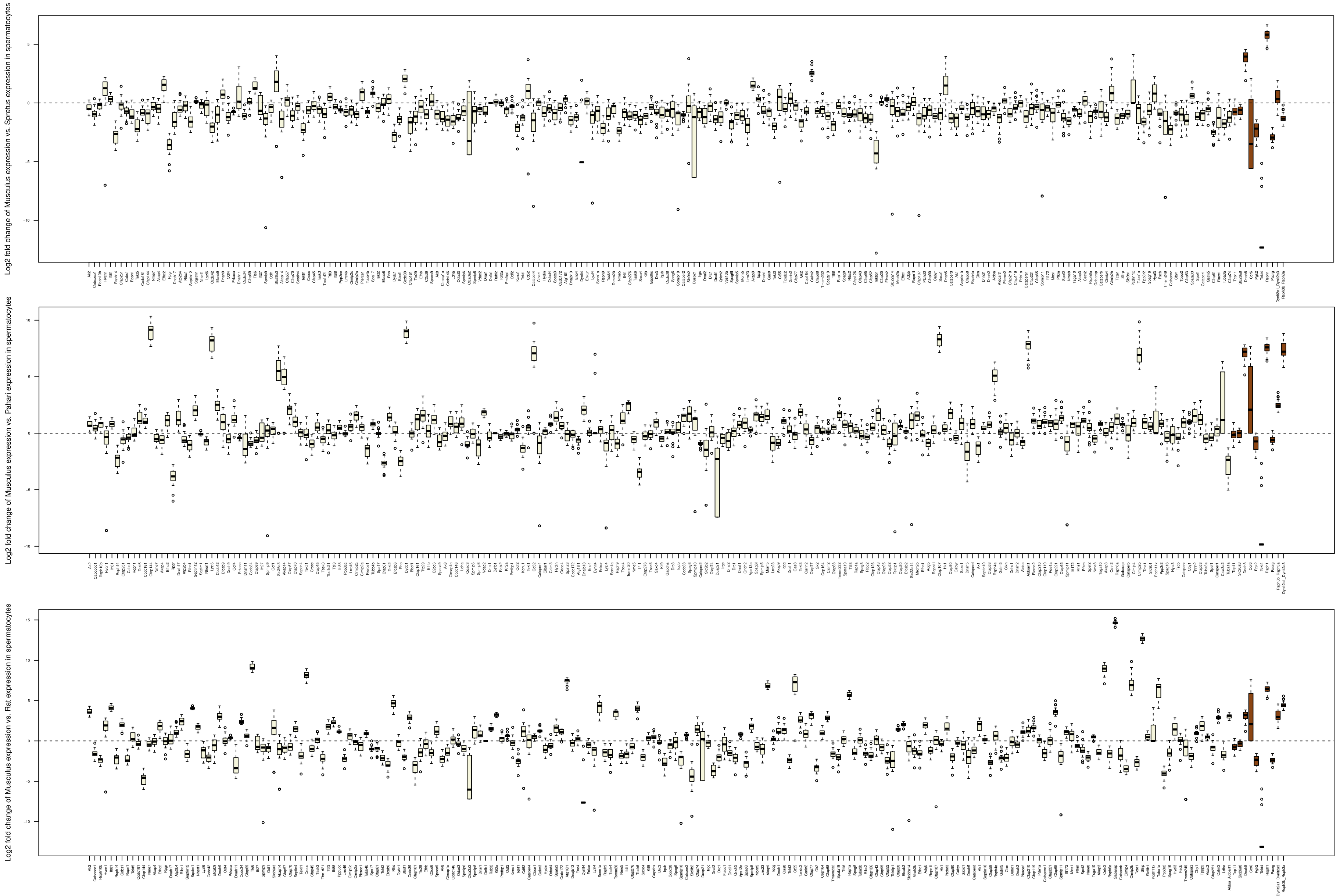


Supplementary Figure 21: Boxplots of log2 fold changes in spermatocyte expression of genes in the "9+2 motile cilium" gene ontology category between pairs of species (see y axis labels). Genes in the $t$ complex are shown in brown, while those encoded outside the $t$ complex are shown in beige.

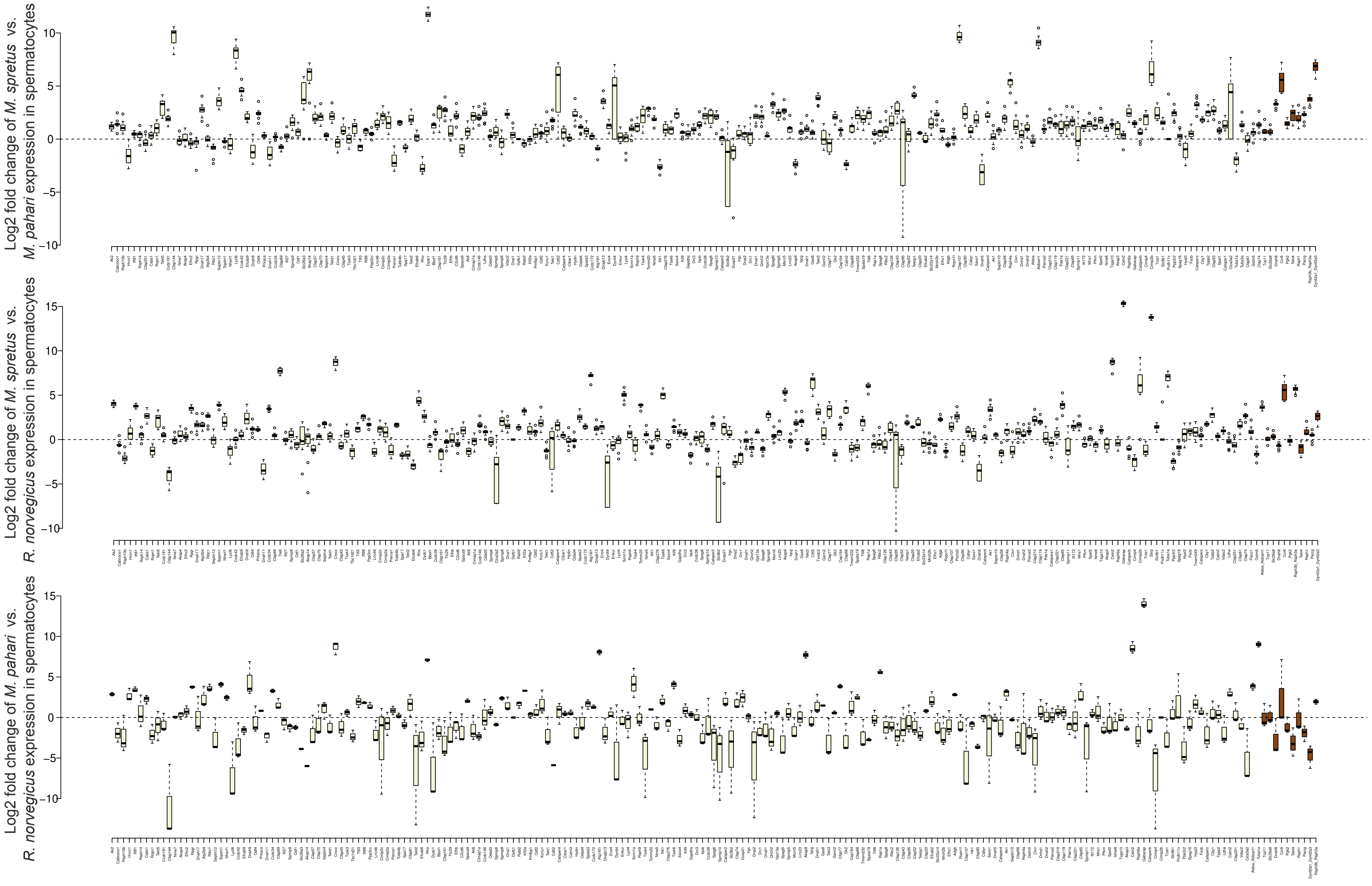


Supplementary Figure 22: Boxplots of log2 fold changes in spermatocyte expression of genes in the "9+2 motile cilium" gene ontology category between pairs of species (see y axis labels). Genes in the $t$ complex are shown in brown, while those encoded outside the $t$ complex are shown in beige.

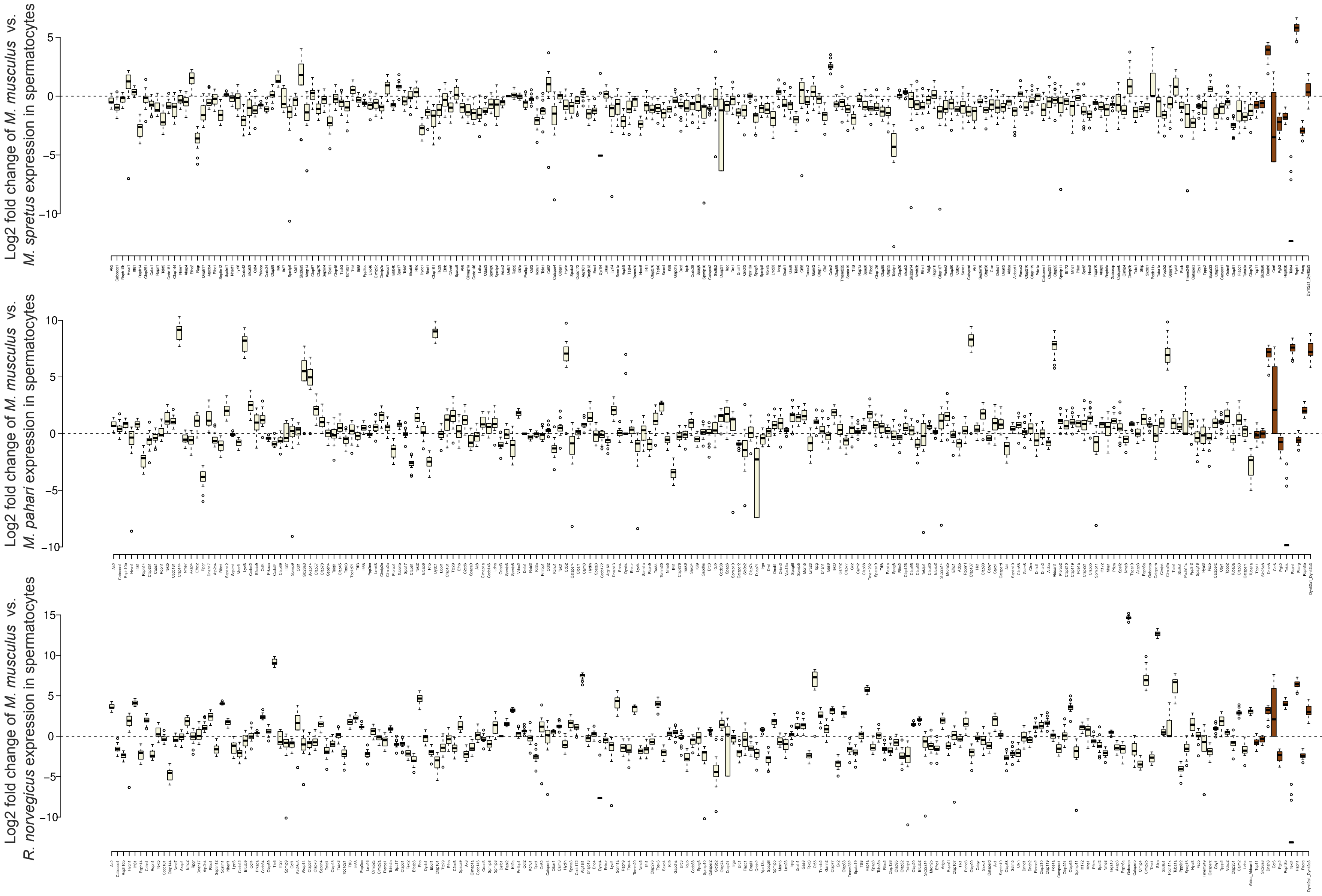


Supplementary Figure 23: Boxplots of log2 fold changes in spermatocyte expression of genes in the "sperm flagellum" gene ontology category between pairs of species (see y axis labels). Genes in the *t* complex are shown in brown, while those encoded outside the *t* complex are shown in beige.

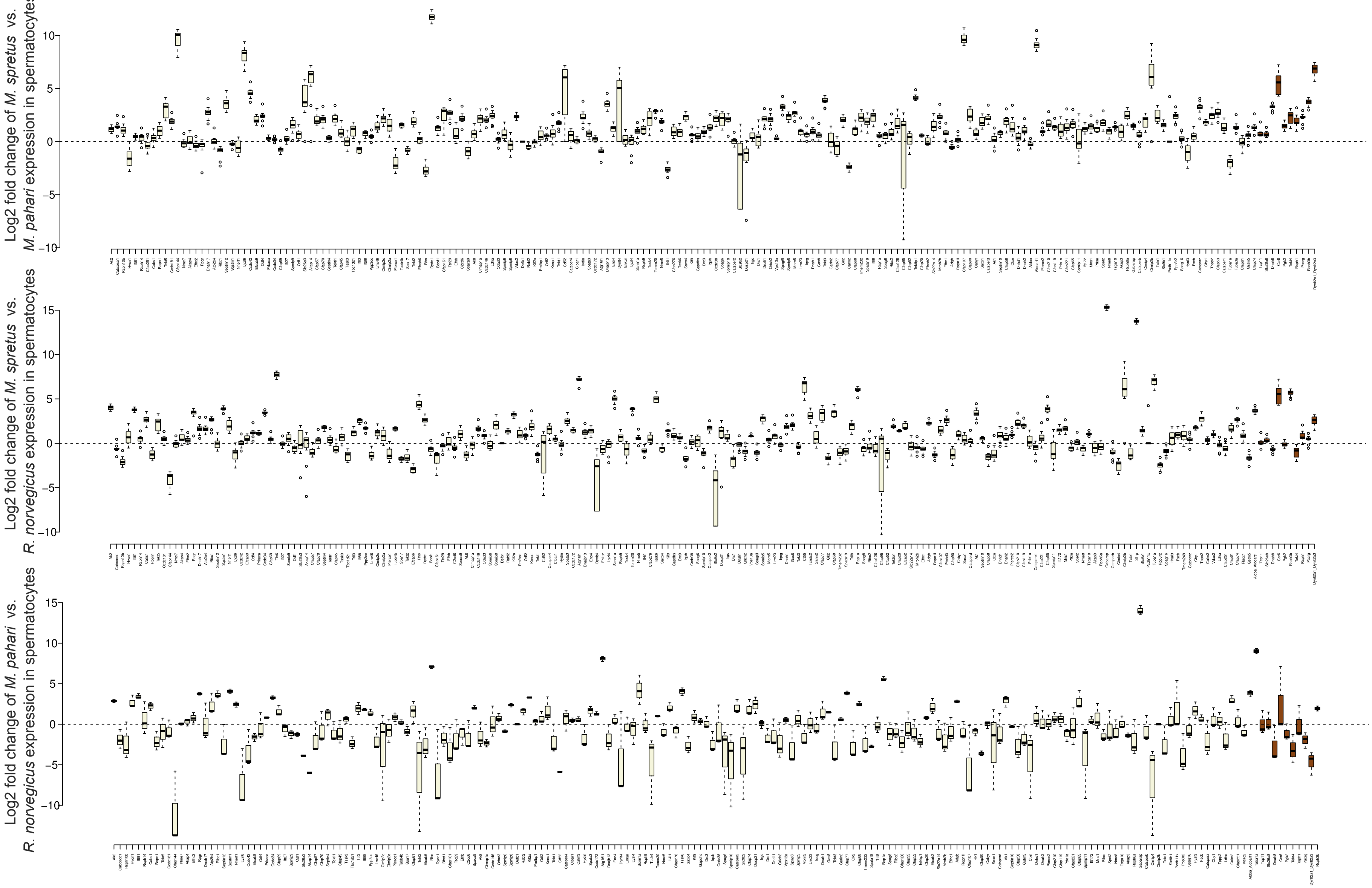


Supplementary Figure 24: Boxplots of log2 fold changes in spermatocyte expression of genes in the "sperm flagellum" gene ontology category between pairs of species (see y axis labels). Genes in the $t$ complex are shown in brown, while those encoded outside the $t$ complex are shown in beige.

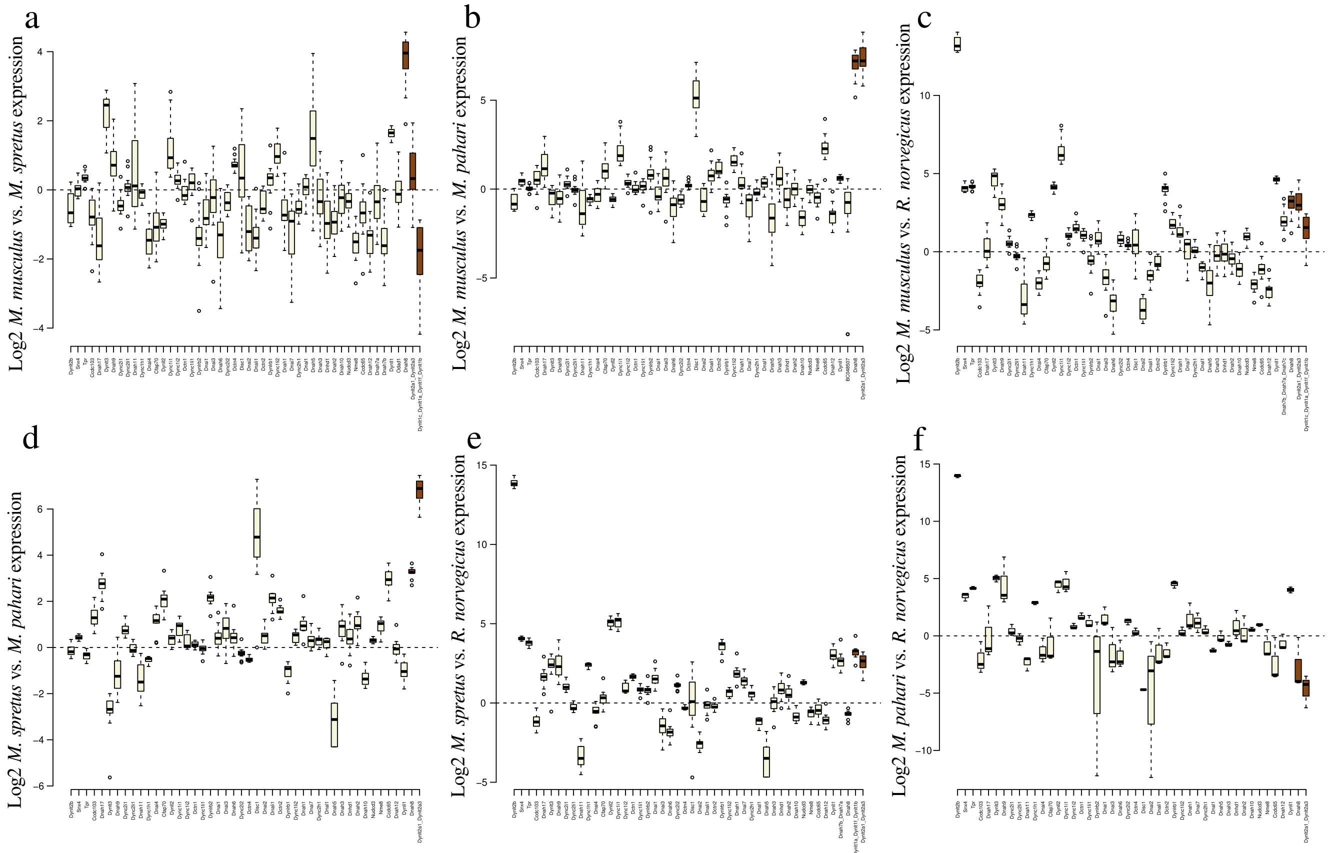


Supplementary Figure 25: Boxplots of log2 fold changes in spermatocyte expression of dynein complex genes between pairs of species. (a) *M. musculus* vs. *M. spretus*, (b) *M. musculus* vs. *M. pahari*, (c) *M. musculus* vs. *R. norvegicus*, (d) *M. spretus* vs. *M. pahari*, (e) *M. spretus* vs. *R. norvegicus* and (f) *M. pahari* vs. *R. norvegicus*. Dynein complex genes in the *t* complex are shown in brown, while those encoded outside the *t* complex are shown in beige.

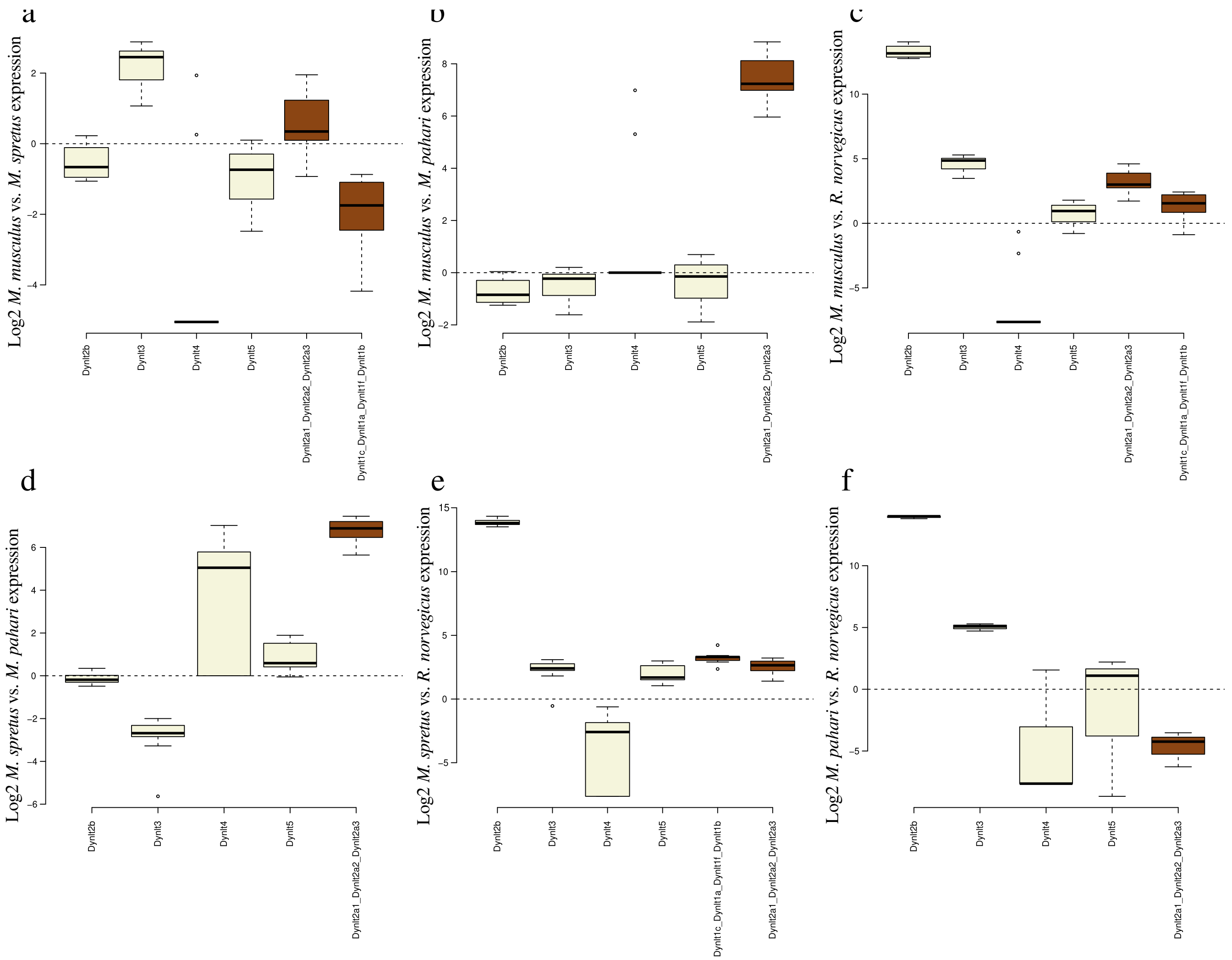


Supplementary Figure 26: Boxplots of log2 fold changes in spermatocyte expression of Tctex-family genes between pairs of species. (a) *M. musculus* vs. *M. spretus*, (b) *M. musculus* vs. *M. pahari*, (c) *M. musculus* vs. *R. norvegicus*, (d) *M. spretus* vs. *M. pahari*, (e) *M. spretus* vs. *R. norvegicus* and (f) *M. pahari* vs. *R. norvegicus*. Tctex-family genes in the *t* complex are shown in brown, while those encoded outside the *t* complex are shown in beige.

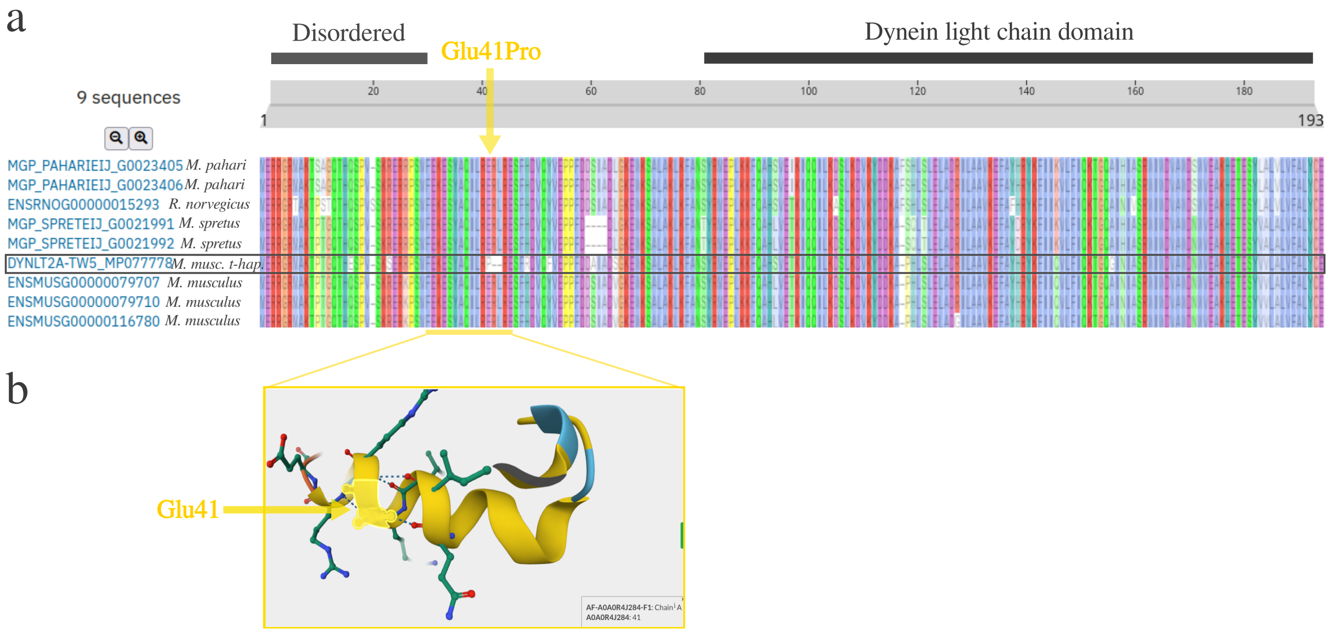


Supplementary Figure 27: Alignment of Dynlt2a protein sequences from the *M. musculus* GRCm39 reference genome, the published *t*-haplotype sequence, *R. norvegicus*, *M. pahari* and *M. spretus*. Structural domains are indicated based on the annotations from UniProt. The substitution of Glu41Pro in DYNLT2A$^{T}$ is indicated with a yellow arrow and the Alphafold prediction of the affected amino acid in its three dimensional structure is depicted.

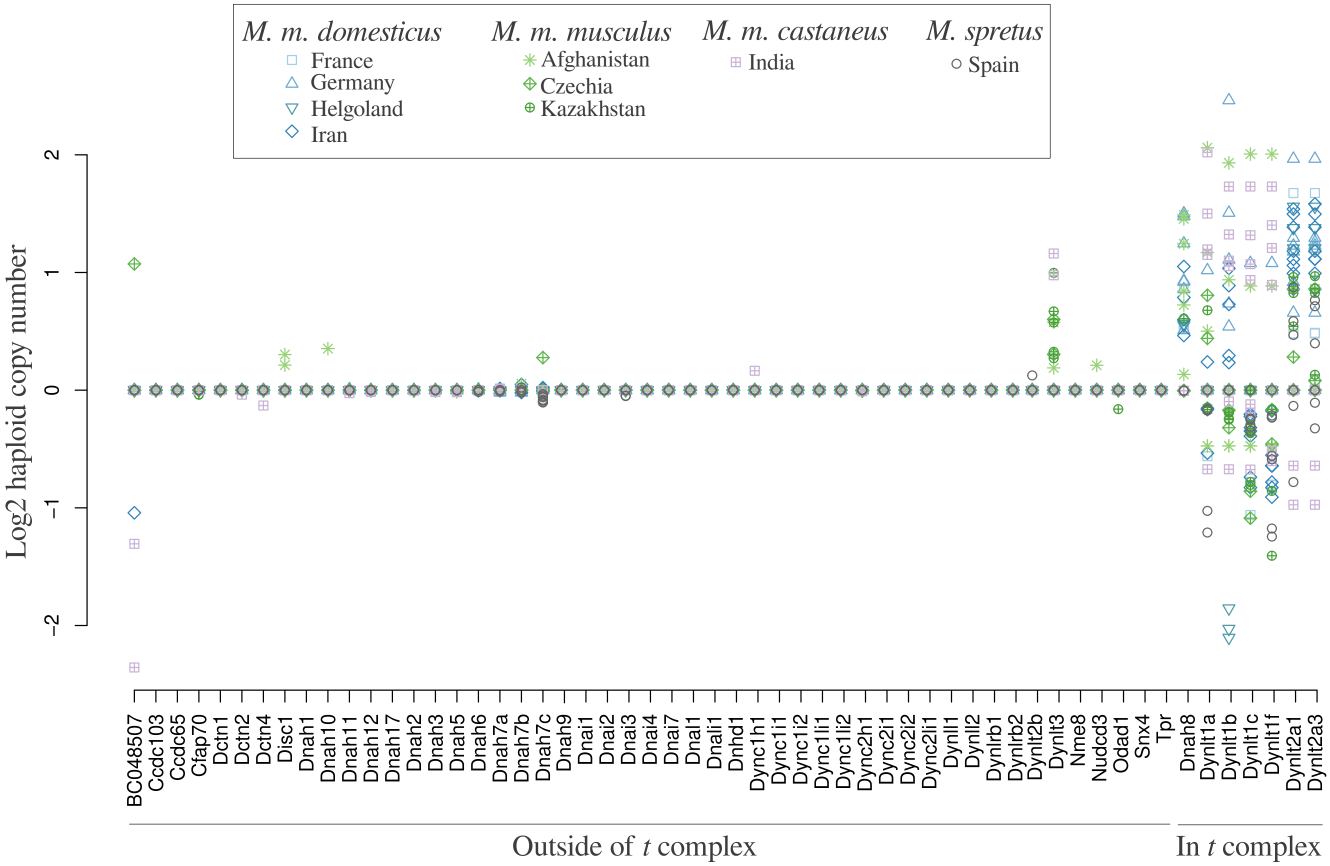


Supplementary Figure 28: Haploid copy number of dynein complex genes in and outside of the *t* complex in wild-caught mice from all three house mouse subspecies and the sister species, *Mus spretus*, obtained from the publication of [Harr et al., 2016]. The geographical location of each captured mouse is indicated with different symbols and colors shown in the figure legend. Haploid copy number estimates were computed by Harr and colleagues based on DNA-seq read coverages using the software CNVnator [Harr et al., 2016]. For each dynein coding sequence we averaged the estimated copy number for each overlapping CNV window.

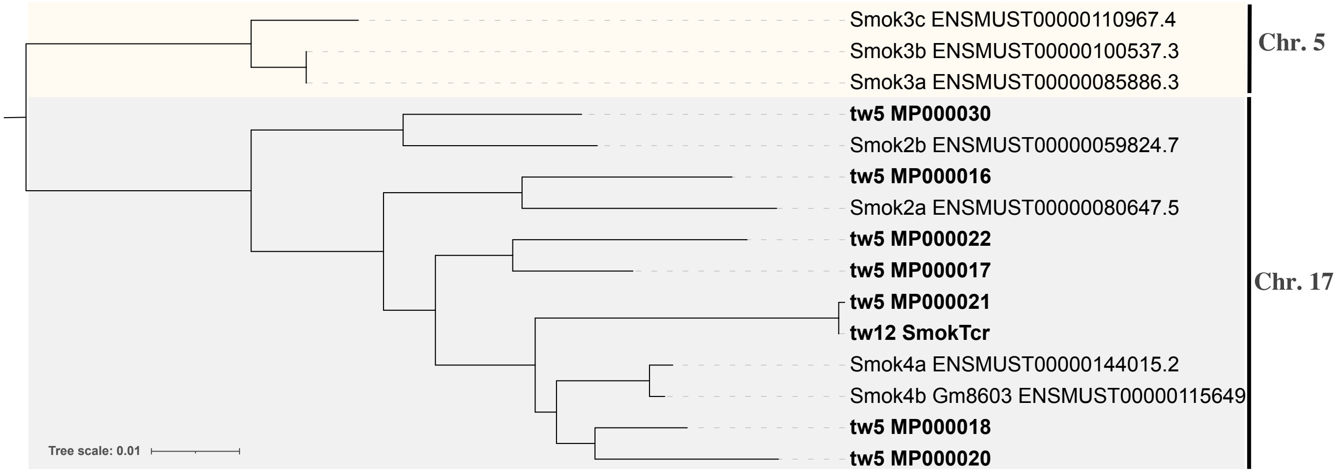


Supplementary Figure 29: Phylogenetic tree of the coding sequences of *Smok* genes in the *t*-haplotype assembly published by [Swanepoel et al., 2025] (shown in bold) and in the house mouse reference genome GRCm39. The chromsome on which each gene is encoded is shown on the right of the figure. The *Smok* genes in the *t*-haplotype assembly were extracted using the tools miniprot and gffread, and the phylogenetic tree was constructed with the software IQTree.

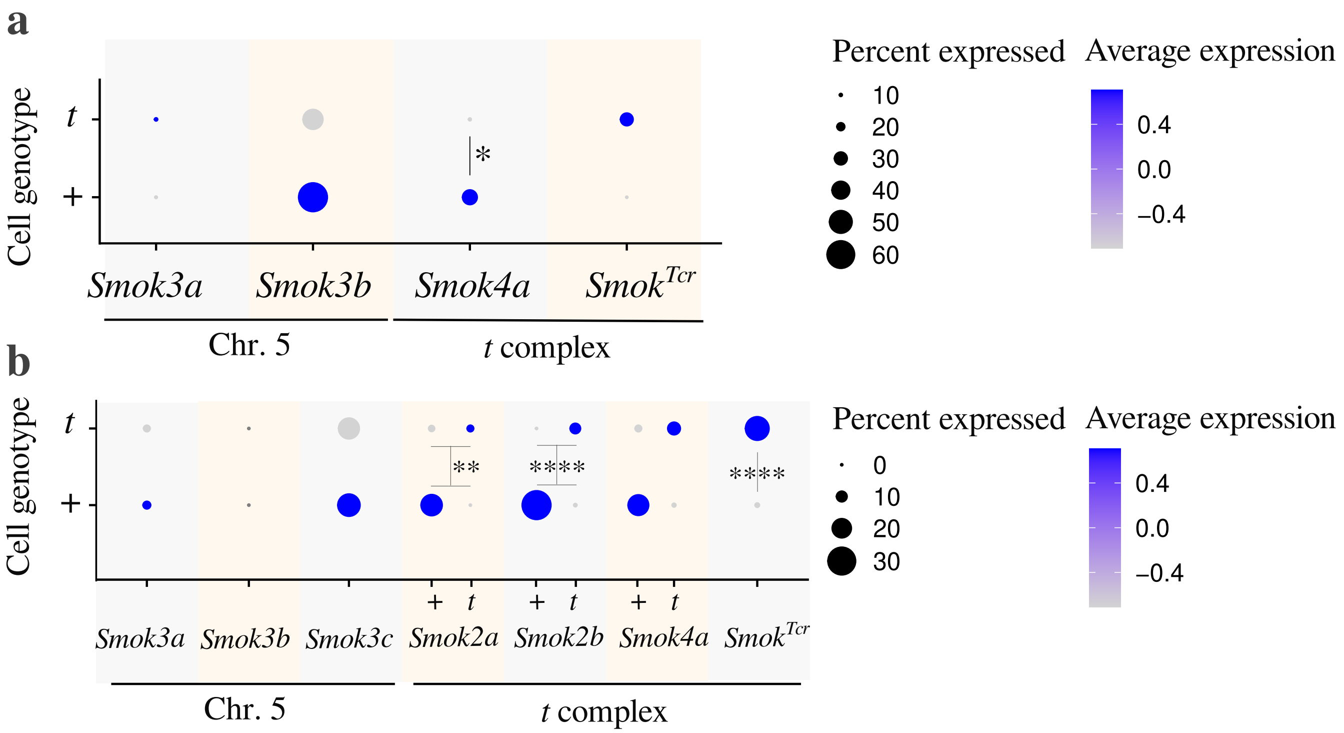


Supplementary Figure 30: (a) The percent of condensing spermatids of either + or *t* genotype expressing assembled transcripts annotated as *Smok* genes (*Smok2a* and *Smok2b* were not separately assembled ). The scaled mean expression per nucleus is shown by the color of the circle. *Smok4a* shows significantly higher expression in *t* vs. + condensing spermatids (adjusted p-value = 0.01, likelihood ratio test of models fitted to expression values by the MAST R package) (b) The percent of condensing spermatids of either + or *t* genotype expressing *Smok* genes in the mouse genome, and their pseudo-t-alleles. The scaled mean expression per nucleus is shown by the color of the circle. Genes with significant differences in expression between + and *t* spermatids are indicated with asterisks in the dotplot (Wilcoxon rank sum test of the summed *t* and + allele expression of + vs. *t* condensing spermatids).

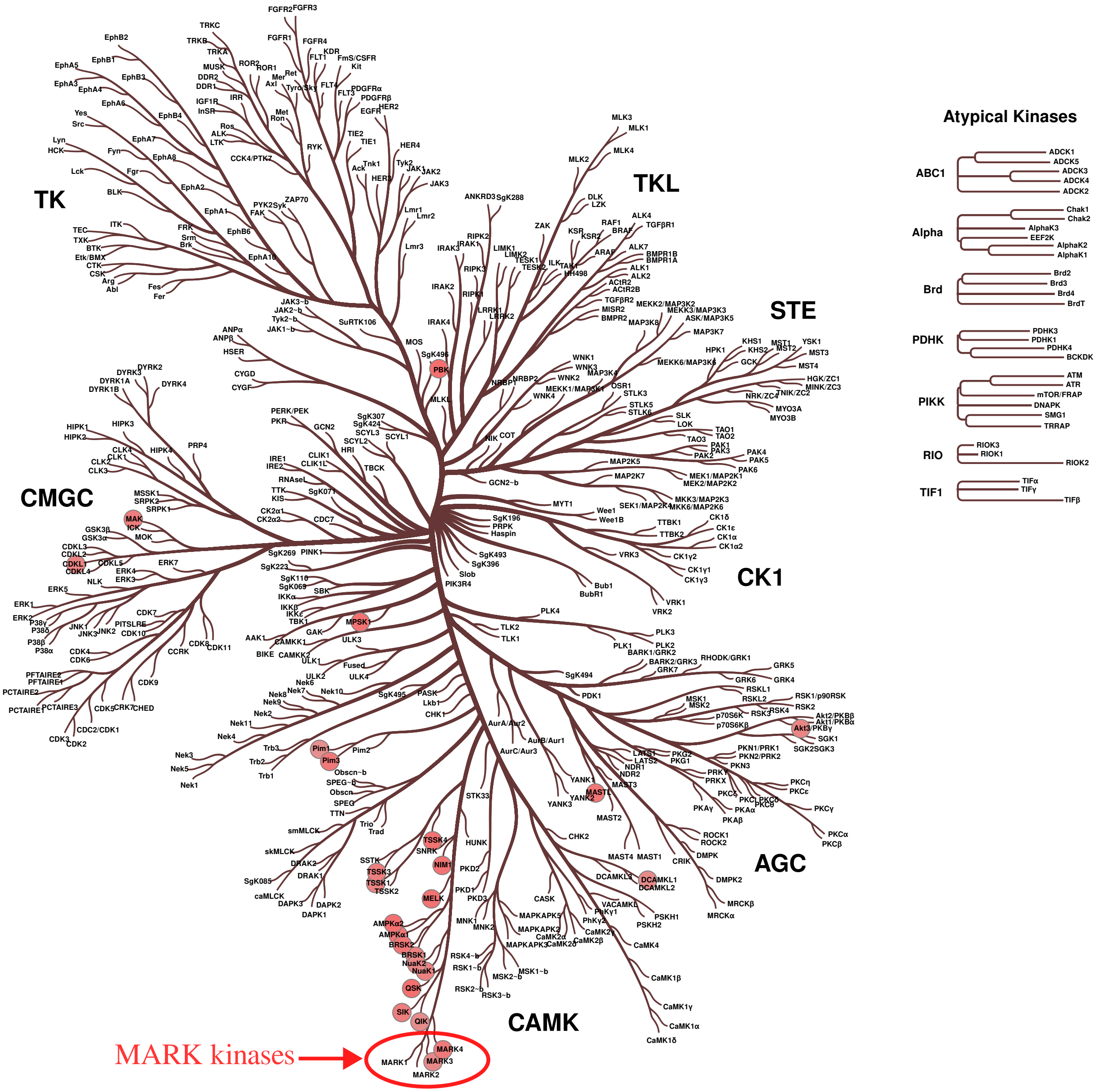


Supplementary Figure 31: Phylogenetic tree of kinases with red bubbles indicating those that are in the top 25th percentile of the most likely phosphorylators of S272 on PP1CC2. A red circle on the bottom of the tree indicates the MARK (Microtubule Affinity-Regulating Kinases) kinases, to which SMOK proteins show highest similarity.

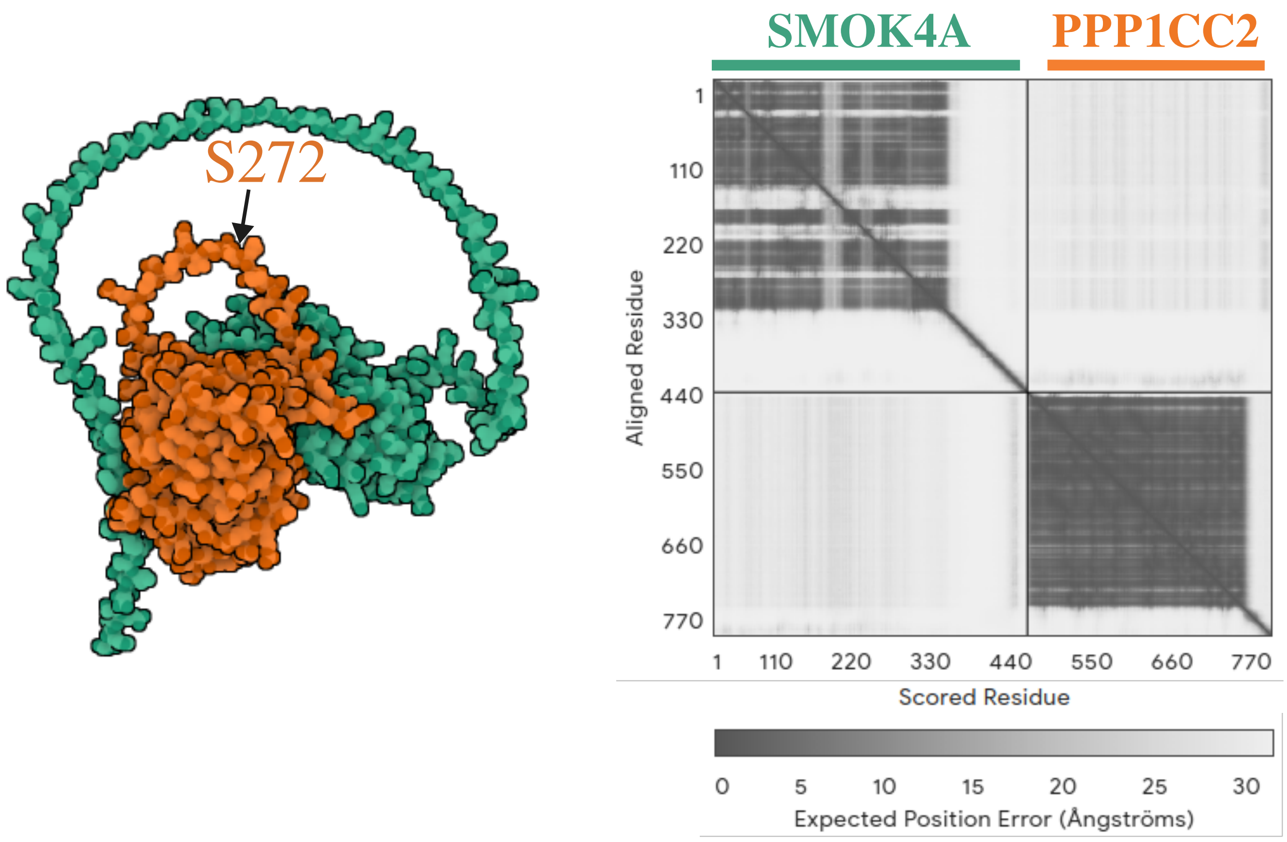


Supplementary Figure 32: Three dimensional model of the binding of (a) SMOK2A (green) and PPP1CC2 (orange) and (b) SMOK$^{TCR}$ (green) and PPP1CC2 (orange), as predicted by AlphaFold. The differentially phosphorylated S272 on PP1 is indicated by an arrow. The shading of the off-diagonal squares in the heatmap is inversely proportional to the expected position error of the amino acids.

Supplementary Table 1: Sample information for the testis samples included in this study

| Sample ID | Mouse genotype | Mouse strain | Full brothers | Date sampled | Accession number |
|---|---|---|---|---|---|
| 223616 | $+/t^{\mathrm{wIll}}$ | ILL | yes | 06.03.2023 | TBD |
| 223617 | $+/t^{\mathrm{wIll}}$ | ILL | yes | 06.03.2023 | TBD |
| 223618 | $+/+$ | ILL | yes | 06.03.2023 | TBD |
| 272100 | $+/+$ | ILL | no | 07.11.2023 | TBD |

Supplementary Table 2: Gene names, log2 fold change in expression between compared groups, the percent of nuclei expressing the gene in each compared group, the FDR-corrected p-value, the cell type and compared groups and the Ensembl gene IDs are shown for each gene that was differentially expressed in any of the cell types and comparisons in our study.

| Gene | LogFC | Pct1 | Pct2 | PvalAdj | Celltype | Grp1 | Grp2 | EnsemblID |
|---|---|---|---|---|---|---|---|---|
| 4930578E11Rik | -1.62 | 0.324 | 0.585 | 2.93e-09 | Spermatocytes 1 | $+/t$ | $+/+$ | ENSMUSG00000117618 |
| 9530077C14Rik | 0.78 | 0.391 | 0.219 | 3.57e-04 | Elongating SDs | $t^{+/t}$ | $+^{+/+}$ | ENSMUSG00000120262 |
| Actr1b | 0.83 | 0.323 | 0.207 | 3.22e-04 | Elongating SDs | $t^{+/t}$ | $+^{+/+}$ | ENSMUSG00000037351 |
| Agpat2 | 0.66 | 0.577 | 0.392 | 1.10e-02 | Elongating SDs | $+^{+/t}$ | $+^{+/+}$ | ENSMUSG00000026922 |
| Aldh1a1 | 1.23 | 0.601 | 0.306 | 1.32e-10 | Sertoli cells | $+/t$ | $+/+$ | ENSMUSG00000053279 |
| Ankrd17 | 0.8 | 0.298 | 0.184 | 3.52e-03 | Elongating SDs | $+^{+/t}$ | $+^{+/+}$ | ENSMUSG00000055204 |
| Arf5 | 0.69 | 0.295 | 0.178 | 3.08e-02 | Elongating SDs | $t^{+/t}$ | $+^{+/+}$ | ENSMUSG00000020440 |
| Calm3 | 0.98 | 0.386 | 0.203 | 3.25e-03 | Spermatocytes 1 | $+/t$ | $+/+$ | ENSMUSG00000019370 |
| Ccdc183 | 1.15 | 0.699 | 0.457 | 9.76e-04 | Early round SDs | $t^{+/t}$ | $+^{+/+}$ | ENSMUSG00000026940 |
| Cox8c | 0.93 | 0.489 | 0.317 | 1.97e-02 | Spermatocytes 1 | $+/t$ | $+/+$ | ENSMUSG00000043319 |
| Cpne5 | -4.26 | 0.021 | 0.256 | 7.42e-32 | Elongating SDs | $t^{+/t}$ | $+^{+/t}$ | ENSMUSG00000024008 |
| Cpne5 | -4.42 | 0.045 | 0.42 | 1.87e-18 | Condensing SDs | $t^{+/t}$ | $+^{+/t}$ | ENSMUSG00000024008 |
| Cyp4f40 | 1.44 | 0.25 | 0.111 | 3.84e-08 | Elongating SDs | $t^{+/t}$ | $+^{+/+}$ | ENSMUSG00000090700 |
| Ddx5 | 0.92 | 0.912 | 0.709 | 5.22e-03 | Elongating SDs | $t^{+/t}$ | $+^{+/+}$ | ENSMUSG00000020719 |
| Dnah8 | -0.85 | 0.877 | 0.916 | 2.47e-06 | Spermatocytes 1 | $+/t$ | $+/+$ | ENSMUSG00000033826 |
| Dynlt1c | 2.54 | 0.791 | 0.357 | 1.94e-06 | Spermatocytes 2 | $+/t$ | $+/+$ | ENSMUSG00000000579 |
| Dynlt1c | 1.12 | 0.431 | 0.3 | 2.90e-02 | Early round SDs | $t^{+/t}$ | $+^{+/+}$ | ENSMUSG00000000579 |
| Dynlt1c | 1.93 | 0.682 | 0.344 | 2.33e-22 | Spermatocytes 1 | $+/t$ | $+/+$ | ENSMUSG00000000579 |
| Dynlt1c | 1.67 | 0.502 | 0.3 | 1.11e-15 | Early round SDs | $+^{+/t}$ | $+^{+/+}$ | ENSMUSG00000000579 |
| Dynlt2a1 | -4.92 | 0.025 | 0.306 | 1.43e-15 | Sertoli cells | $+/t$ | $+/+$ | ENSMUSG00000079707 |
| Dynlt2a1 | -3.21 | 0.137 | 0.48 | 3.19e-44 | Early round SDs | $+^{+/t}$ | $+^{+/+}$ | ENSMUSG00000079707 |
| Dynlt2a1 | -4.46 | 0.042 | 0.386 | 1.95e-08 | Leydig cells | $+/t$ | $+/+$ | ENSMUSG00000079707 |
| Dynlt2a1 | -3.79 | 0.041 | 0.321 | 4.01e-13 | Late round SDs | $+^{+/t}$ | $+^{+/+}$ | ENSMUSG00000079707 |
| Dynlt2a1 | -1.75 | 0.367 | 0.688 | 3.04e-26 | Spermatocytes 2 | $+/t$ | $+/+$ | ENSMUSG00000079707 |
| Dynlt2a1 | -2.37 | 0.187 | 0.38 | 1.31e-43 | Spermatocytes 1 | $+/t$ | $+/+$ | ENSMUSG00000079707 |
| Dynlt2a1 | -5.35 | 0.025 | 0.321 | 1.16e-13 | Late round SDs | $t^{+/t}$ | $+^{+/+}$ | ENSMUSG00000079707 |
| Dynlt2a1 | -3.15 | 0.118 | 0.48 | 1.64e-38 | Early round SDs | $t^{+/t}$ | $+^{+/+}$ | ENSMUSG00000079707 |
| Dynlt2a1 | -3.07 | 0.091 | 0.357 | 3.62e-10 | Muscle cells | $+/t$ | $+/+$ | ENSMUSG00000079707 |
| Dynlt2a2 | -1.87 | 0.724 | 0.925 | 2.78e-17 | Spermatocytes 1 | $+/t$ | $+/+$ | ENSMUSG00000079710 |
| Dynlt2a2 | -2.41 | 0.073 | 0.336 | 3.33e-08 | Early round SDs | $t^{+/t}$ | $+^{+/t}$ | ENSMUSG00000079710 |
| Dynlt2a2 | -3.31 | 0.073 | 0.51 | 1.54e-08 | Early round SDs | $t^{+/t}$ | $+^{+/+}$ | ENSMUSG00000079710 |
| Dynlt2a2 | -2.11 | 0.117 | 0.418 | 3.19e-05 | Late round SDs | $t^{+/t}$ | $+^{+/t}$ | ENSMUSG00000079710 |
| E430024P14Rik | -2.16 | 0.226 | 0.752 | 2.72e-18 | Condensing SDs | $t^{+/t}$ | $+^{+/t}$ | ENSMUSG00000090069 |
| E430024P14Rik | -2.45 | 0.226 | 0.69 | 3.06e-06 | Condensing SDs | $t^{+/t}$ | $+^{+/+}$ | ENSMUSG00000090069 |
| E430024P14Rik | -3.32 | 0.058 | 0.428 | 3.52e-18 | Elongating SDs | $t^{+/t}$ | $+^{+/+}$ | ENSMUSG00000090069 |
| E430024P14Rik | -2.93 | 0.058 | 0.364 | 1.76e-32 | Elongating SDs | $t^{+/t}$ | $+^{+/t}$ | ENSMUSG00000090069 |
| Eif1b | 1.93 | 0.304 | 0.093 | 1.57e-06 | Elongating SDs | $+^{+/t}$ | $+^{+/+}$ | ENSMUSG00000006941 |
| Exosc2 | 0.92 | 0.656 | 0.387 | 2.07e-03 | Elongating SDs | $+^{+/t}$ | $+^{+/+}$ | ENSMUSG00000039356 |

| | | | | | | | | |
|---|---|---|---|---|---|---|---|---|
| Fbl | -3.45 | 0.058 | 0.541 | 3.63e-13 | Late round SDs | $t^{+/t}$ | $+^{+/t}$ | ENSMUSG00000046865 |
| Fbl | -2.84 | 0.065 | 0.401 | 5.58e-08 | Condensing SDs | $t^{+/t}$ | $+^{+/t}$ | ENSMUSG00000046865 |
| Fbl | -3.25 | 0.077 | 0.598 | 1.00e-72 | Elongating SDs | $t^{+/t}$ | $+^{+/t}$ | ENSMUSG00000046865 |
| Fbl | -3.61 | 0.058 | 0.535 | 1.15e-05 | Late round SDs | $t^{+/t}$ | $+^{+/+}$ | ENSMUSG00000046865 |
| Fbl | -3.28 | 0.077 | 0.563 | 5.67e-32 | Elongating SDs | $t^{+/t}$ | $+^{+/+}$ | ENSMUSG00000046865 |
| Fbxo44 | 1.17 | 0.402 | 0.207 | 4.33e-03 | Sertoli cells | $+/t$ | $+/+$ | ENSMUSG00000029001 |
| Ggt7 | 0.77 | 0.513 | 0.33 | 9.92e-04 | Elongating SDs | $+^{+/t}$ | $+^{+/+}$ | ENSMUSG00000027603 |
| Ggt7 | 0.69 | 0.527 | 0.33 | 2.49e-06 | Elongating SDs | $t^{+/t}$ | $+^{+/+}$ | ENSMUSG00000027603 |
| Gm10505 | -1.72 | 0.658 | 0.864 | 6.33e-13 | Spermatocytes 1 | $+/t$ | $+/+$ | ENSMUSG00000073430 |
| Gm13977 | 0.74 | 0.305 | 0.178 | 1.83e-05 | Elongating SDs | $t^{+/t}$ | $+^{+/+}$ | ENSMUSG00000086735 |
| Gm28940 | 0.97 | 0.338 | 0.181 | 3.90e-03 | Elongating SDs | $+^{+/t}$ | $+^{+/+}$ | ENSMUSG00000100786 |
| Gm29534 | 0.85 | 0.282 | 0.168 | 2.75e-02 | Elongating SDs | $+^{+/t}$ | $+^{+/+}$ | ENSMUSG00000101647 |
| Gm3182 | -0.67 | 0.705 | 0.728 | 8.10e-03 | Early round SDs | $+^{+/t}$ | $+^{+/+}$ | ENSMUSG00000091148 |
| Gm3182 | -0.71 | 0.646 | 0.728 | 3.15e-02 | Early round SDs | $t^{+/t}$ | $+^{+/+}$ | ENSMUSG00000091148 |
| Gm35256 | 1.16 | 0.787 | 0.441 | 2.83e-04 | Late round SDs | $+^{+/t}$ | $+^{+/+}$ | ENSMUSG00000110554 |
| Gm35256 | 1.04 | 0.476 | 0.281 | 1.09e-02 | Early round SDs | $t^{+/t}$ | $+^{+/+}$ | ENSMUSG00000110554 |
| Gm35256 | 0.95 | 0.49 | 0.29 | 2.54e-08 | Elongating SDs | $t^{+/t}$ | $+^{+/+}$ | ENSMUSG00000110554 |
| Gm35256 | 1.04 | 0.541 | 0.29 | 4.52e-08 | Elongating SDs | $+^{+/t}$ | $+^{+/+}$ | ENSMUSG00000110554 |
| Gm44715 | 1.31 | 0.991 | 0.773 | 1.03e-37 | Sertoli cells | $+/t$ | $+/+$ | ENSMUSG00000109196 |
| Gm49441 | 1.77 | 0.969 | 0.657 | 3.46e-06 | Leydig cells | $+/t$ | $+/+$ | ENSMUSG00000115988 |
| Gm55407 | 1.17 | 0.299 | 0.133 | 1.21e-02 | Elongating SDs | $t^{+/t}$ | $+^{+/+}$ | ENSMUSG00002075328 |
| Gm55407 | 1.24 | 0.318 | 0.133 | 4.04e-02 | Elongating SDs | $+^{+/t}$ | $+^{+/+}$ | ENSMUSG00002075328 |
| Gm6566 | 0.68 | 0.499 | 0.324 | 6.77e-03 | Elongating SDs | $+^{+/t}$ | $+^{+/+}$ | ENSMUSG00000113918 |
| Gm9937 | -1.8 | 0.103 | 0.305 | 1.13e-02 | Elongating SDs | $t^{+/t}$ | $+^{+/+}$ | ENSMUSG00000118119 |
| Gstm7 | 2.75 | 0.291 | 0.041 | 2.57e-05 | Sertoli cells | $+/t$ | $+/+$ | ENSMUSG00000004035 |
| H2-Ab1 | -1.28 | 0.12 | 0.274 | 2.97e-05 | Elongating SDs | $t^{+/t}$ | $+^{+/t}$ | ENSMUSG00000073421 |
| H2-Ab1 | -1.75 | 0.12 | 0.321 | 3.93e-04 | Elongating SDs | $t^{+/t}$ | $+^{+/+}$ | ENSMUSG00000073421 |
| Inpp4a | 1.56 | 0.6 | 0.391 | 3.58e-02 | Condensing SDs | $t^{+/t}$ | $+^{+/+}$ | ENSMUSG00000026113 |
| Inpp4a | 1.59 | 0.675 | 0.391 | 4.18e-05 | Condensing SDs | $+^{+/t}$ | $+^{+/+}$ | ENSMUSG00000026113 |
| Ldhc | 3.75 | 0.279 | 0.029 | 1.06e-10 | Late round SDs | $+^{+/t}$ | $+^{+/+}$ | ENSMUSG00000030851 |
| Ldhc | 3.26 | 0.439 | 0.062 | 6.35e-05 | Spermatogonia | $+/t$ | $+/+$ | ENSMUSG00000030851 |
| Ldhc | 2.21 | 0.354 | 0.069 | 4.09e-30 | Early round SDs | $t^{+/t}$ | $+^{+/+}$ | ENSMUSG00000030851 |
| Ldhc | 2.83 | 0.344 | 0.064 | 1.23e-08 | Condensing SDs | $+^{+/t}$ | $+^{+/+}$ | ENSMUSG00000030851 |
| Ldhc | 2.29 | 0.297 | 0.064 | 7.38e-07 | Condensing SDs | $t^{+/t}$ | $+^{+/+}$ | ENSMUSG00000030851 |
| Ldhc | 2.67 | 0.684 | 0.161 | 2.86e-14 | Spermatocytes 2 | $+/t$ | $+/+$ | ENSMUSG00000030851 |
| Ldhc | 3.19 | 0.396 | 0.073 | 1.09e-44 | Spermatocytes 1 | $+/t$ | $+/+$ | ENSMUSG00000030851 |
| Ldhc | 2.81 | 0.461 | 0.069 | 1.90e-46 | Early round SDs | $+^{+/t}$ | $+^{+/+}$ | ENSMUSG00000030851 |
| Ldhc | 2.79 | 0.364 | 0.054 | 4.24e-11 | Muscle cells | $+/t$ | $+/+$ | ENSMUSG00000030851 |
| Ldhc | 3.07 | 0.267 | 0.029 | 2.30e-06 | Late round SDs | $t^{+/t}$ | $+^{+/+}$ | ENSMUSG00000030851 |
| Lypla2 | 1.69 | 0.27 | 0.103 | 4.56e-03 | Late round SDs | $+^{+/t}$ | $+^{+/+}$ | ENSMUSG00000028670 |
| Mas1 | -5.88 | 0.004 | 0.382 | 3.56e-27 | Early round SDs | $t^{+/t}$ | $+^{+/t}$ | ENSMUSG00000068037 |
| Mas1 | -5.44 | 0.004 | 0.283 | 1.09e-13 | Early round SDs | $t^{+/t}$ | $+^{+/+}$ | ENSMUSG00000068037 |
| mt-Co2 | -0.63 | 0.273 | 0.473 | 1.17e-13 | Muscle cells | $+/t$ | $+/+$ | ENSMUSG00000064354 |
| mt-Co2 | -0.6 | 0.404 | 0.688 | 1.45e-04 | Spermatogonia | $+/t$ | $+/+$ | ENSMUSG00000064354 |
| mt-Co2 | -1.31 | 0.157 | 0.257 | 3.98e-04 | Leydig cells | $+/t$ | $+/+$ | ENSMUSG00000064354 |
| mt-Co2 | -0.68 | 0.39 | 0.702 | 7.57e-39 | Sertoli cells | $+/t$ | $+/+$ | ENSMUSG00000064354 |
| mt-Co3 | 3.72 | 0.271 | 0.056 | 8.10e-25 | Elongating SDs | $t^{+/t}$ | $+^{+/+}$ | ENSMUSG00000064358 |
| mt-Co3 | 1.08 | 0.641 | 0.429 | 8.72e-03 | Muscle cells | $+/t$ | $+/+$ | ENSMUSG00000064358 |
| mt-Co3 | 3.58 | 0.286 | 0.056 | 8.59e-20 | Elongating SDs | $+^{+/t}$ | $+^{+/+}$ | ENSMUSG00000064358 |
| mt-Cytb | 1.2 | 0.296 | 0.106 | 1.01e-02 | Spermatocytes 1 | $+/t$ | $+/+$ | ENSMUSG00000064370 |
| mt-Rnr2 | 1.19 | 0.298 | 0.117 | 1.04e-03 | Spermatocytes 1 | $+/t$ | $+/+$ | ENSMUSG00000064339 |
| mt-Rnr2 | 4.24 | 0.361 | 0.047 | 5.10e-04 | Late round SDs | $+^{+/t}$ | $+^{+/+}$ | ENSMUSG00000064339 |
| Oard1 | 1.47 | 0.29 | 0.115 | 2.57e-05 | Elongating SDs | $+^{+/t}$ | $+^{+/+}$ | ENSMUSG00000040771 |
| Or10h28 | 0.98 | 0.423 | 0.242 | 2.28e-06 | Elongating SDs | $+^{+/t}$ | $+^{+/+}$ | ENSMUSG00000054666 |
| Or5b106 | 3.29 | 0.955 | 0.317 | 3.21e-41 | Condensing SDs | $t^{+/t}$ | $+^{+/+}$ | ENSMUSG00000057503 |

| | | | | | | | | |
|---|---|---|---|---|---|---|---|---|
| Or5b106 | 2.36 | 0.898 | 0.489 | 1.77e-51 | Early round SDs | $t^{+/t}$ | $+^{+/+}$ | ENSMUSG00000057503 |
| Or5b106 | 3.11 | 0.981 | 0.459 | 2.11e-175 | Elongating SDs | $t^{+/t}$ | $+^{+/+}$ | ENSMUSG00000057503 |
| Or5b106 | 2.79 | 0.981 | 0.553 | 3.57e-202 | Elongating SDs | $t^{+/t}$ | $+^{+/t}$ | ENSMUSG00000057503 |
| Or5b106 | 2.3 | 0.898 | 0.548 | 1.18e-47 | Early round SDs | $t^{+/t}$ | $+^{+/t}$ | ENSMUSG00000057503 |
| Or5b106 | 3.16 | 0.992 | 0.418 | 5.38e-53 | Late round SDs | $t^{+/t}$ | $+^{+/+}$ | ENSMUSG00000057503 |
| Or5b106 | 2.72 | 0.992 | 0.525 | 8.91e-50 | Late round SDs | $t^{+/t}$ | $+^{+/t}$ | ENSMUSG00000057503 |
| Or5b106 | 2.38 | 0.955 | 0.471 | 2.74e-40 | Condensing SDs | $t^{+/t}$ | $+^{+/t}$ | ENSMUSG00000057503 |
| Or8b3 | -1.22 | 0.135 | 0.29 | 1.22e-02 | Spermatocytes 1 | $+/t$ | $+/+$ | ENSMUSG00000049098 |
| Phf1 | 2.93 | 0.297 | 0.05 | 8.37e-08 | Condensing SDs | $t^{+/t}$ | $+^{+/+}$ | ENSMUSG00000024193 |
| Pkig | 0.85 | 0.489 | 0.303 | 3.90e-05 | Spermatocytes 1 | $+/t$ | $+/+$ | ENSMUSG00000035268 |
| Ppp1r11 | 2.93 | 0.98 | 0.449 | 5.63e-67 | Early round SDs | $t^{+/t}$ | $+^{+/+}$ | ENSMUSG00000036398 |
| Ppp1r11 | 1.99 | 0.98 | 0.676 | 7.83e-63 | Early round SDs | $t^{+/t}$ | $+^{+/t}$ | ENSMUSG00000036398 |
| Ppp1r11 | 4.21 | 0.787 | 0.121 | 9.18e-14 | Condensing SDs | $t^{+/t}$ | $+^{+/+}$ | ENSMUSG00000036398 |
| Ppp1r11 | 2.42 | 0.958 | 0.494 | 9.30e-23 | Late round SDs | $t^{+/t}$ | $+^{+/+}$ | ENSMUSG00000036398 |
| Ppp1r11 | 2.03 | 0.787 | 0.408 | 9.97e-17 | Condensing SDs | $t^{+/t}$ | $+^{+/t}$ | ENSMUSG00000036398 |
| Ppp1r11 | 2.12 | 0.572 | 0.196 | 6.55e-31 | Elongating SDs | $t^{+/t}$ | $+^{+/+}$ | ENSMUSG00000036398 |
| Ppp1r11 | 1.66 | 0.427 | 0.167 | 4.19e-16 | Spermatocytes 1 | $+/t$ | $+/+$ | ENSMUSG00000036398 |
| Ppp1r11 | 1.36 | 0.572 | 0.298 | 1.06e-19 | Elongating SDs | $t^{+/t}$ | $+^{+/t}$ | ENSMUSG00000036398 |
| Ppp1r11 | 1.81 | 0.958 | 0.672 | 9.56e-23 | Late round SDs | $t^{+/t}$ | $+^{+/t}$ | ENSMUSG00000036398 |
| Ppp1r2 | 1.8 | 0.408 | 0.131 | 3.42e-02 | Late round SDs | $t^{+/t}$ | $+^{+/t}$ | ENSMUSG00000047714 |
| Ppp1r2 | 1.46 | 0.39 | 0.137 | 5.84e-05 | Early round SDs | $t^{+/t}$ | $+^{+/t}$ | ENSMUSG00000047714 |
| Ptma | 1.03 | 0.264 | 0.121 | 2.71e-04 | Spermatocytes 1 | $+/t$ | $+/+$ | ENSMUSG00000026238 |
| Ropn1l | 0.85 | 0.567 | 0.305 | 4.97e-02 | Spermatocytes 1 | $+/t$ | $+/+$ | ENSMUSG00000022236 |
| Rpl13 | 1.06 | 0.441 | 0.228 | 4.27e-04 | Spermatocytes 1 | $+/t$ | $+/+$ | ENSMUSG00000000740 |
| Rpl23a | 1.31 | 0.434 | 0.162 | 3.38e-02 | Late round SDs | $+^{+/t}$ | $+^{+/+}$ | ENSMUSG00000058546 |
| Rsph3a | 1.84 | 0.299 | 0.097 | 2.67e-02 | Early round SDs | $+^{+/t}$ | $+^{+/+}$ | ENSMUSG00000073471 |
| Rsph3a | 2.36 | 0.495 | 0.119 | 1.35e-22 | Spermatocytes 1 | $+/t$ | $+/+$ | ENSMUSG00000073471 |
| Sipa1l3 | 0.78 | 0.35 | 0.191 | 4.58e-02 | Elongating SDs | $+^{+/t}$ | $+^{+/+}$ | ENSMUSG00000030583 |
| Smco2 | -1.43 | 0.121 | 0.282 | 1.39e-04 | Spermatocytes 1 | $+/t$ | $+/+$ | ENSMUSG00000030292 |
| Smim24 | 1.3 | 0.402 | 0.18 | 1.41e-03 | Spermatocytes 1 | $+/t$ | $+/+$ | ENSMUSG00000078439 |
| Smok-Tcr | 2.3 | 0.297 | 0.076 | 9.55e-03 | Condensing SDs | $t^{+/t}$ | $+^{+/t}$ | ENSMUSG00000073458 |
| Smok4a | -2 | 0.097 | 0.338 | 1.13e-02 | Condensing SDs | $t^{+/t}$ | $+^{+/t}$ | ENSMUSG00000121414 |
| Spata2 | -4.53 | 0.021 | 0.391 | 1.90e-29 | Elongating SDs | $t^{+/t}$ | $+^{+/+}$ | ENSMUSG00000047030 |
| Spata2 | -4.51 | 0.021 | 0.41 | 1.57e-56 | Elongating SDs | $t^{+/t}$ | $+^{+/t}$ | ENSMUSG00000047030 |
| Spata2 | -2.88 | 0.058 | 0.382 | 1.31e-07 | Condensing SDs | $t^{+/t}$ | $+^{+/t}$ | ENSMUSG00000047030 |
| Synj2bp | 1.12 | 0.366 | 0.186 | 4.88e-15 | Elongating SDs | $+^{+/t}$ | $+^{+/+}$ | ENSMUSG00000090935 |
| Synj2bp | 0.59 | 0.436 | 0.292 | 8.20e-04 | Early round SDs | $+^{+/t}$ | $+^{+/+}$ | ENSMUSG00000090935 |
| Synj2bp | 0.64 | 0.451 | 0.292 | 6.01e-07 | Early round SDs | $t^{+/t}$ | $+^{+/+}$ | ENSMUSG00000090935 |
| Synj2bp | 1.01 | 0.346 | 0.186 | 9.67e-17 | Elongating SDs | $t^{+/t}$ | $+^{+/+}$ | ENSMUSG00000090935 |
| t-Ppp1cb | 5.88 | 0.386 | 0 | 3.59e-04 | Spermatocytes 2 | $+/t$ | $+/+$ | ENSMUSG00000014956 |
| Vmn2r112 | 0.7 | 0.344 | 0.207 | 2.31e-02 | Elongating SDs | $t^{+/t}$ | $+^{+/+}$ | ENSMUSG00000094921 |
| Zwilch | 2.6 | 0.29 | 0.05 | 1.43e-09 | Elongating SDs | $t^{+/t}$ | $+^{+/+}$ | ENSMUSG00000032400 |
| Zwilch | 2.78 | 0.314 | 0.05 | 5.41e-12 | Elongating SDs | $+^{+/t}$ | $+^{+/+}$ | ENSMUSG00000032400 |

Supplementary Table 3: Enriched terms and the list of proteins in each category. Observed - Number of genes observed in category. Background - Total number of genes in the category.

| Category | Term ID | Term description | Observed | Background | Strength | Signal | FDR | Observed proteins |
|---|---|---|---|---|---|---|---|---|
| GO Component | GO:0070069 | Cytochrome complex | 3 | 37 | 1.85 | 0.68 | 0.02270 | Cox8c,mt-Co2,mt-Cytb |
| GO Component | GO:0031514 | Motile cilium | 5 | 362 | 1.08 | 0.47 | 0.04660 | Ldhc,Calm3,Rsph3a,Ropn1l,Dnah8 |
| STRING clusters | CL:8158 | Protein phosphatase type 1 complex, and Phosphorylase kinase complex | 3 | 32 | 1.91 | 0.58 | 0.04120 | Ppp1cb,Ppp1r11,Ppp1r2 |
| KEGG | mmu00190 | Oxidative phosphorylation | 3 | 191 | 1.14 | 0.48 | 0.04910 | Cox8c,mt-Co2,mt-Cytb |
| KEGG | mmu04260 | Cardiac muscle contraction | 3 | 95 | 1.44 | 0.52 | 0.04910 | Cox8c,mt-Co2,mt-Cytb |
| KEGG | mmu04814 | Motor proteins | 4 | 265 | 1.12 | 0.47 | 0.04910 | Tcte3,Dynlt1c,Gm3448,Dnah8 |
| KEGG | mmu04932 | Non-alcoholic fatty liver disease | 3 | 165 | 1.20 | 0.49 | 0.04910 | Cox8c,mt-Co2,mt-Cytb |
| KEGG | mmu05010 | Alzheimer disease | 4 | 409 | 0.93 | 0.43 | 0.04910 | Calm3,Cox8c,mt-Co2,mt-Cytb |
| KEGG | mmu05012 | Parkinson disease | 4 | 276 | 1.10 | 0.47 | 0.04910 | Calm3,Cox8c,mt-Co2,mt-Cytb |
| KEGG | mmu05014 | Amyotrophic lateral sclerosis | 4 | 386 | 0.96 | 0.44 | 0.04910 | Cox8c,mt-Co2,mt-Cytb,Dnah8 |
| KEGG | mmu05016 | Huntington disease | 4 | 313 | 1.05 | 0.46 | 0.04910 | Cox8c,mt-Co2,mt-Cytb,Dnah8 |
| KEGG | mmu05022 | Pathways of neurodegeneration - multiple diseases | 5 | 490 | 0.95 | 0.44 | 0.04910 | Calm3,Cox8c,mt-Co2,mt-Cytb,Dnah8 |
| KEGG | mmu05171 | Coronavirus disease - COVID-19 | 4 | 280 | 1.10 | 0.47 | 0.04910 | Rpl13,Ptma,Mas1,Rpl23a |
| KEGG | mmu05415 | Diabetic cardiomyopathy | 4 | 227 | 1.19 | 0.48 | 0.04910 | Ppp1cb,Cox8c,mt-Co2,mt-Cytb |
| TISSUES | BTO:0001363 | Testis | 9 | 1287 | 0.79 | 0.54 | 0.00700 | Ldhc,Tcte3,Ppp1r11,Cox8c,mt-Cytb,Dynlt1c,Smim24,Ropn1l,Dnah8 |
| TISSUES | BTO:0000083 | Female reproductive system | 10 | 2449 | 0.55 | 0.35 | 0.03260 | Ldhc,Calm3,Tcte3,Ppp1r11,Cox8c,mt-Cytb,Dynlt1c,Smim24,Ropn1l,Dnah8 |
| COMPARTMENTS | GOCC:0031514 | Motile cilium | 6 | 272 | 1.28 | 0.99 | 0.00097 | Ldhc,Calm3,Tcte3,Rsph3a,Ropn1l,Dnah8 |
| COMPARTMENTS | GOCC:0097729 | 9+2 motile cilium | 5 | 234 | 1.27 | 0.78 | 0.00560 | Calm3,Tcte3,Rsph3a,Ropn1l,Dnah8 |
| COMPARTMENTS | GOCC:0030286 | Dynein complex | 3 | 52 | 1.70 | 0.70 | 0.01860 | Tcte3,Dynlt1c,Dnah8 |
| COMPARTMENTS | GOCC:0036126 | Sperm flagellum | 4 | 218 | 1.20 | 0.53 | 0.03710 | Calm3,Tcte3,Ropn1l,Dnah8 |
| COMPARTMENTS | GOCC:0001534 | Radial spoke | 2 | 15 | 2.07 | 0.56 | 0.04720 | Rsph3a,Ropn1l |
| Pfam | PF03645 | Tctex-1 family | 3 | 11 | 2.38 | 1.27 | 0.00100 | Tcte3,Dynlt1c,Gm3448 |
